# Sporadic ERK pulses drive non-genetic resistance in drug-adapted BRAF^V600E^ melanoma cells

**DOI:** 10.1101/762294

**Authors:** Luca Gerosa, Christopher Chidley, Fabian Froehlich, Gabriela Sanchez, Sang Kyun Lim, Jeremy Muhlich, Jia-Yun Chen, Gregory J. Baker, Denis Schapiro, Tujin Shi, Lian Yi, Carrie D. Nicora, Allison Claas, Douglas A. Lauffenburger, Wei-Jun Qian, H. Steven Wiley, Peter K. Sorger

## Abstract

Anti-cancer drugs commonly target signal transduction proteins activated by mutation. In patients with BRAF^V600E^ melanoma, small molecule RAF and MEK kinase inhibitors cause dramatic but often transient tumor regression. Emerging evidence suggests that cancer cells adapting by non-genetic mechanisms constitute a reservoir for the development of drug-resistant tumors. Here, we show that few hours after exposure to RAF/MEK inhibitors, BRAF^V600E^ melanomas undergo adaptive changes involving disruption of negative feedback and sporadic pulsatile reactivation of the MAPK pathway, so that MAPK activity is transiently high enough in some cells to drive proliferation. Quantitative proteomics and computational modeling show that pulsatile MAPK reactivation is possible due to the co-existence in cells of two MAPK cascades: one driven by BRAF^V600E^ that is drug-sensitive and a second driven by receptors that is drug-resistant. Paradoxically, this may account both for the frequent emergence of drug resistance and for the tolerability of RAF/MEK therapy in patients.

## INTRODUCTION

Mammalian cells transduce extracellular mitogenic signals via signaling cascades comprising transmembrane receptors, intracellular kinases and second messengers. Oncogenic mutations are common in these proteins and “addiction” of tumor cells to mutated signaling pathways is a rationale for targeted anti-cancer therapy (Weinstein, 2002)(Weinstein and Joe, 2006). Targeted therapies have been developed for multiple oncogenes and types of cancer, including the ∼50% of melanomas with mutated BRAF (canonically BRAF^V600E^, the focus of this paper). Oncogenic RAF mutations constitutively activate the mitogenic MAPK signaling cascade comprising RAF, MEK and ERK kinases. In BRAF^V600E^ melanoma patients the use of FDA-approved small molecule inhibitors of RAF (vemurafenib, Zelboraf® or dabrafenib, Tafinlar®) and MEK (cobimetinib, Cotellic®, or trametinib, Mekinist®) effectively blocks oncogenic signaling. Therapeutic response to RAF/MEK therapy is rapid but often transitory due to the emergence of drug resistance (Groenendijk and Bernards, 2014). Many patients eventually acquire genetic mutations in MAPK pathway proteins (e.g. in RAS or MEK) and relapse because of ERK reactivation (Johnson et al., 2015). A better understanding of mechanisms of drug resistance and their blockade are widely regarded as keys to achieving durable responses to targeted therapies.

While the focus on drug resistance is often heritable genetic mutations, emerging evidence suggests that the early adaptive changes by which melanoma cells respond to targeted drugs may be crucial for their survival, the retention of minimal residual disease, and the eventual acquisition of drug-resistant cancers (Pazarentzos and Bivona, 2015). In melanoma cell lines, adaptive resistance is observed soon after drug exposure; it is reversible following a drug holiday and resembles the non-mutational drug resistance of slow dividing “persister” cells (Ramirez et al., 2016)(Fallahi-Sichani et al., 2017)(Shaffer et al., 2017). Multiple studies across cancer types and targeted therapies suggest that adaptive drug resistance is enabled by mechanisms of signaling homeostasis (Carver et al., 2011)(Niederst and Engelman, 2013)(Goel et al., 2016). The role of negative feedback in homeostasis and drug resistance is particularly well established in the case of BRAF^V600E^ cancers. When BRAF^V600E^ signaling is inhibited by drugs, synthesis of phosphatases and other negative regulators of MAPK signaling falls. This makes cells more sensitive to reactivation of the pathway, for example, by growth factors in the tumor microenvironment (Lito et al., 2012)(Chandarlapaty, 2012)(Prahallad et al., 2012). Research into drug adaption in BRAF^V60OE^ melanomas therefore affords an opportunity to investigate the basic biology of MAPK homeostatis (Kolch et al., 2015)(Rauch et al., 2016), determine how targeted drugs rewire signaling networks, and potentially improve the treatment of metastatic melanoma (Chapman et al., 2011)(Flaherty et al., 2012).

Despite elegant studies by Rosen and others (Lito et al., 2012)(Sun et al., 2014), the role played by early adaptive reactivation of MAPK signaling in BRAF^V60OE^ melanoma cells remains unclear. Some reports suggest that ERK remains strongly inhibited (Pratilas et al., 2009)(Montero-Conde et al., 2013)(Fallahi-Sichani et al., 2015), whereas others suggest that it quickly rebounds to a low extent (Lito et al., 2012). The extracellular environment, including autocrine or paracrine growth factors, has been shown to play a role in promoting drug resistance (Straussman et al., 2012)(Wilson et al., 2012) but how mitogenic signals are transduced when MAPK is inhibited remains unclear: like many other types of mammalian cells, melanocytes require MAPK activity to divide. Thus, the fundamental mystery of the adapted state is how melanoma cells can grow and divide when MAPK signaling is profoundly inhibited by RAF and MEK inhibitors.

In this paper we investigate the drug-adapted state of BRAF^V600E^ melanoma cells using live and fixed cell imaging combined with proteomics and mathematical modeling. We find that drug-induced network rewiring causes BRAF^V600E^-mutant melanoma cells to experience sporadic ERK pulses of sufficient duration (∼60-90 min) to drive cell division. ERK pulses are locally triggered by autocrine/paracrine factors in the local microenvironment so that ERK activity levels are low on average, but transiently high enough in patches of cells to explain the infrequent division characteristic of persister cells. A dynamic computational model of MAPK signaling informed by targeted proteomics reveals that pulsatile MAPK reactivation is possible due to the co-existence in cells of two MAPK cascades: one driven by BRAF^V600E^ that is drug-sensitive and a second driven by the microenvironment and transmembrane receptors that is highly resistant to both RAF and MEK inhibitors.

## RESULTS

### BRAF^V600E^ melanoma cells exposed to RAF inhibitors exhibit spontaneous ERK pulsing

When BRAF^V600E^ A375 melanoma cells were treated with a clinically-relevant concentration of vemurafenib for 4 days (1 µM), a minority of cells died and the remainder arrested in G0/G1 (Figure 1A). As previously described, the activity of the MAPK cascade fell 10-fold within 20 min of drug exposure and then rebounded by 2hr to only ∼1-5% of its initial level (Figure 1B)(Lito et al., 2012). Activity remained low concomitant with the emergence of a subpopulation of slowly dividing cells 1-4 days later (∼5% of cells incorporating EdU over a 3h exposition; Figure 1A). Single-cell data acquired over the course of adaptation by immunofluorescence imaging using antibodies against activating sites on ERK1^T202,Y204^ and ERK2^T185,Y187^ (henceforth pERK) showed that rebound involved the appearance of a minority of cells (∼1-2% at 24 hr and ∼5% at 4 days) with high ERK activity rather than a small increase in ERK activity in all cells. In ERK^high^ cells, pERK levels were similar to those in drug-naïve cells (Figure 1B). ERK^high^ cells were eliminated and cell proliferation reduced by addition of saturating concentrations of a MEK inhibitor, showing that both are dependent on MAPK activity (in this experiment cobimetinib was used at 1 µM, >10-fold above the clinically relevant concentration; Figure 1C-D). Moreover, across a more physiological range of MEK inhibitor concentrations, there was a good correlation between the number of ERK^high^ cells and cell cycle progression (Figure 1C). ERK^high^ cells were observed for multiple RAF inhibitors (**Figure S1A**), hours to days after drug exposure, and in five independent A375 subclones, suggesting adaptive rather than genetic modes of ERK activation (**Figure S1B**). In seven other BRAF-mutant melanoma lines 0.1% to 5% of cells were ERK^high^ following exposure to RAF inhibitors and complete suppression of ERK by saturating doses of a MEK inhibitor reduced the number of surviving cells (Figure 1D and **Figure S1C)**. Thus, the small ERK rebound observed in the presence of RAF inhibitors comprises rare cells with high ERK activity and this is associated with fractional proliferation during treatment.

**Figure 1.**
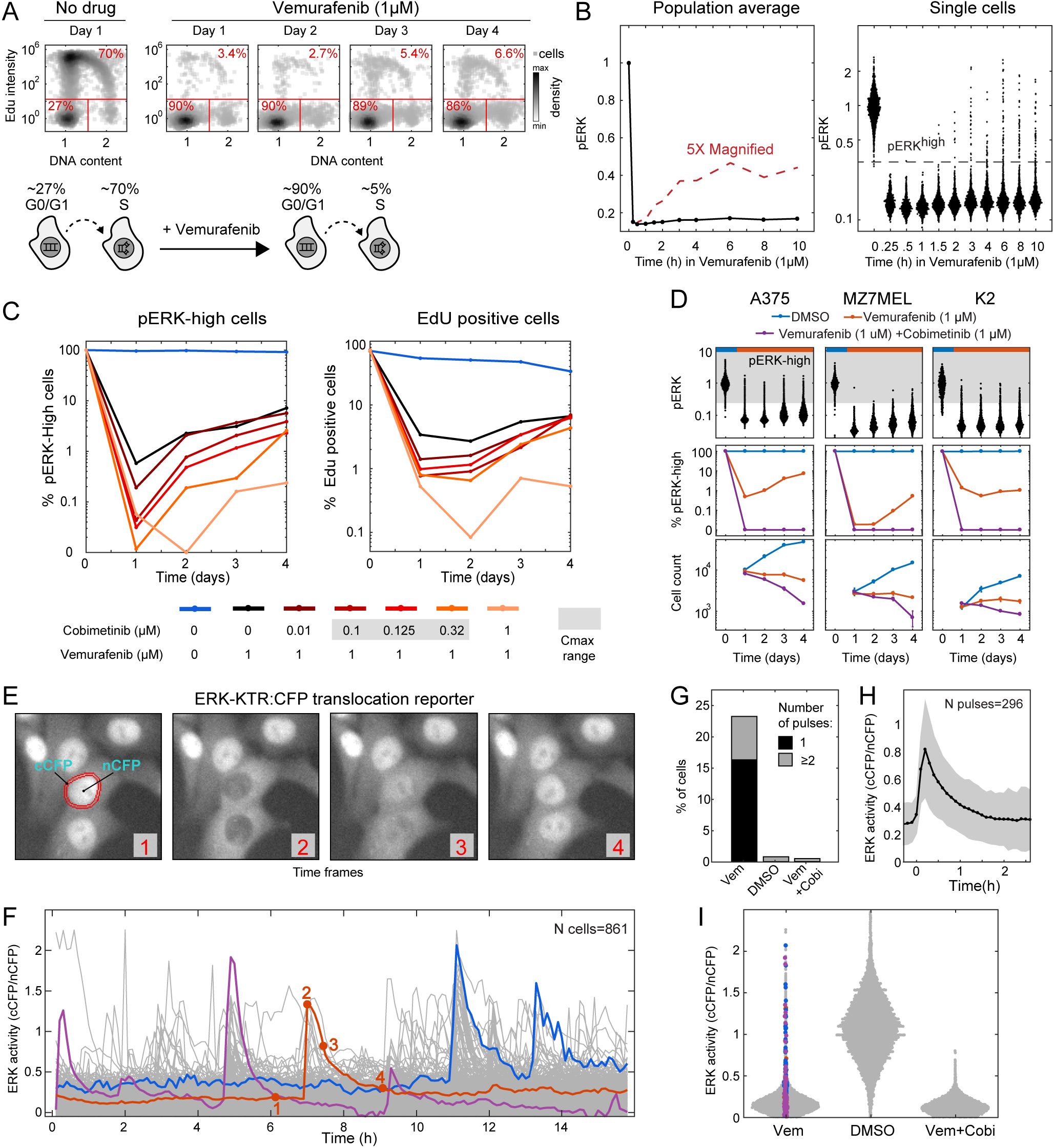
BRAF^V600E^ melanoma cells exposed to RAF inhibitors exhibit spontaneous ERK pulsing. **A)** DNA content and EdU incorporation (for 3 hr) in A375 cells exposed to vemurafenib (1 μM) for 1-4 days; percent of cells in G0/G1 (EdU-negative with DNA<1.5) and EdU-positive cells are shown in red. **B)** Average (right) and single-cell (left) pERK levels as measured by immunofluorescence microscopy in A375 cells in 1 μM vemurafenib. **C)** Percentage of pERK^HIGH^ cells and EdU positive A375 cells treated with vemurafenib (1 μM) or varying cobimetinib concentrations for 4 days. **D)** Single-cell pERK distributions for three BRAF^V600E^ melanoma cell lines treatment with vemurafenib. Percentage of pERK^HIGH^ cells (3^rd^ row) and viable cell counts (4^th^ row) during treatment with vemurafenib with or without 1 μM cobimetinib. **E)** ERK activity in A375 cells quantified by the cytosolic to nuclear ratio of in the CFP channel (cCFP/nCFP) of the ERK-KTR reporter. **F)** ERK activity traces (N=861) from A375 cells treated with vemurafenib (1 uM, 24h) imaged every 6 min for 15 hr. Cells with a single (red), two distinct (purple) and two consecutive ERK pulses (blue) are highlighted. Red numbers correspond to images in panel (E). **G)** Percentage of A375 cells with one or more pulses over a 15 hr period. **H)** Shape of a typical ERK pulse obtained by averaging N=296 live-cell trajectories. Shading denotes one standard deviation around the mean (solid line). **I)** ERK activity distributions of A375 single-cells treated with DMSO, Vemurafenib (1 uM) alone or vemurafenib plus a high dose of cobimetinib (1 uM). Colored dots denote ERK activity levels in the colored traces in panel (F).

To determine the origin of ERK^high^ A375 cells, the ERK activity reporter ERK-KTR:CFP (Regot et al., 2014)(Fallahi-Sichani et al., 2017) was stably introduced using transposons. The reporter translocates from the nucleus to the cytosol when phosphorylated by ERK, providing time-resolved, single cell kinase activity data (Figure 1E). To facilitate tracking, cells co-expressed the histone fusion H2B:YFP. When reporter-expressing cells were exposed to 1 µM vemurafenib for 24 hr and imaged every 6 min over a period of 15 hr (n=861 cells), ERK was inactive most of the time but pulsed on in individual cells at irregular intervals (compare blue, red and purple trajectories; Figure 1F). We observed 134 examples of a single pulse over a 15 hr observation period (∼16% of cells) and 57 examples of multiple pulses (∼7% of cells) (Figure 1G). Individual pulses were found to have a characteristic amplitude and duration: they typically rose to a maximum within 10 min of detection and fell to baseline within 60-90 min (Figure 1H). Maximum ERK activity was similar to that of cells not treated with drug. When assayed by live-cell imaging, ERK was active in ∼1 in 100 vemurafinib-treated A375 cells at any single point in time, consistent with population-average measurements showing ∼1-5% average rebound (Figure 1I). However, over a 15 hr period, as many as ∼25% of cells experienced one or more ERK pulses (Figure 1G). Thus, the RAF-inhibitor adapted state in BRAF^V600E^ melanoma is associated with a dramatic reduction in average MAPK activity but at the level of single cells, spontaneous and irregular ERK pulses are sufficiently frequent to explain ongoing cell division.

### Self-limiting receptor-driven ERK pulses induce expression of cell cycle genes in drug-adapted cells

Because spontaneous ERK pulsing in vemurafinib-adapted cells are infrequent and unpredictable, we sought to recreate them under controlled conditions. An extensive literature exists on the sensitivity of melanoma cells to growth factors in the microenvironment (Straussman et al., 2012)(Wilson et al., 2012). We tested EGF, neuregulin (NRG1), FGF8 and HGF. When added to the medium of normally growing cells, none of these ligands appreciably altered ERK activity, even at supra-physiological doses (Figure 2A). However, cells that had been exposed to vemurafenib for >2hr became highly sensitive to growth factor addition (Figure 2A). Over an ∼10^3^-fold concentration range (spanning approximately physiological to saturating) growth factors caused pERK levels to rise rapidly and transiently, reaching a maximum of 50-150% of their pre-treatment level within 5 to 20 min. and then falling to baseline 45-60 min later (Figure 2B).

**Figure 2.**
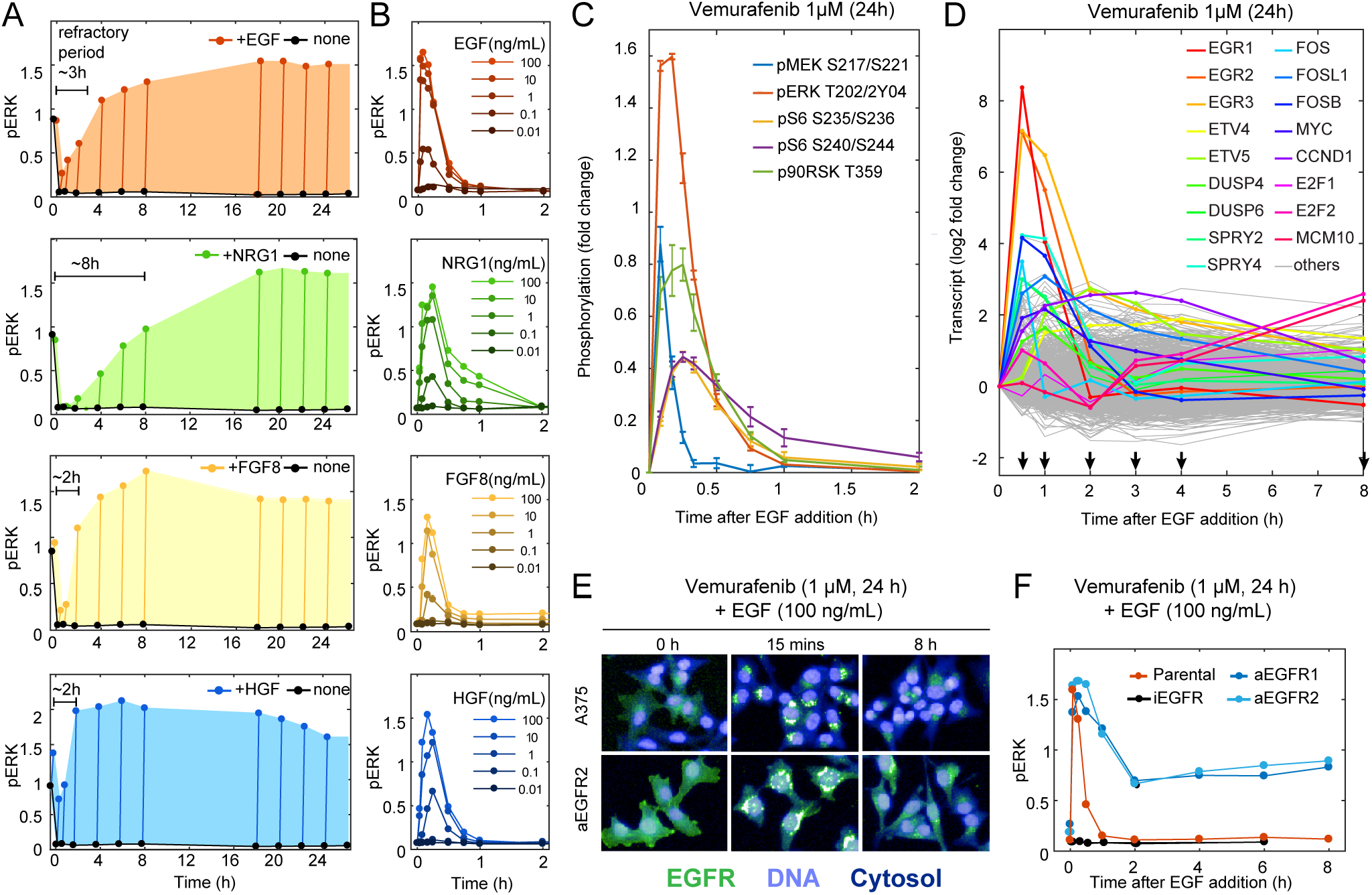
Self-limiting receptor-driven ERK pulses induce expression of cell cycle genes in drug-adapted cells. Cells were exposed to 1 μM vemurafenib for 24 hr unless noted otherwise **A)** pERK levels in cells exposed to one of four growth factors (100 ng/mL) for 15 min at different times after the addition of vemurafenib. Shading interpolates maximal pERK levels. **B)** pERK in cells exposed to different concentrations of one of four growth factors in A375 cells exposed to vemurafenib. **C)** Time course of phosphorylation of various MAPK and downstream signaling proteins on activating sites in cells exposed to vemurafenib and then EGF (100 ng/mL). **D)** Transcript levels in A375 cells exposed to vemurafenib and then EGF (100 ng/mL); data from a next-generation mRNA sequencing involving 4 replicate samples are shown. **E)** EGFR localization by immunofluorescence microscopy in normal and EGFR-overexpressing (A375 aEGFR1) cells exposed to vemurafenib and then to EGF (100 ng/mL) for the times indicated. **F)** pERK levels are shown for parental, A375 aEGFR1 and A375 aEGFR2 CRISPRa over-expressing cell lines and an EGFR CRISPRi down-regulated line and exposed to vemurafenib and then to EGF (100 ng/mL) for the times shown. Measurements are normalized to a DMSO control. In C and D mean and standard deviations from at least 2 replicates are shown.

In adapted cells, exposure to growth factors caused a transient increase in MEK phosphorylation on the activating sites S217/S221 (henceforth pMEK) and the same was true of p90RSK^T359^, pS6^S240/244^ and pS6^S235/236^ (Figure 2C and **Figure S2A**). A brief MAPK pulse generated by ligand addition was sufficient to induce significant expression of multiple mRNA transcripts, including those for immediate-early response genes such as EGR1 and FOSL1 and feedback regulators such as DUSP4/6 and SPRY2/4 (Figure 2D and **Figure S2B**). Immediate early genes were expressed 1-2 hr after ligand addition and early genes such as CCND1 (Cyclin D) and E2F1/2 peaked between 2 and 8 hr. It has previously been shown that cyclin D can drive cell division of A375 cells (Chen et al., 2018) potentially explaining the correlation between ERK pulsing and S phase entry in vemurafenib and cobimetinib treated cells (Figure 1C).

The time scale of ERK activation by exogenous ligand was similar to that of spontaneous ERK pulses and inactivation kinetics were also similar to those of receptor internalization and degradation (Figure 2E, **top panels**). Receptor degradation and internalization are known explanations for transient MAPK signaling when RTKs are present at low levels (Resat et al., 2003), as they are in melanoma cells (Lito et al., 2012). Calibrated ELISA assays of A375 cell extracts showed EGFR, HER2, HER3 and c-MET to be present at ∼200-2000 copies per cell following vemurafenib exposure (**Figure S2C**). On average, we found that RTKs fell ∼3-fold in abundance in the presence of a cognate ligand (**Figure S2C**). To test if receptor abundance truly determines ERK reactivation dynamics, we used CRISPRa and two different RNA guides to over-express EGFR ∼5-8 fold (**Figure S2D, E)**. When these cells were treated with 1 µM vemurafinib for 24 hr followed by exposure to EGF pERK levels rose rapidly and then maintained substantial ERK activity (∼70% of their levels in drug-naïve cells) for at least 24 hr thereafter (Figure 2F). Under these conditions, a fraction of EGFR remained on the plasma membrane (where it is expected to be active) and a fraction moved to endosomes (where degradation and recycling occur; Figure 2E**, bottom panels**). These data strongly suggest that receptor internalization and degradation are involved in the self-limiting nature of ERK pulses in drug-treated BRAF^V600E^ cells, specifically when RTKs are present at low levels.

We conclude that: (i) in normally growing A375 cells MAPK signaling is chronically active, as expected for cells expressing BRAF^V600E^, and is not further activated by growth factor exposure; (ii) cells become highly sensitive to exogenous growth factors only when BRAF^V600E^ activity is blocked for >2hr by inhibitors such as vemurafenib; (iii) MAPK activation by exogenous growth factors is transient and its kinetics and magnitude are similar to those of spontaneous ERK pulses (iv) transient MAPK activation is likely due to low levels of receptors that are internalized and degraded following growth factor binding and (v) transient MAPK pulses are sufficient to induce expression of genes needed for cell division.

### Proteomics reveals stoichiometric down-regulation of negative MAPK pathway regulators in drug-adapted cells

Sensitization of MAPK signaling to growth factors in drug-adapted cells is caused by relief of negative feedback from MAPK activity itself (Lito et al., 2012). Feedback control over MAPK signaling is thought to involve three main mechanisms: (i) dual specificity protein phosphatases (DUSPs), which inactivate MAPK directly (ii) inhibitory phosphorylation of proteins such as EGFR, SOS1 and CRAF; and (iii) proteins that stoichiometrically inhibit assembly of signaling complexes such as sprouty proteins (SPRYs) and MIG6 (Figure 3A; see **Supplementary Table S1** for abbreviations and HUGO names). To establish the concentrations and modification states of these proteins in response to MAPK activity, we performed quantitative targeted mass spectrometry on A375 cells treated for 24 hr with four doses of vemurafenib (from 0.01 to 1 µM), yielding absolute abundances for 21 proteins across a range of MAPK activities (Figure 3B). Phosphorylation stoichiometry at key regulatory sites was also measured by mass spectrometry and mRNA transcript levels by RNA-seq (Figure 3C-E); see also (Shi et al., 2016).

**Figure 3.**
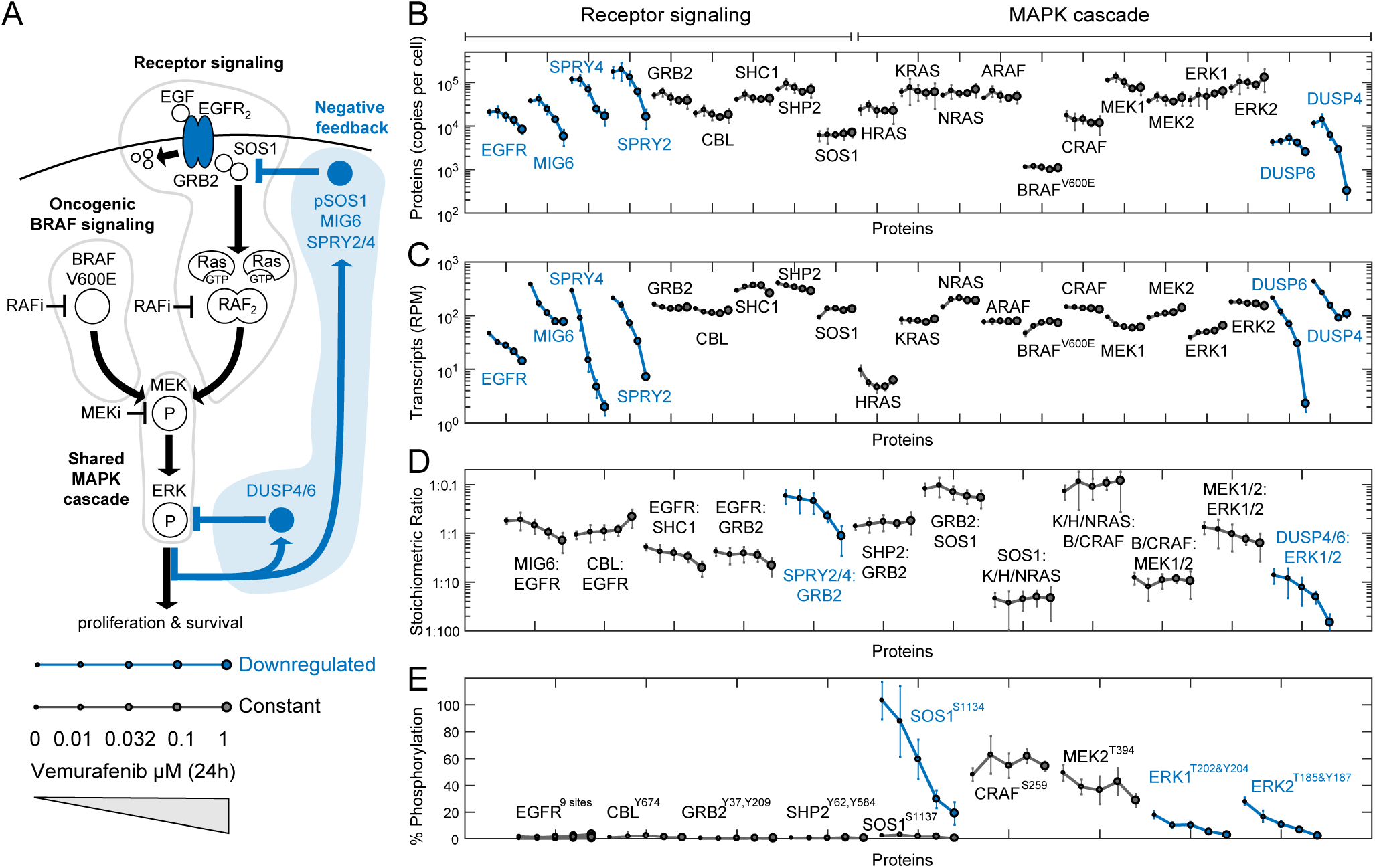
Proteomics reveals stoichiometric down-regulation of negative MAPK pathway regulators in drug-adapted cells. **A)** Schematic of the ERK pathway in BRAF^V600E^ cells; negative regulators of the MAPK pathway are shown in blue. **B)** Absolute abundances (in copies per cell) of selected MAPK pathway proteins. **C)** Changes in mRNA levels for the same proteins as measured by next-generation mRNA sequencing. **D)** Stoichiometric ratios of selected proteins known to interact directly. **E)** Phosphorylation of key regulatory sites on selected proteins. Mean values and standard deviations derive from 4 replicates.

In untreated A375 cells ∼20% of total ERK1^T202,Y204^ and ∼30% ERK2^T185,Y187^ is phosphorylated; this falls to 2– 3% in the presence of 1 µM vemurafenib (matching data from single-cell imaging). The abundance of six proteins also fell with drug exposure, including the EGF receptor (EGFR; maximal fold-change of 2.5), SPRY2/4 (10 and 7-fold change), the ERBB inhibitor MIG6 (6-fold change), and DUSP4/6 (35 and 1.6-fold change) (Figure 3B). RNA-seq data confirmed that these six proteins were controlled at the level of gene expression (Pratilas et al., 2009) (Figure 3C). Phosphorylation of SOS1 also fell nearly 10-fold following vemurafenib treatment (Figure 3E). Thus, MAPK inhibition rapidly and dramatically reduces the levels or activities of three classes of MAPK regulators.

Data on protein abundances and stoichiometric ratios provides insight into MAPK homeostasis (Figure 3B, E). For example, SPRY2 is known to competitively inhibit recruitment of the RAS guanine nucleotide exchange factor SOS1 to GRB2, an adaptor protein that binds to phospho-tyrosine (pY) sites on activated RTKs (Lao et al., 2006). When MAPK signaling was active (vemurafenib absent) the level of SPRY2 plus SPRY4 was in 6-fold excess to GRB2, but in cells exposed to 1 µM vemurafenib, GRB2 was more abundant (Figure 3D). Thus, competitive inhibition of SOS1-GRB2 binding by stoichiometric binding to SPRY proteins is expected to switch from efficient to relatively inefficient. A similar change in relative levels (from ∼1:10 to 1:100) was observed for the DUSP4/6 phosphatases with respect to their pERK1/2 targets. pSOS1^S1134^ levels also decreased dramatically with MAPK inhibition; phosphorylation on S1134 is known to reduce SOS1 activity and SOS1-GRB2 association (Saha et al., 2012);Figure 3D). Thus, negative regulation of the MAPK pathway is relieved by multiple mechanisms in drug adapted BRAF^V600E^ cells. Low levels of phosphorylation on EGFR, CBL, GRB2, SHP2 (<5%) show that mitogenic signaling downstream of RTKs remains largely inactive under these conditions (Lito et al., 2012).

### Receptor-mediated signaling in drug-adapted cells is resistant to both single-agent RAF and MEK inhibitors

To study how drug adaptation makes cells permissive for RTK-mediated signaling, we pre-treated A375 cells for 24 hr to vemurafenib and dabrafenib over a wide range of concentrations, added EGF at a single high concentration and measured ERK and MEK phosphorylation 5 min. later (Figure 4A). Levels of ligand-induced pERK were found to correlate directly with the concentration of RAF inhibitor in the pre-treatment period, and thus, with the magnitude of BRAF^V600E^ inhibition; at the highest drug doses, ERK could be transiently activated well above its levels in the absence of drug (Figure 4A). MEK phosphorylation was also inducible by EGF confirming that the MEK-ERK cascade operates in the normal way (Figure 4A). We observed similar ERK reactivation after stimulation with other three growth factors at clinically relevant drug concentrations (Figure 4B; denoted in gray). Moreover, maximum ERK levels in ERK^high^ cells (a measure of spontaneous ERK reactivation) correlated well with sensitivity to EGF, showing that spontaneous ERK pulses in single-cells are permitted to the same extent as whole-population MAPK activation by exogenous ligands (Figure 4C and **Figure S1B**).

**Figure 4.**
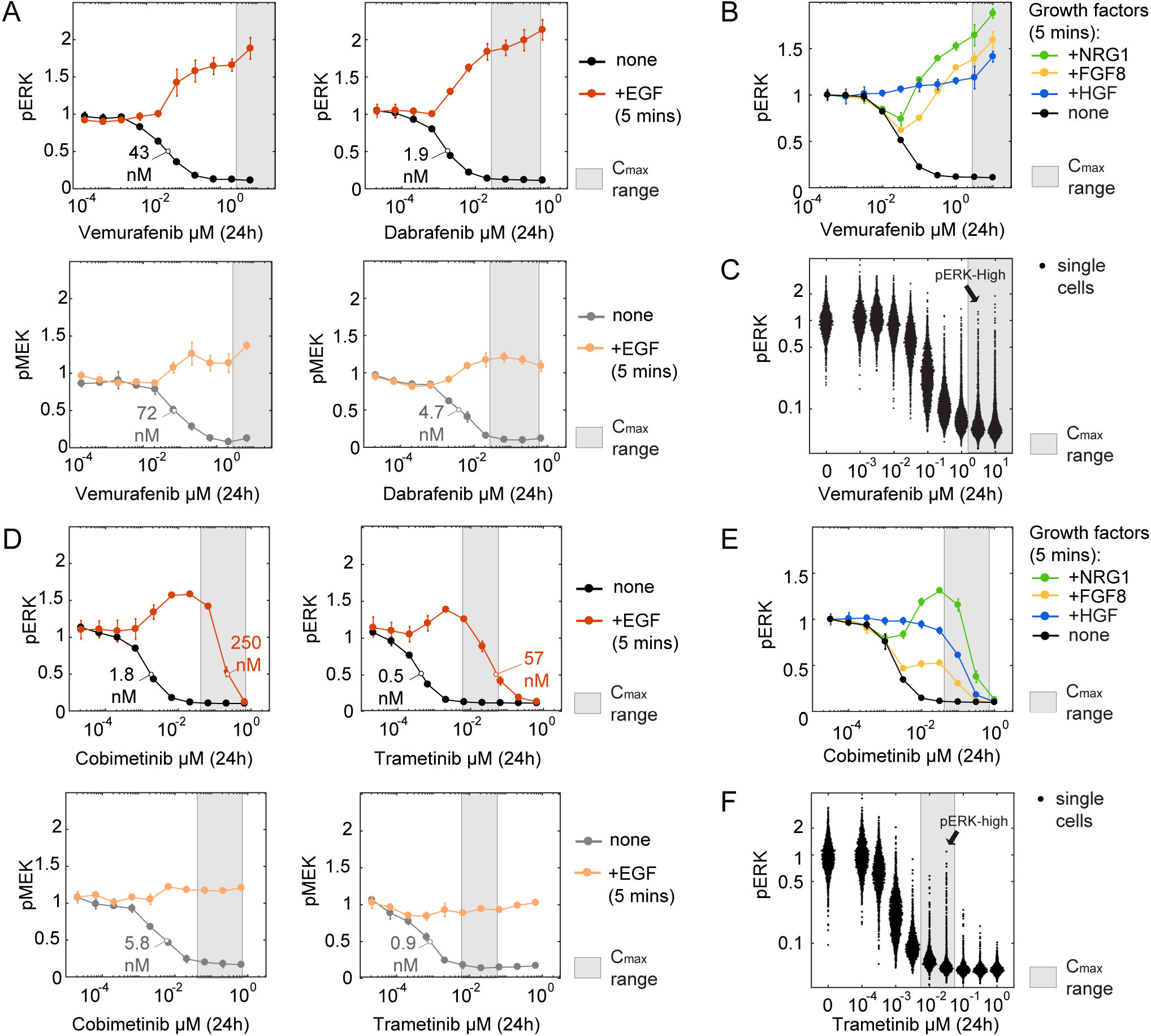
Receptor-mediated signaling in drug-adapted cells is resistant to both single-agent RAF and MEK inhibitors. All data are normalized to contemporaneous DMSO controls; mean values and standard deviations derive from two or more replicate experiments. **A)** pERK and pMEK levels following 24 hr exposure of cells to vemurafenib or dabrafenib over a 10^4^ concentration range followed by addition of EGF (100 ng/mL) for 5 mins. **B)** pERK levels following exposure to vemurafenib for 24 hr over a 10^4^ concentration range followed by addition of one of four growth factors for 5 min. **C)** Distributions of pERK levels in single cells following 24 hr exposure to vemurafenib at the concentrations shown. **D)** pERK and pMEK levels following 24 hr exposure to cobimetinib or trametinib treatment followed by 100 ng/mL EGF for 5 min. **E)** pERK levels following exposure to cobimetinib for 24 hr over a 10^4^ concentration range following by addition of one of four growth factors for 5 min. **F)** Distributions of pERK levels in single cells following 24 hr exposure to trametinib at the concentrations shown.

Since MEK kinases lie downstream of both oncogenic BRAF^V600E^ and receptor-mediated RAF signaling, MEK inhibitors should, in principle, suppress both routes to ERK activation. We found that exposure of A375 cells to cobimetinib reduced pERK and pMEK levels with an IC50 ∼ 1 to 5 nM (Figure 4D). Surprisingly, when cobimetinib-treated cells were exposed to EGF, pMEK and pERK were found to be at least 100-fold more drug resistant. As cobimetinib concentrations increased above 5 nM, pERK levels rose above levels in drug-naïve cells to a maximum at ∼20 nM cobimetinib and then fell again, with an IC50 ∼ 250 nM. pERK was not fully suppressed until cobimetinib concentrations were well above the clinical range (∼1 µM). A similar biphasic pERK response to MEK inhibition was observed for three other growth factors (Figure 4E) and four other MEK inhibitors, including trametinib (Figure 4D, **Figure S3**). In all cases, pMEK levels rose slightly above pre-treatment levels and remained constant even at the highest drug doses tested. Spontaneous pulses in single-cell were similar in magnitude to ligand-induced pERK levels at clinically-relevant doses (Figure 4F). Thus, MEK inhibitors are ∼100-fold less efficient at blocking RTK-mediated ERK activation than BRAF^V600E^-mediated ERK activation.

### A computational model of MAPK regulation identifies mechanisms of resistance due to drug-induced network rewiring

To understand the mechanism responsible for silencing of BRAF^V600^-signaling by RAF/MEK inhibitors under conditions that potentiate ligand-mediated signaling we integrated a wide range of literature-based structural, biochemical and cell-based data in a computational model. This mass-action biochemical model (Melanoma Adaptive Resistance Model; MARM1) included many features of previously published MAPK models (Kholodenko, 2015)(Rauch et al., 2016)(Rukhlenko et al., 2018) and was constructed using the PySB rules-based language (Lopez et al., 2013) extend to support an energy-based description of cooperativity (Sekar et al., 2017)(see **Text S1**). The model included multiple molecular mechanisms relevant to MAPK signaling in BRAF^V600^ melanoma cells: (i) induction of active hetero- and homo-CRAF/BRAF dimers by RAS-GTP binding, downstream of RTKs (Freeman et al., 2013)(ii) a lower affinity of RAF inhibitors for CRAF/BRAF dimers (Yao et al., 2015)(Kholodenko, 2015)(Rukhlenko et al., 2018); (iii) efficient phosphorylation of MEK by RAF dimers but not by BRAF^V600E^ when MEK is bound to a MEK inhibitor (Lito et al., 2014); (iv) a lower affinity of MEK inhibitors for phosphorylated v. unphosphorylated MEK (Hatzivassiliou et al., 2013)(Rukhlenko et al., 2018); (v) negative feedback mediated by SOS1, SPRY and DUSP4/6 (Figure 3) and (vi) growth-factor dependent activation of RTKs that are then inactivated by internalization and degradation (Figure 2E). The resulting model comprised 13 proteins (shown in Figure 5A) and 1007 ordinary differential equations with 70 kinetic parameters; initial conditions were derived from mass spectrometry data and kinetic rates estimated from prior knowledge and signaling dynamics (Figure 4B-D and **Figure S4**) (Fröhlich et al., 2017). Simulations were performed using the 100 best-fit parameter sets to account for uncertainty in parameter estimation (Figure 5A).

**Figure 5.**
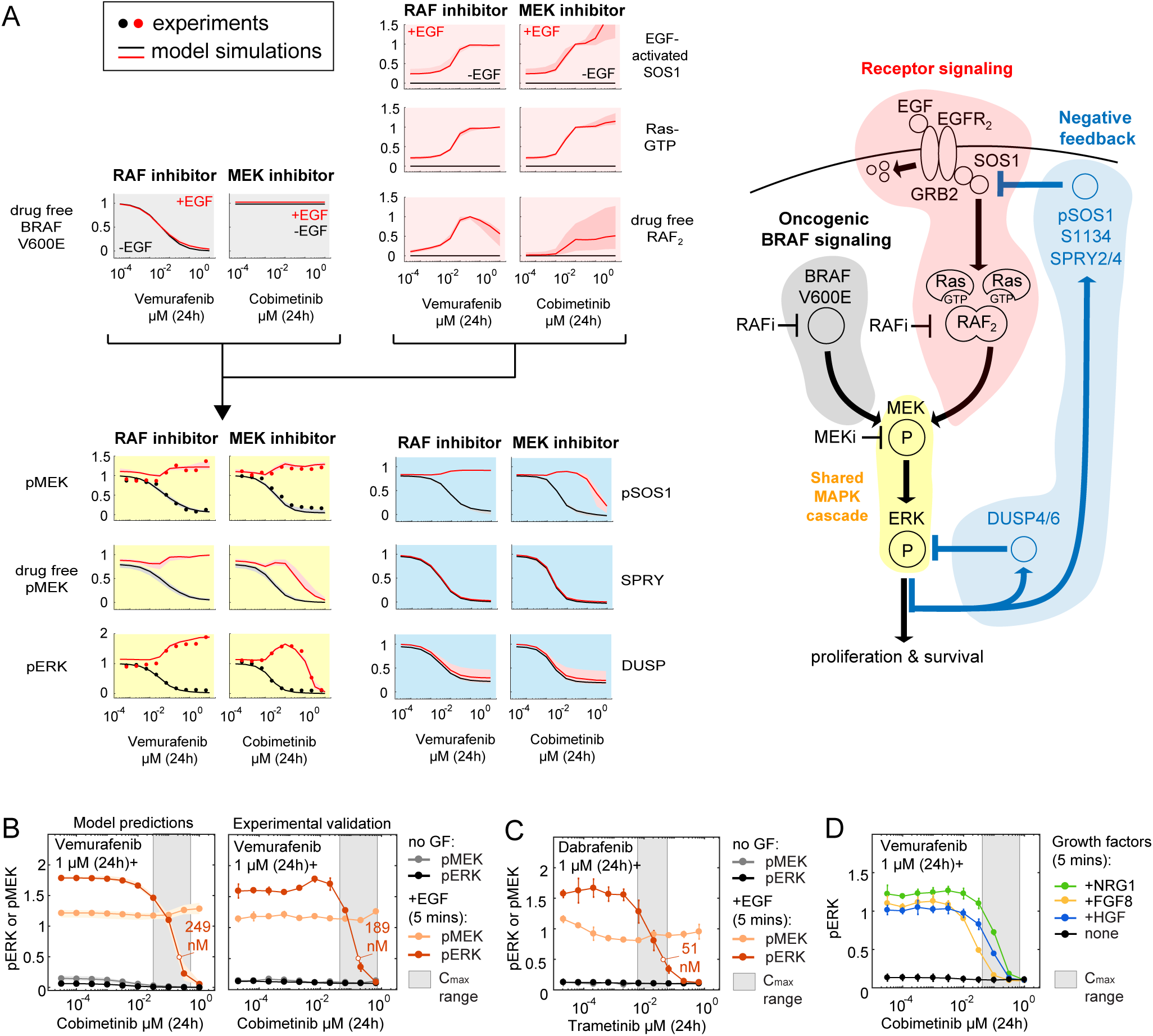
A computational model of MAPK regulation identifies mechanisms of resistance due to drug-induced network rewiring. All data are normalized to contemporaneous DMSO controls; mean values and standard deviations derive from two or more replicate experiments. **A)** Simulation from the MARM1 model showing fold changes of specific proteins and protein complexes in cells treated for 24 hr with vemurafenib or cobimetinib over a 10^4^ concentration range in the presence (red lines) or absence (black lines) of EGF (100 ng/mL for 5 mins). Shading denotes the interquartile range of simulations for 100 best parameter sets. **B).** Model predictions (left panel) and experimental validation (right) for cells treated for 24 hr with vemurafenib (1 uM) plus cobimetinib over a 10^4^ concentration range with or without subsequent addition of EGF (100 ng/mL for 5 min). **C)** Data from cells treated for 24 hr with dabrafenib (1 uM) plus trametinib over a 10^4^ concentration range with or without subsequent addition of EGF (100 ng/mL for 5 mins) **D)** Cells treated for 24 hours with vemurafenib (1 uM) plus cobimetinib over a 10^4^ concentration range with or without addition of EGF, NRG1, FGF8 and HGF (100 ng/mL for 5 mins).

The ability of the MARM1 to reproduce signaling dynamics demonstrates that molecular mechanisms included in the model are sufficient to explain receptor-mediated resistance to RAF and MEK inhibitors (Figure 5A). In drug-naive BRAF^V600E^ cells, pERK levels are chronically high and model analysis showed that exposure to growth factors has little impact on pERK because SPRY2/4 and pSOS1 efficiently block RTK-mediated signaling (Figure 5A).Thus, EGFR (the RTK present in the model) can be activated by dimerization but this does not result in downstream activation of RAS-GTP (Lito et al., 2012). With increasing drug (∼10-50 nM vemurafenib), BRAF^V600E^ and ERK activities are suppressed and negative feedback progressively relieved. This leaves cells in an ERK-low state but with substantially lower concentrations of negative regulators than in the drug-naïve state. Due to reductions in SPRY2/4 and pSOS1, EGFR can dimerize in the presence of ligand and activate RAS, promoting formation of BRAF/CRAF homo- and hetero-dimers (CRAF is 10-fold more abundant than BRAF).

In the model, three mechanisms were responsible for MEK and ERK phosphorylation by BRAF/CRAF dimers in the presence of inhibitory drugs and growth factor. First, RAF dimers were >100-fold more resistant to RAF inhibitors than BRAF^V600E^ monomers due to the stereochemistry of drug binding (Figure 5A, the “drug free RAF2” species in the model). Second, RAF dimers were also substantially more efficient at phosphorylating drug-bound MEK than monomeric BRAF^V600E^; thus, active MEK accumulated and activated ERK at cobimetinib concentrations up to 250 nM (Figure 5A, see “pMEK”). At yet higher drug doses, the substantial larger pool of pMEK generated by RAF dimers was eventually completely bound by the MEK inhibitor and its catalytic activity blocked, causing pERK levels to fell (Figure 5A, see “drug free pMEK”) and giving rise to the biphasic pERK dose-response. However, pMEK levels remained high across the full cobimetinib dose range because pMEK reports on the activity of upstream kinases, in this case ligand-induced RAF dimers. Third, because DUSP levels are ∼2 to 10-fold lower in drug adapted cells, ERK reached higher levels than in untreated cells even at similar levels of MEK phosphorylation. Modeling therefore demonstrates that multiple mechanisms must work in concert to rewire MAPK signaling in the presence of drugs. The fundamental insight from the model is that MAPK reactivation by ligands in drug-adapted melanoma cells is possible because both RAF and MEK inhibitors, albeit for different reasons, selectively inhibit MEK phosphorylation from BRAF^V600E^ while allow efficient MEK phosphorylation from ligand-stimulated RAF dimers.

### Combined RAF and MEK inhibition fails to suppress receptor-driven MAPK reactivation

RAF and MEK inhibitors are usually described as being synergistic with respect to their effects on MAPK activity in patients (Tolcher et al., 2018), however their differential potency on oncogenic and RTK-mediated signaling suggests a more complex reality. In particular, experiments and modelling suggest that RAF and MEK inhibitors are poor inhibitors of RTK-mediated signaling. To assess if RTK-mediated signaling indeed lowers the potency of combined RAF and MEK inhibition, we used MARM1 to predict pERK and pMEK following exposure to cobimetinib and a high dose of vemurafenib (Figure 5B). In cells exposed to EGF for 5 mins, MEK was reactivated ∼10-fold and the IC50 values for pERK increased nearly 100-fold compared to oncogenic signaling. Experiments confirmed these predictions (Figure 5B) as well as similar studies using trametinib plus dabrafenib (Figure 5C) and other growth factors (Figure 5D). Pharmacological interaction was assessed by isobologram analysis over a 10^4^-fold concentration range for vemurafenib and cobimetinib. Modeling and experiments showed that the drugs were strictly additive in the absence of EGF (assayed at 24 hr.; the positive curvature in the isoboles in Figure 6A is due to use of a log10 scale for drug concentration) and that cobimetinib acted as a low potency single agent in the presence of EGF (assayed at 5 min) (Figure 6B). Moreover, at clinically-accessible drug concentrations we found that ERK reactivation by receptors cannot be fully suppressed by combinations of RAF and MEK inhibitors (Figure 6B-C).

**Figure 6.**
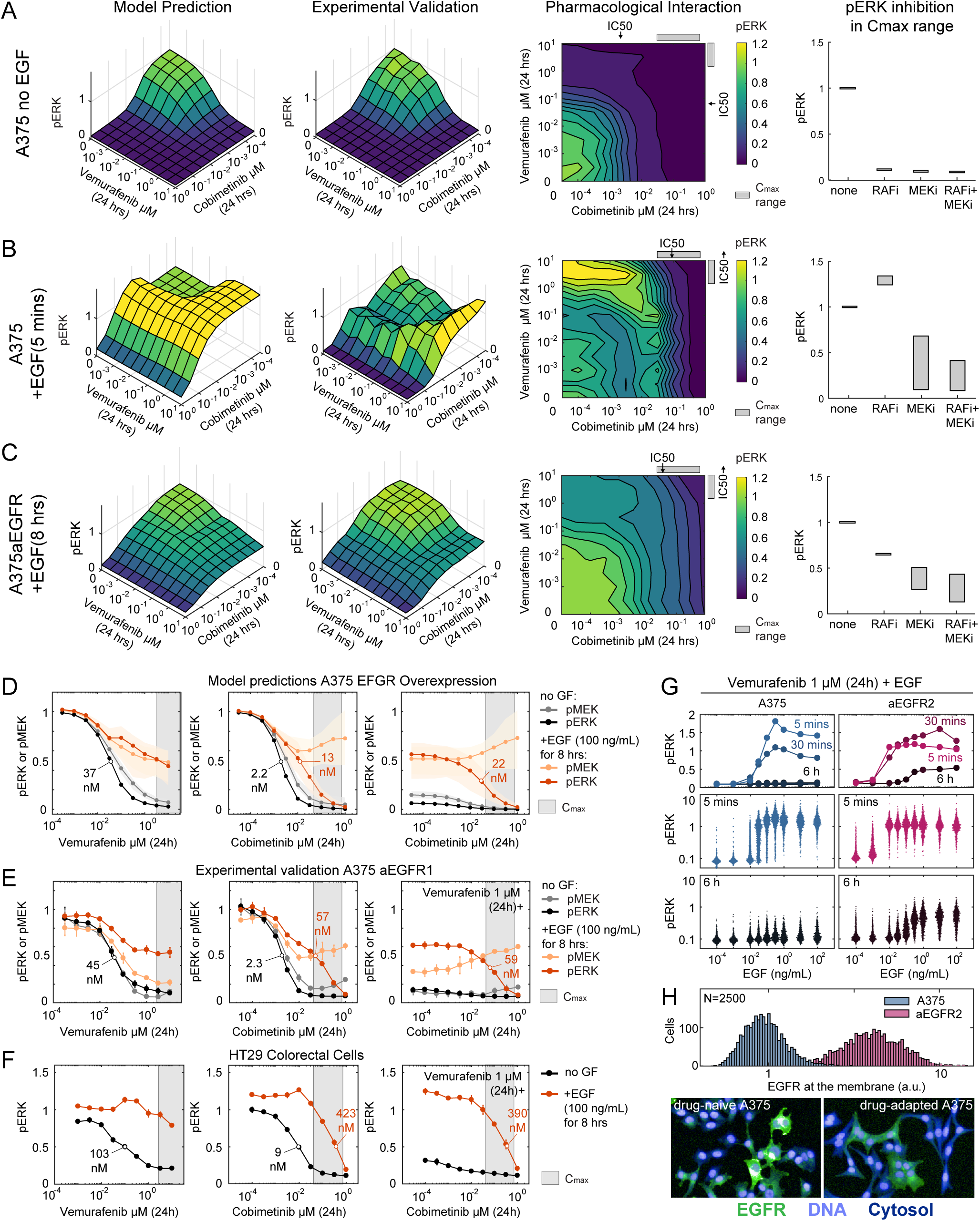
Combined RAF and MEK inhibition fails to suppress receptor-driven MAPK reactivation. All data are normalized to contemporaneous DMSO controls; mean values and standard deviations derive from two or more replicate experiments. **(A-C)** MARM1-based simulation (left panels) and experimental validation (right panels) of pERK levels following 24 hr exposure to vemurafenib plus cobimetinib in: **A)** in parental A375 cells without addition of EGF, **B)** parental A375 cells exposed to EGF (100 ng/mL for 5 mins) following vemurafenib treatment **C)** A375-aEGFR overexpressing cells exposed to EGF (100 ng/mL for 8 hr). Model-based simulation **(D)** and experimental validation **(E)** of pERK and pMEK levels in A375-aEGFR cells treated for 24 hr with vemurafenib or cobimentinib individually or in combination with EGF (100 ng/mL) for 8 h. **F)** HT29 analyzed as in panel E. **G)** EGFR abundance as measured by immunofluorescence microscopy at the membrane of A375 or A375_aEGFR2 cells exposed for 24 hr to 1µM Vemurafenib. **H)** Quantification of pERK in single cells and a population of A375 and A375_aEGFR2 exposed to vemurafenib (1 uM, 24 h) and then to 5 min of EGF over a 10^6^-fold concentration range.

MARM1 predicted, and experiments confirmed, the resistance of sustained ERK activity in EGFR-overexpressing cells to the highest vemurafenib doses tested (10 µM); pERK was also 30-fold more resistant to cobimetinib than in parental A375 cells (Figure 6C). Modeling ascribed this to sustained MEK phosphorylation by BRAF/CRAF dimers (Figure 6D-E). This prediction was confirmed by the ability of LY3009120, but not other RAF inhibitors, to completely suppress pMEK and pERK levels (**Figure S5**); LY3009120 is a pan-RAF inhibitor active on multiple BRAF/CRAF isoforms in their monomeric and dimeric states (Peng et al., 2015). Some BRAF^V600E^ cancers cannot be treated successfully with RAF and MEK inhibitors and HT29 cells are representative of a colorectal line that naturally expresses ∼5 fold more EGFR receptors than A375 cells (**Figure S2D)**(Prahallad et al., 2012). We found that EGF exposure made HT29 highly resistant to vemurafenib and cobimetinib individually and in combination (Figure 6F). This suggests that ligand-induced RAF dimers underlie both transient and sustained receptor-mediated ERK reactivation in BRAF^V600E^ cancer cells under RAF/MEK treatment.

### Spontaneous adaptive ERK pulses occur in spatially localized cell clusters

Which factors contribute to the non-uniformity of spontaneous ERK reactivation in melanoma cell lines? We examined the relationship between EGFR levels and sensitivity to EGF in A375 cells exposed to 1µM vemurafenib for 24 hr. We found that ERK was fully activated at 1 ng/mL EGF (∼0.1 nM) in parental A375 cells whereas in EGFR over-expressing cells (which have ∼8-fold more EGFR), ERK was maximally activated at a 100-fold lower ligand concentration (0.01ng/mL; Figure 6G). Moreover, whereas all EGFR over-expressing cells responded to physiological concentrations of EGF (0.0001-0.01 ng/mL) this was true of only a subset of parental cells (Figure 6G). At a single cell level, EGFR abundance was log-normally distributed in both parental and over-expressing lines, implying natural variability in ligand sensitivity (Shaffer et al., 2017) (Figure 6H). Cells with high receptor levels were also found in spatially localized clusters (Figure 6H). Thus, one explanation for non-homogenous ERK pulses is cell-to-cell variability in receptor levels.

When we examined the spatial organization of ERK pulses in live-cell data we found that ERK was often activated at nearly the same time in groups of neighboring cells, and that synchronously pulsing cells were often in contact (although contact was not necessary; see Figure 7A and **Movie S1**). Fixed cell imaging confirmed the presence of ERK^high^ patches with different shapes and sizes (Figure 7B). These data are most consistent with release and response to autocrine/paracrine factors. Thus, it seems likely that both non-uniform receptor abundance and localized ligand release contribute to ERK^high^ patches. Similar patterns of sender and receiver cells underlie “firework-like” bursts of ERK activity found in the mouse skin cells and mediated by EGFR ligands (Hiratsuka et al., 2015).

**Figure 7.**
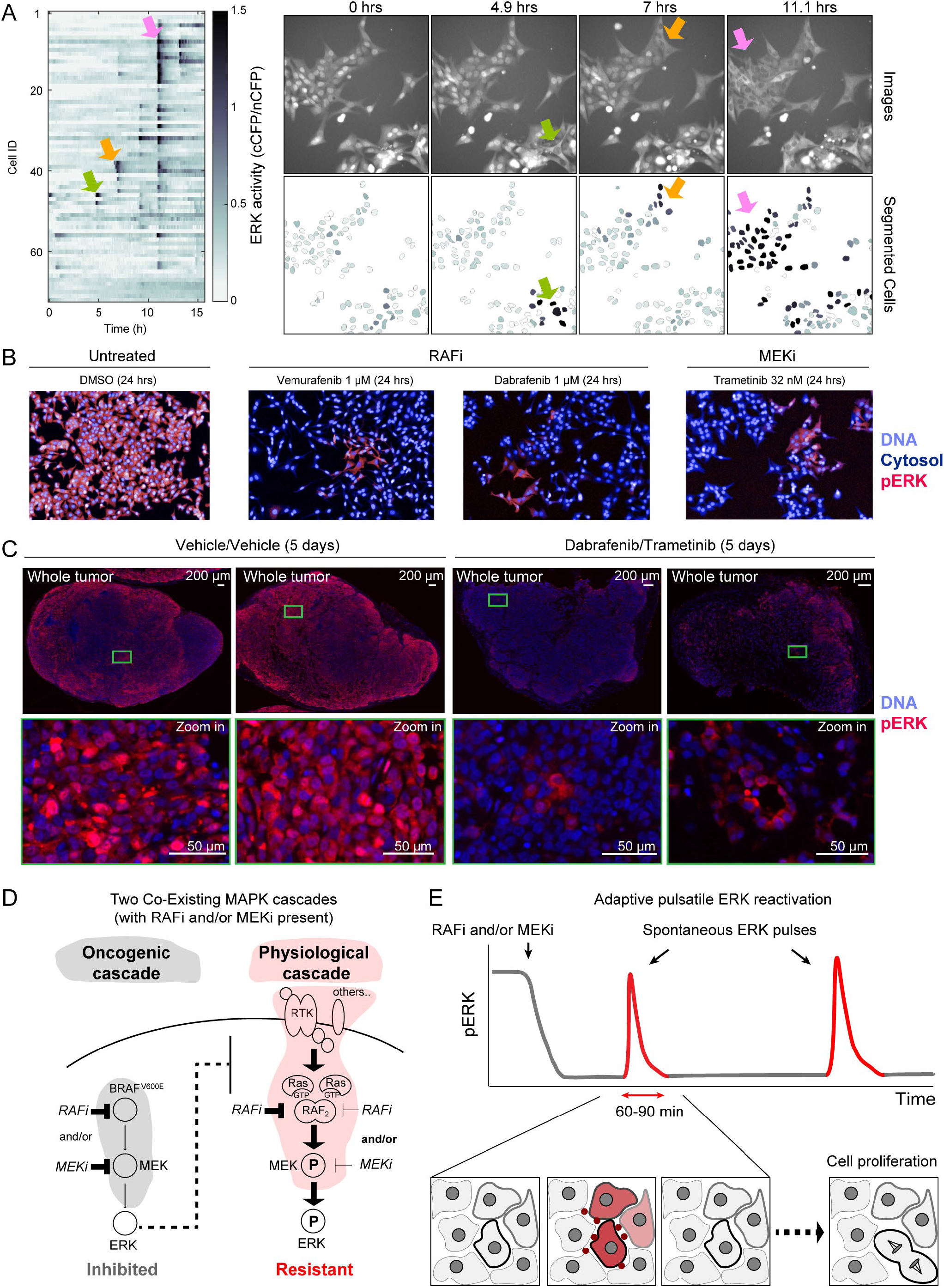
Spontaneous adaptive ERK pulses occur in spatially localized cell clusters. **A)** Heatmap of ERK activity as in Figure 1 for individual cells in one of ten fields of vemurafenib treated cells (1uM, 24h). Arrows denote synchronous ERK pulses. On the right, images show spatial localization of pulsing cells in the ERK:KTR CFP channel (upper row) and corresponding ERK activity (cCFP/nCFP ratio) projected onto the segmentation mask (lower panel). Selected pulsing cells are highlighted in the image, segmentation mask and heatmap. **B)** Immunofluorescence microscopy of pERK in cultured A375 cells treated with DMSO or the RAF or MEK inhibitors for 24 hours. **C)** Immunofluorescence microscopy of pERK in A375 mouse xenografts treated either with vehicle alone or dabrafenib plus trametinib for 5 days. **D)** Schematic of co-existing oncogenic and physiological MAPK cascades in A375 cells. **E)** Schematic of sporadic ERK pulses that induce cell division in the presence of RAF and/or MEK inhibitors.

As a first step in studying ERK reactivation *in-vivo*, we used immunofluorescence imaging to measure pERK levels in A375 mouse xenografts treated for 5 days with dabrafenib plus trametinib. When single cells were segmented and pERK levels quantified (n > 10^4^ cells; 3-4 animals per condition, **Figure S6**) we found that pERK levels were substantially reduced, confirming that RAF and MEK inhibitors had their intended effects (Figure 7C and **Figure S6E**). Under these conditions, pERK^high^ cells were found in small clusters that resembled the clusters found in cultured cells (Figure 7C). Xenograft data are not time-resolved, and we cannot exclude confounding effects such as uneven drug penetration. However, our findings are consistent with the idea that islands of ERK^high^ cells remain in tumors in which average ERK levels have been greatly reduced by RAF and MEK inhibitors.

## DISCUSSION

When BRAF^V600E^ melanoma cells are exposed to RAF/MEK inhibitors, MAPK signaling is inhibited and most cells die or arrest. However, some cells escape and then proliferate as slow-dividing ‘persister’ cells. These cells constitute a potential source of residual disease and a reservoir for the emergence of drug-resistant tumors (Ahn et al., 2017)(Fallahi-Sichani et al., 2017)(Shaffer et al., 2017)(Paudel et al., 2018). The fundamental mystery of the adapted state is how melanoma cells can grow and divide when ERK is profoundly inhibited. Here, we show that localized clusters of drug-adapted cells, in which oncogenic ERK signaling is strongly inhibited, experience sporadic pulses of high ERK activity that are sufficient to promote their division. Computational and experimental analysis of ERK pulsing shows that this is possible because MAPK signaling in BRAF^V60OE^ cells is composed of two co-existing pathways. A pathway driven by oncogenic BRAF^V600E^ monomers chronically activates ERK and is highly sensitive to RAF and MEK inhibitors. A second pathway involving factors in the microenvironment as well as RTKs, RAS-GTP and BRAF/CRAF dimers is highly resistant to inhibitory drugs. Thus, the low potency of RAF and MEK inhibitors on receptor-mediated MAPK signaling (Figure 7D) allows individual melanoma cells to respond to factors in the microenvironment and escape drug-induced cell-cycle arrest via sporadic ERK pulses (Figure 7E).

When RTKs are abundant, MAPK reactivation by the tumor microenvironment is a known source of resistance to RAF inhibitors in BRAF^V60OE^ cancers - by EGFR signaling in colon cancer for example (Prahallad et al., 2012)(Montero-Conde et al., 2013)(Sun et al., 2014). Due to low receptor levels, BRAF^V60OE^ melanomas treated with RAF/MEK inhibitors are thought to lack such a route to resistance. However, our data show that even when RTK levels are low (∼10^3^ molecules per cell) they can drive brief ERK pulses and expression of genes necessary for cell cycle progression, such as EGR1 and cyclin D (Zwang et al., 2011). ERK pulses are known to control cell proliferation (Albeck et al., 2013) and they play a role in both normal homeostasis and wound healing in the skin (Hiratsuka et al., 2015). Our data show that drug-adapted melanoma cells pulse in patches, and live-cell imaging suggests the involvement of autocrine/paracrine signaling (Aoki et al., 2013). We observed patches of ERK^HIGH^ cells in mouse xenografts treated with RAF and MEK inhibitors but further work will be required to show that the underlying mechanisms are similar. Pulsing in response to growth factors is a potential explanation for the demonstrated ability of different microenvironments to confer a growth advantage to BRAF^V600E^ cancers treated with targeted inhibitors (Straussman et al., 2012)(Wilson et al., 2012).

It is possible to induce a ERK pulse in the whole cell population that is similar in duration and magnitude to spontaneous pulses of single cells by adding exogenous growth factors to growth media. In this context, low receptor levels are a likely explanation for the short duration of pulses: once RTKs bind ligand they are efficiently endocytosed and degraded, terminating signaling. It appears that multiple growth factors and receptors can substitute for each other (EGF, NRG1, FGF8, HGF in the current work), perhaps explaining why fully characterizing the microenvironment or blocking its contributions to resistance is challenging (Ahmed and Haass, 2018). The spatial heterogeneity of spontaneous ERK pulsing likely arises from localized release of autocrine/paracrine factors enhanced by low and non-uniform distribution of RTKs at a single cell level (Spencer et al., 2009)(Shaffer et al., 2017). Dying cells are also a likely source for some of these autocrine/paracrine factors including presentation of AXL ligands in the context of phosphatidylserine lipids (Müller et al., 2014). Non-uniform distribution of RTKs (e.g. EGFR and AXL) has been associated with drug resistance *in-vitro* and in patients and may act by increasing the propensity for pulsatile MAPK reactivation (Tirosh et al., 2016)(Shaffer et al., 2017). Additional work is required to determine whether other downstream signaling proteins, such as JNK or AKT, pulse (Fallahi-Sichani et al., 2015).

A key function of MEK inhibitors in RAF/MEK combination therapy is to prevent MAPK reactivation (Chapman et al., 2014). However, we find that MEK inhibitors are 100-fold less potent on receptor-mediated signaling than on oncogenic signaling, making them unlikely to fully inhibit adaptive MAPK reactivation at a clinically accessible dose range. To understand precisely how this arises, we constructed and analyzed a computational model of MAPK signaling that integrates ‘thermodynamic, kinetic, structural and mutation information’ (Rukhlenko et al., 2018) and is trained on quantitative proteomics and signaling data. The model extends foundational work by Kholodenko and colleagues (Kholodenko, 2015)(Rukhlenko et al., 2018) and reproduces the responses of MAPK signaling to single-agent RAF and MEK inhibitors and predicts their combination. Model analysis shows that efficient MEK phosphorylation by receptor-mediated RAF dimers is the likely mechanism of resistance to both RAF and MEK inhibitors. Relief of negative feedback and resistance of BRAF/CRAF dimers to RAF inhibition are well established contributors to drug resistance (Lito et al., 2012). The precise mechanisms involved in the selective disruption of BRAF^V600E^-driven MEK phosphorylation by MEK inhibitors are less understood, and we pragmatically described a mechanism in which the MEK inhibitor protects MEK from phosphorylation by BRAF^V600E^. Previous studies suggest a potential role for drug-induced changes in the stability of RAF-MEK complexes (Hatzivassiliou et al., 2013)(Lito et al., 2014), in MEK dimerization (Yuan et al., 2018) and in association with KSR scaffolding proteins (Dhawan et al., 2016). Our use of a rule-based framework makes extending the scope and mechanistic details in MARM1 straightforward, contingent on new mechanistic evidences.

The pharmacological consequences of differences between oncogenic and receptor-mediated signaling are striking. Despite the general assumption that RAF and MEK inhibitors are synergic in their pharmacology, we observe potent additivity on oncogenic BRAF^V600E^ signaling and low potency single-agent activity (for MEK inhibitors) on receptor-driven signaling. Computational modeling predicts, and experiments confirm the drug resistance of cells expressing high EGFR levels (e.g. BRAF^V600E^ colorectal cancers; (Kirouac et al., 2017)). Since resistance is ascribed to Ras-GTP activation, similar mechanisms may underlie the inefficacy of MEK inhibitors in RAS-mutant cancers (Hatzivassiliou et al., 2013)(Lito et al., 2014). Because many receptors can activate the MAPK pathway, inhibitors of individual RTKs are unlikely to be broadly effective in reducing resistance to RAF and MEK inhibitors. Drugs acting downstream of multiple RTKs such as SHP2 inhibitors (Chen et al., 2016) might block adaption, as might RAF dimer inhibitors (Rukhlenko et al., 2018), and MEK inhibitors effective in suppressing MEK phosphorylation by RAF dimers. However, it seems likely that the differences between the oncogenic and physiological MAPK signaling uncovered in this work may actually underlie the good tolerability of both MEK and RAF inhibitors in patients (Chapman et al., 2011)(Flaherty et al., 2012). Drugs able to fully repress both cascades would likely suppress resistance but will almost certainly be more damaging to non-transformed cells by disrupting their physiological signaling. Thus, the achilles heel of MAPK inhibitors with respect to resistance might very well be their saving grace with respect to tolerability.

## Supporting information

Key Resource Table

## ACKNOWLEDGMENTS

We thank S. Boswell, A. Sokolov, Z. Maliga, C. Yapp, M. Chung, M. Fallahi-Sichani, M. Rogava, I. Gasic and R. Madsen for their help and G. Lahav and the Nikon Imaging Center at HMS for assistance with microscopy and the O2 High Performance Compute Cluster for computing support. The work was funded by NCI grant U54-CA225088 to PKS, U54-CA210180 to DAL, U01-CA227544 to WJQ and HSW, a NIGMS grants P41GM103493 to TS, LY and CDN, a Novartis Foundation fellowship to LG, a HFSP grant LT000259/2019-L1 to FF and a SNSF Early Postdoc Mobility fellowship P2ZHP3_181475 to D.S. Proteomics work was performed at EMSL, a national scientific user facility sponsored by the DOE under Contract DE-AC05-76RL0 1830.

## AUTHOR CONTRIBUTIONS

LG conceived and designed the study. LG, CC, GS, SKL, JM, JYC, GJB, DS, AC and DAL generated reagents and performed experiments. LG performed data analysis. LG and FF performed model analysis. DL, TS, LY, CDN, WQ and HSW performed proteomics studies. PKS supervised the work and LG and PKS wrote the manuscript. All authors reviewed and approved the final version.

## Methods

### Contact for Reagent and Resource Sharing

Further information and requests for resources and reagents should be directed to and will be fulfilled by the Lead Contact, Peter Sorger.

### Experimental Model and Subject Details

#### Cell lines and tissue culture

Cell lines used in this study were obtained from the Massachusetts General Hospital Cancer Center and originated from the following primary sources: A375, C32, K2, RVH421, WM115, SKMEL28, and WM1552C (ATCC); MMACSF (RIKEN BioResource Center); and MZ7MEL (Johannes Gutenberg University Mainz). HT29 was obtained from Merrimack Pharmaceuticals and HEK293T was from ATCC. C32, K2, MMACSF, SKMEL28, RVH421, and WM115 cell lines were grown in DMEM/F12 (Invitrogen) supplemented with 5% heat inactivated fetal bovine serum (FBS) (Gibco) and 1% sodium pyruvate (Invitrogen). MZ7MEL and WM1552C were grown in RPMI-1640 (Corning) supplemented with 5% FBS and 1% sodium pyruvate (Invitrogen). A375 cells were grown in Dulbecco’s modified eagle medium with 4.5 g/l D-glucose, 4 mM L-glutamine, and 1 mM sodium pyruvate (DMEM) (Corning), supplemented with 5% FBS. HEK293T cells were grown in DMEM supplemented with 10% FBS. Penicillin and streptomycin were added to all growth media at final concentrations of 100 U/mL and 100 μg/mL, respectively (Corning). Cells were tested for mycoplasma contamination using the MycoAlert mycoplasma detection kit (Lonza).

#### Xenografts

Formalin-Fixed Paraffin-Embedded (FFPE) tissue slides of A375 xenografts were prepared as previously described (Fallahi-Sichani et al., 2017). Briefly, six week-old NU/J mice (Jackson Laboratory, Stock# 002019) were injected subcutaneously in the right flank with 2.5 million A375 cells. Tumor xenografts were allowed to grow and mice were then randomly split into two groups. The first group were treated daily for 5 days with 200 μL dabrafenib (25 mg/kg) plus trametinib (2.5 mg/kg) via oral gavage (OG) and with an intraperitoneal (IP) injection control (DT treated group). The second group were given equivalent volumes of OG and IP vehicle controls (VV control group). The vehicle for OG was 0.5% hydroxypropylmethylcellulose, 0.2% Tween 80 in pH 8.0 distilled water. Tumor volume was calculated from daily measurements by caliper (see **Figure S6B**). After 5 days, mice were transcardially perfused with oxygenated and heparinized Tyrode’s solution which allowed for simultaneous euthanasia and exsanguination. Flank xenografts were then surgically removed and fixed in 4% paraformaldehyde (PFA) in PBS and stored at 4 °C for 48 h. All fixed tumors from a single group were uniformly paraffin embedded into a single block holder and sectioned in 5 μm slices.

### Method Details

#### Construction of CRISPRi and CRISPRa A375 cell lines

All lentiviral particles were produced in HEK293T cells transfected with the lentiviral plasmid of interest (as mentioned in relevant sections below), psPAX2 (Addgene #12260) and pCMV-VSV-G (Addgene #8454) in a 2:2:1 molar ratio using lipofectamine 3000 (Invitrogen) according to the manufacturer’s instructions. After 2 days, growth medium containing the lentiviral particles was harvested, centrifuged, filtered through a 0.45 μm low-protein binding membrane and stored at −80 °C.

To generate an A375 cell line stably expressing dCas9-KRAB (A375_i), A375 cells were transduced with lentiviral particles produced using vector pMH0001 (Addgene #85969; expresses dCas9-BFP-KRAB from a spleen focus forming virus (SFFV) promoter with an upstream ubiquitous chromatin opening element) in the presence of 8 μg/mL polybrene (Sigma). A pure polyclonal population of dCas9-KRAB expressing cells was generated by 2 rounds of fluorescence activated cell sorting (FACS) gated on the top half of BFP positive cells (BD FACS Aria II). The performance of A375_i in knocking down endogenous genes was evaluated by individually targeting 3 control genes (ST3GAL4, SEL1L, DPH1) and measuring gene expression changes by RT-qPCR (dataset provided in Synapse database, see **Table S4** and **Data availability**).

To generate an A375 cell line stably co-expressing dCas9 fused to SunTag, and a SunTag-binding antibody fused to the VP64 transcriptional activator (A375_a), A375 cells were first transduced with lentiviral particles produced using vector pHRdSV40-dCas9-10xGCN4_v4-P2A-BFP (Addgene #60903; expresses dCas9 tagged with 10 copies of the GCN4 peptide v4 and BFP) in the presence of 8 μg/mL polybrene. After selection of BFP positive cells using one round of FACS, cells were transduced with lentiviral particles produced using vector pHRdSV40-scFv-GCN4-sfGFP-VP64-GB1-NLS (Addgene #60904; expresses a single-chain variable fragment that binds to the GCN4 peptide fused to GFP and VP64) in the presence of 8 μg/mL polybrene. Single cells with high GFP levels (top 25% of GFP positive cells) and high BFP levels (top 50% of BFP positive cells) were isolated by FACS. Monoclonal cell lines were expanded and a single clone exhibiting robust growth and robust overexpression of target genes was selected as cell line A375_a. The performance of A375_a in overexpressing endogenous genes was evaluated by individually targeting 3 control genes (CDKN1C, SLC4A1, POU5F1) and measuring gene expression changes by RT-qPCR (dataset provided in Synapse database, see **Table S4** and **Data availability**).

#### *CRISPRi and CRISPRa A375* cell *lines targeting EGFR*

Pairs of complementary synthetic oligonucleotides (Integrated DNA Technologies) forming sgRNA protospacers flanked by BstXI and BlpI restriction sites were annealed and ligated into BstXI/BlpI double digested plasmid pU6-sgRNA EF1Alpha-puro-T2A-BFP (Addgene #60955). Protospacer sequences used to target individual genes and synthetic oligonucleotides used to build them are listed in Table S2. The sequence of all sgRNA expression vectors was confirmed by Sanger sequencing. Lentivirus particles were produced using these vectors as described earlier.

A375_i and A375_a cells were infected with sgRNA expression vectors by addition of lentivirus supernatant to the culture medium in the presence of 8 μg/mL polybrene. Transduced cells were selected using puromycin (1.0 μg/mL) starting 48 hr post-transduction and over the course of a minimum of 7 days with daily addition of the antibiotic. After 24 hr growth in puromycin-free medium, 1.0E5 cells were harvested and total RNA was extracted using the RNeasy Plus Mini kit (Qiagen). cDNA was synthesized from 0.1 μg total RNA using Superscript IV reverse transcriptase (Invitrogen) and oligo(dT)20 primers (Invitrogen), following the manufacturer’s instructions. Reactions were diluted four-fold with water and qPCR was performed in 10 μL reactions in 384-well plates using PowerUp SYBR Green PCR Master mix (ThermoFisher Scientific), 2 μL diluted cDNA preparation, and 0.4 μM of primers using a QuantStudio 6 real-time PCR system (Applied Biosystems). All qPCR primers are listed in the dataset provided in Synapse database, see **Table S4** and **Data availability**. To calculate changes in expression level of target genes, all gene specific Ct values were first normalized to GAPDH Ct values (ΔCt). Log2 fold changes in expression were then determined by the difference between the ΔCt value of targeting sgRNAs and that of a non-targeting negative control sgRNA (A375_iNC or A375_aNC) (ΔΔCt).

#### Generation of clonal cell lines

A375 cells were grown for at least 2 passages to 80% confluence in a 10 cm dish. Cells were washed with PBS and then incubated with 0.25% trypsin/2.21 mM EDTA (Corning) for 2 min. After adding 20 mL complete medium, cells were thoroughly homogenized into a single cell suspension. Cell count and homogeneity was determined using a TC20 automated cell counter (Bio-Rad). The cell suspension was serially diluted with medium adjusted to 10% FBS to a final concentration of 20 cells in 15 mL and 150 μL of that dilution was dispensed in wells of a 96-well plate (0.2 cells per well). After about 14 days, wells that showed clonal growth (15-20 per 96-well plate) were expanded by passaging cells into larger dishes in complete medium.

#### ERK activity fluorescent reporter cell line

A375 cells were transfected with plasmids 4_pPB_ERKKTRmTq2_H2BVenus_mCherryGeminin (Fallahi-Sichani et al., 2017) and pCMV_hyPBase (Fallahi-Sichani et al., 2017) at a 5:2 ratio (w/w) using lipofectamine 3000 according to the manufacturer’s instructions. Transfected cells were selected using puromycin (1 μg/mL) starting 48 hr after transfection and over the course of 7 days. A polyclonal population of cells expressing all 3 fluorescent proteins (mturquoise2, Venus and mCherry) was subsequently generated by two rounds of FACS (BD FACS Aria II).

#### Drugs and growth factors

The following chemicals from MedChem Express were dissolved in dimethyl sulfoxide (DMSO) at 10 mM: vemurafenib, dabrafenib, PLX8394, AZ628, LY3009120, cobimetinib, trametinib, selumetinib, binimetinib, PD0325901, lapatinib, erlotinib, SHP099. The following ligands were from Peprotech (catalogue number): EGF (100-15), NRG1 (100-03), FGF8 (100-25A), HGF (100-39). All ligands were prepared in media supplemented with 0.1% bovine serum albumin.

#### Immunofluorescence staining, quantitation, and analysis for cell cultures

The following primary and conjugated antibodies with specified vendor, animal sources and catalogue numbers were used in immunofluorescence analysis of cells and tissues at the specified dilution ratios: p-ERKT202/Y204 rabbit mAb (Cell Signaling Technology, clone D13.14.4E, Cat# 4370), 1:800; p-MEKS217/221 rabbit mAb (Cell Signaling Technology, Cat# 9121), 1:200; p90RSKT359 rabbit mAb (Cell Signaling Technology, D1E9, Cat# 8753), 1:400; EGFR mouse mAb (Thermo Fisher, 199.12, Cat# MA5-13319), 1:100; pS6S240/S244 Alexa Fluor 488 Conjugate rabbit mAb (Cell Signaling Technology, D68F8, Cat# 5018), 1:800; pS6S235/S236 Alexa Fluor 555 Conjugate rabbit mAb (Cell Signaling Technology, D57.2.2E, Cat# 3985), 1:400. Immunofluorescence assays for cultured cells were performed using cells seeded in either 96-well plates (Corning Cat#3603) or 384-well plates (CellCarrier Cat#6007558) for 24 hr and then treated with compounds or ligands either using a Hewlett-Packard D300 Digital Dispenser or by manual dispensing.

Cells were fixed in 4% PFA for 30 min at room temperature (RT) and washed with PBS with 0.1% Tween-20 (Sigma) (PBS-T), permeabilized in methanol for 10 min at RT, rewashed with PBS-T, and blocked in Odyssey blocking buffer (OBB LI-COR Cat. No. 927401) for 1 hr at RT. Cells were incubated overnight at 4 °C with primary antibodies in OBB. Cells were then stained with rabbit and/or with mouse secondary antibodies from Molecular Probes (Invitrogen) labeled with Alexa Fluor 647 (Cat# A31573) or Alexa Fluor 488 (Cat# A21202) both at 1:2000 dilution. Cells were washed with PBS-T and then PBS, and were next incubated in 250 ng/mL Hoechst 33342 and 1:2000 HCS CellMask™ Blue Stain solution (Thermo Scientific) for 20 min. Cells were washed twice with PBS and imaged with a 10× objective using a PerkinElmer Operetta High Content Imaging System. 9–11 sites were imaged in each well for 96-well plates and 4-6 sites for 384-well plates.

Image segmentation, analysis, and signal intensity quantitation were performed using the Columbus software (PerkinElmer). Cytosol and nuclear areas were identified by using two different thresholds on the CellMask™ Blue Stain (low intensity) and Hoechst channels (∼100-fold more intense) and cell boundaries were used to define membrane (2 pixels surrounding cytosolic area), cytosolic and nuclear cell masks (see **Figure S6A** for examples). Cells were identified and enumerated according to successful nuclear segmentation. Apart when otherwise specified, immunofluorescence quantifications are average signals of the cytosolic area. Population average and single cell data were analyzed using custom MATLAB 2017a code. Single cell density scatter plots were generated using signal intensities for individual cells.

#### EdU incorporation experiments

Before immunofluorescence staining (as above), cells were pulsed for 3 hr with EdU (Lumiprobe, Hunt Valley, MD). After fixing, antibody staining and image segmentation, DNA content was quantified by the total Hoechst intensity within the nuclear mask and EdU incorporation was quantified by the average intensity within the nuclear mask. The threshold for identification of EdU positive cells was determined by evaluating DMSO control cells (Edu intensity of 300 a.u.).

#### Immunofluorescence staining, quantitation, and analysis of FFPE xenograft tissue slides

Dewaxing, antigen retrieval and staining of FFPE samples was performed as previously described (Lin et al., 2018) with the following modifications. Samples were subjected to three rounds of photochemical bleaching, then stained with Hoechst 33342 (2 µg/mL) in OBB and antibodies against p-ERK (1:200, Cell Signaling Technologies, Cat. No. 4370S), then anti-rabbit IgG (1:2000, Thermo Fisher, Cat. No. A31573) and finally S100 (1:200, Abcam, ab207367), p-S6 (1:100, Cell Signaling Technologies, Cat. No. 3985S). Slides were imaged using a CyteFinder fluorescence microscope (RareCyte) with 10× (NA=0.3) and 40× (NA=0.6) objectives. We generated whole tumor images using Ashlar, a software tool that performs simultaneous stitching and registration of cyclic microscopy images of large tissue sections. Ashlar uses phase correlation between individual images and a statistical model of microscope stage behavior to correct for stage positioning error and to produce a single seamless mosaic image across all imaging cycles (https://github.com/labsyspharm/ashlar). We combined Ilastik1.3.2 (Sommer et al., 2011) and CellProfiler3.1.8 (Lamprecht et al., 2007) to enable robust single cell segmentation across all xenograft samples (see **Figure S6C** for examples). We used random forest classification implemented in Ilastik to train three distinct classes (nuclei, membrane, and background) to create probability maps for each class. Large images were divided into two images (left and right) to improve performance. Subsequently, CellProfiler was used to segment those probability maps to create labeled single cell masks. Intensity of pERK and S100 staining was quantified within the cellular masks and CSV files were created for downstream analysis using histoCAT (Schapiro et al., 2017).

#### Western Blot staining, quantitation, and analysis

Cells were harvested at 70-80% confluence and lysates were prepared by incubation in Laemmli Sample Buffer (Bio-Rad, #1610737) for 5 min at 95 °C. Lysates were subjected to SDS-PAGE using 4-20% polyacrylamide gels (Bio-Rad, #456-9036). After transfer onto nitrocellulose, proteins were detected using primary antibodies from Cell Signaling (EGFR, #4267, 1:1000) and Santa Cruz Biotechnology (Actin-HRP, #sc-47778 HRP, 1:5,000) and secondary HRP-conjugated antibodies from Cell Signaling (#7074 and #7076, 1:10,000). HRP was detected using ECL substrate purchased from Thermo Scientific (#34076) using a myECL Imager, and signals were quantified using the Image Studio Lite software (LI-COR Biosciences) by normalizing the specific signal for each sample to the Actin signal.

#### Receptor quantification by Luminex bead-based ELISA

Absolute receptor abundances for EGFR, Her2, Her3 and c-Met were quantified using a Luminex bead-based ELISA procedure as previously described (Claas et al., 2018). A375 cells were cultured in the presence or absence of vemurafenib (1 μM) for 24 hr and then one of four growth factors (EGF, NRG1, FGF8 and HGF at 100 ng/mL) was added for 24 hr. Cells were lysed in 50 μL NP40 lysis buffer (20 mM Tris–HCl, 150 mM NaCl, 2 mM EDTA, 1% NP40, 10% Glycerol, pH 7.4) at 4 °C and lysates were were clarified by centrifugation for 15 min at ∼2300 ×g and stored at - 20 °C prior to use. For Met, EGFR, Her2 and Her3, capture antibodies (MAB3581, AF231, MAB1129, and MAB3481, respectively), biotinylated detection antibodies (BAF358, BAF231, BAF1129, and BAM348, respectively) and recombinant proteins used for quantification (8614-MT, 344-ER, 1129-ER, and 348-RB respectively) were from R&D systems. Streptavidin Phycoerythrin (SAPE) was from Biorad (#171304501).

Capture antibody conjugated beads were generated by incubation of MagPix beads (Luminex Corp.) with 5 mg/mL EDC (N-(3-dimethylaminopropyl)-N’-ethylcarbodiimide, Sigma) and 5 mg/mL S-NHS (N-hydroxysulfosuccinimide, Pierce) in 80 mM NaH2PO4 pH 6.3 for 20 min in the dark at RT. After washing, the beads were incubated with 0.1 mg/mL capture antibody in 50 mM HEPES, pH 7.4 at 4 °C overnight. After washing, beads were stored at 4 °C in PBS + 1% BSA and 1% Tween20. After addition of conjugated beads to wells of a 384-well Optiplate (Perkin Elmer) and washing with assay buffer (PBS + 0.1% BSA and 0.1% Tween20), cell lysate samples were added and incubated overnight at 4 °C with shaking at ∼8000 rpm. Samples were diluted if necessary to be in the log-linear range of the standard curve determined with recombinant proteins. After washing, detection antibodies were diluted 1000× in assay buffer and incubated at RT for 1 hr with the sample. After washing, SAPE was diluted 100× in assay buffer, added to wells and incubated at RT for 15 min. After final washing, samples were read on a Flexmap 3D machine (Luminex Corp) according to manufacturer’s protocol.

#### Live-cell microscopy

A375 cells expressing the ERK activity fluorescent reporter (see above) were plated on poly-D-lysine-coated glass bottom dishes (MatTek, P35G-0.170-14-C) 48 hr prior to imaging at a density of 30,000 cells per dish. Drugs were added 24 hr before imaging. Time lapse imaging was performed in DMEM without phenol red with 4.5 g/l glucose, L-glutamine, and sodium pyruvate (Corning) and supplemented with 5% FBS. Cells were imaged on a Nikon Eclipse TE-2000 inverted microscope with a 20× plan apo objective (NA 0.7 5) with a CCD camera. The microscope was enclosed with an environmental chamber to maintain humidity, a temperature of 37°C and 5% CO2. Images were captured in CFP, YFP and Texas Red channels every 6 min using the MetaMorph software (Molecular Devices). The filter sets used were as follows: CFP– 436/20nm, 455nm, 480/40nm (excitation filter, beam splitter, emission filter); YFP – 500/20nm, 515 nm, 520nm; Texas Red – 560/40nm, 585nm, 630/75nm; All filters were obtained from Chroma. Cell tracking and data analysis was performed using custom-made MATLAB scripts (Cappell et al., 2016).

#### Targeted proteomics quantification of protein abundance and phosphorylation

Targeted quantification of protein abundances and phosphorylation was performed as previously described (Shi et al., 2016). Briefly, cell pellets from A375 cell lines treated with different doses of vemurafenib were lysed in 100 μL of lysis buffer containing 8 M urea in 100 mM NH4HCO3 (pH 7.8). Proteins were reduced by 5 mM dithiothreitol for 1 hr at 37 °C and alkylated using 20 mM iodoacetamide for 1 hour at RT in the dark. Samples were diluted eightfold with 50 mM NH4HCO3 and digested by sequencing grade modified trypsin at a 1:50 enzyme-to-protein ratio (w/w) at 37 °C for 3 hr. Each sample was then desalted by C18 solid phase extraction and concentrated to ∼100 μL. The final peptide concentration was measured using bicinchoninic acid assay with an average of ∼4 µg/µL. 10 µg and 100 µg of the peptide mixture per sample were used with the addition of 200 fmol and 50,000 fmol of crude heavy peptides for quantification of protein abundance and protein phosphorylation dynamics, respectively.

For protein abundance quantification (Shi et al., 2016)(Yi et al., 2018), crude heavy-isotope labeled synthetic peptides were purchased from Thermo Scientific and the two best response peptides were selected to configure final selected reaction monitoring (SRM) assays for each target protein. All samples were measured by regular LC-SRM using the scheduled SRM algorithm (Shi et al., 2017) for simultaneous quantification of the selected target proteins. For targeted quantification of phosphorylation (Yi et al., 2018), phosphopeptides were selected for core component proteins for the EGFR-MAPK pathway. Crude heavy isotope-labeled phosphopeptides were purchased from New England peptides and spiked into the peptide sample prior to phosphopeptide enrichment. Phosphopeptides were enriched by immobilized metal-ion affinity chromatography (IMAC) with Fe3+-NTA agarose beads. Eluted phosphopeptides were dried down and stored at −80 °C until further LC-MS/MS analysis. Lyophilized phosphopeptides were reconstituted in 0.1% FA and subjected to LC-SRM analysis immediately. All LC-SRM measurements were performed using the nanoACQUITY UPLC system coupled online to a TSQ Vantage triple quadrupole mass spectrometer (Thermo Scientific). SRM data were analyzed using Skyline software (MacLean et al., 2010) and the best transitions without interferences were used for quantification. The SRM peak area ratios of the endogenous light peptides over heavy peptide standards (i.e., the L/H ratio) were reported for all SRM measurements. The SRM proteomics data were deposited in the Panorama public database (https://panoramaweb.org/fWeE7i.url).

#### Transcriptomics

A375 cells were plated in 24-well plates (Corning, #353047) with 475 uL of growth media and allowed to adhere for 24 hours till 50% confluency. Cells were then treated with vemurafenib alone or in combination with cobimetinib and stimulated with EGF (100 ng/mL) at the times indicated by the experimental design. Each condition was performed twice on two different days, for a total of four replicates, two biological with two technical replicates each. Cells were lysed in 96 well plates and RNA was extracted using Applied Biosystems MagMax 96 total RNA isolation kit (Thermo Fisher, # AM1830) with on bead DNase digestion. Sample concentrations were determined by Nanodrop and RNA quality was assessed on a subset of samples by Bioanalyzer (Agilent); all samples scored RINs of > 9.0. RNA sequencing library preparation was performed with the High Throughput TruSeq Stranded mRNA Library Prep Kit (Illumina, # RS-122-2103) following the manufacturer’s protocol at half reaction volume. Input for each sample consisted of 500 ng of RNA and 5 μL of 1:500 diluted ERCC spike-in mix 2 (Ambion). Libraries were amplified for 11 cycles during the final amplification step. Libraries were quantified using the Qubit dsDNA HS assay (Thermo Fisher Scientific). Library size and quality were spot checked for a subset of samples by Bioanalyzer (Agilent). The average size of cDNA fragments in the libraries was 330 bp. Libraries were pooled at equimolar concentrations then the pool was quantitated using the KAPA library quantification kit (KAPA Biosystems). Libraries were sequenced single end 75 base pairs using NextSeq500 (Illumina) at the Biopolymer’s Facility (Harvard School). For data analysis, RNAseq reads were aligned against GRCh38 human reference using HISAT2 (Kim et al., 2015) in the bcbio framework and gene expression counts in reads per million (RPM) and standard deviations from four replicates were calculated and analyzed with custom MATLAB software. Gene expression data (RNA seq) were deposited in the GEO (Gene Expression Omnibus, https://www.ncbi.nlm.nih.gov/geo/, accession number: GSE127988).

#### Estimation of clinically accessible drug dose ranges

Clinically accessible drug dose ranges for vemurafenib, dabrafenib, cobimetinib and trametinib were calculated from published concentrations measured in the plasma of human subjects (see **Table S3** for values and references). We used reported mean and standard deviations of plasma concentrations in patients (ug/mL) to calculate the corresponding micro molar (µM) concentrations from molecular weight of drugs (g/mol). Lower and upper bounds were defined as the lowest and the highest deviation from the mean minus or plus one standard deviation.

#### Model construction and definitions

MARM1 was built using PySB (Lopez et al., 2013) extended to include support for energy-based BNG (Sekar et al., 2017). The main model features are explained in the main text and a detailed description of model construction is provided in **Text S1**.

#### Parameter estimation and model simulations

The PySB generated model was compiled as C++ executable using the AMICI (Fröhlich et al., 2016) function *pysb2amici* (https://doi.org/10.5281/zenodo.2600696). For model simulation we used an absolute and relative integration tolerance of 10^-12^. Drug pretreatment for 24 hr was implemented as a steady-state constraint. Steady-states were computed using a simulation-based approach, in which the model was simulated until the time derivative was numerically equal to zero with absolute and relative tolerance 10^-8^. Parameter estimation was performed using the *pypesto* toolbox (Stapor et al., 2018) (https://doi.org/10.5281/zenodo.2592733) assuming a weighted least-squares objective function. Residual weights were derived from experimentally observed standard deviations assuming a Gaussian error model. For parameter estimation purposes, standard deviations for immunofluorescence measurements were set at 0.2 in normalized fluorescence unit. To solve the respective optimization problem, we used the least-squares trust-region-reflective algorithm implemented in scipy (version 1.1.0) using an optimization variable tolerance of 10^-12^ and gradient tolerance of 10^-8^ as termination criteria. Residual sensitivities were computed using forward sensitivity analysis with an absolute and relative integration tolerance of 10^-8^ for state sensitivities. To avoid local minima, we used a multi-start strategy by repeatedly solving the optimization problem with different initial parameter values. The repeated optimization was parallelized across 100 cores using *snakemake* (Koster and Rahmann, 2012). Each core sequentially performed up to 10 optimization runs with a total computational budget of 5 days. On average, each core performed three complete optimization runs plus one unfinished optimization run, totaling in 402 estimated parameter vectors. For all subsequent model analysis, the single best or top 100 best performing parameter vectors were used, as specified in the main text. In the later, simulations uncertainty was quantified as quartile (2^nd^ and 3^rd^ quartile) ranges across the 100 simulations. Model simulations were exported and visualization scripts were written in MATLAB 2017b.

## Data and Software Availability

Gene expression data (RNA seq) were deposited in the GEO (Gene Expression Omnibus, https://www.ncbi.nlm.nih.gov/geo/, accession number: GSE127988). The SRM proteomics data were deposited in the Panorama public database (https://panoramaweb.org/fWeE7i.url). All experimental and simulation datasets used in the main and supplementary figures are described in **Table S4** and provided in the Synapse database (https://www.synapse.org/#!Synapse:syn20551877/files/, Synapse ID: syn20551877, DOI: 10.7303/syn20551877). The MARM1 computational model is provided in SBML (Hucka et al., 2018), BNG (Sekar et al., 2017) and PySB (Lopez et al., 2013) formats at Github (https://github.com/labsyspharm/marm1-supplement). Jupyter Notebooks for the step-by-step construction and simulation of MARM1 in PySB format are provided in a Docker container at Github (https://github.com/labsyspharm/marm1-supplement).

