## Supplemental Figures and Movie Legend for "Sporadic ERK pulses drive non-genetic resistance in drug-adapted BRAF^V600E^ melanoma cells"

#### SUPPLEMENTAL FIGURES AND MOVIES LEGENDS

### Figure S1

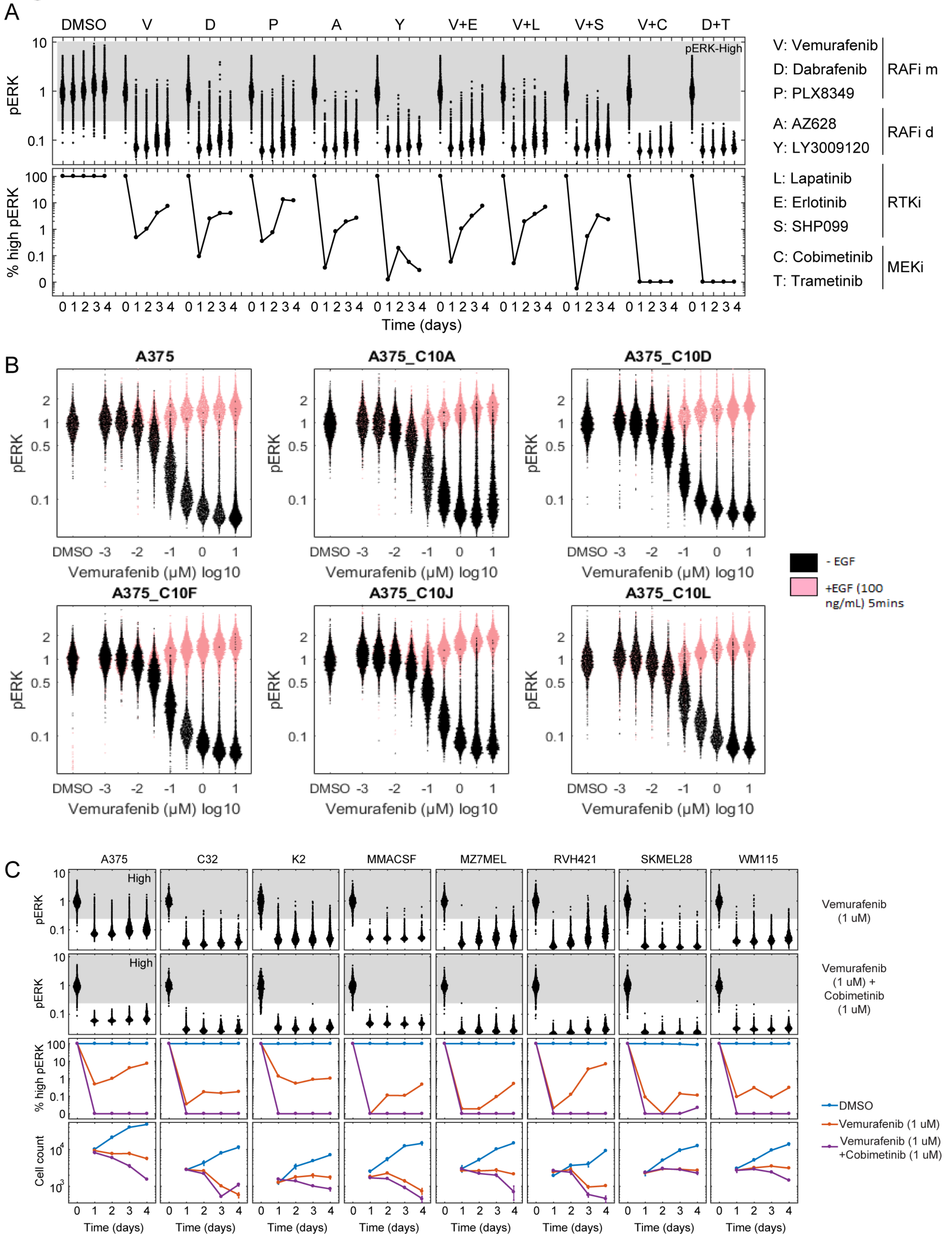

**Figure S1. Single-cell ERK reactivation in BRAF$^{V600E}$ melanoma cell lines treated with various RAF, MEK and RTK targeted inhibitors. Related to Figure 1. A)** Quantification of ERK phosphorylation in single cells (first row) by immunofluorescence and fraction of cells with high ERK phosphorylation (second row) in A375 cells treated for 4 days with single drugs or drug combinations as described in the legend (drugs were all dosed at 1 μM, except SHP099 at 5 μM). 'RAFi m': inhibitor of RAF monomers; 'RAFi d': inhibitor of RAF dimers. **B)** Single-cell ERK phosphorylation levels of A375 cells and five A375-derived clonal cell lines (10A, 10D, 10F, 10J, 10L) treated for 24 hr with vemurafenib in the absence of EGF (black dots) or with addition of 100 ng/mL EGF for 5 min (pink dots), as determined by immunofluorescence. Clonal cell populations reproduce adaptive ERK-high rare single-cell reactivation observed in the parental A375 cell line and respond homogenously with high ERK reactivation to EGF stimulation. **C)** Quantification of ERK phosphorylation in single cells (first and second rows), fraction of cells with high ERK phosphorylation (third row) and cell count (fourth row) for 8 BRAF$^{V600E}$ melanoma cell lines treated for four days with DMSO, vemurafenib (1 μM ), or vemurafenib (1 μM) plus cobimetinib (1 μM ).

### Figure S2

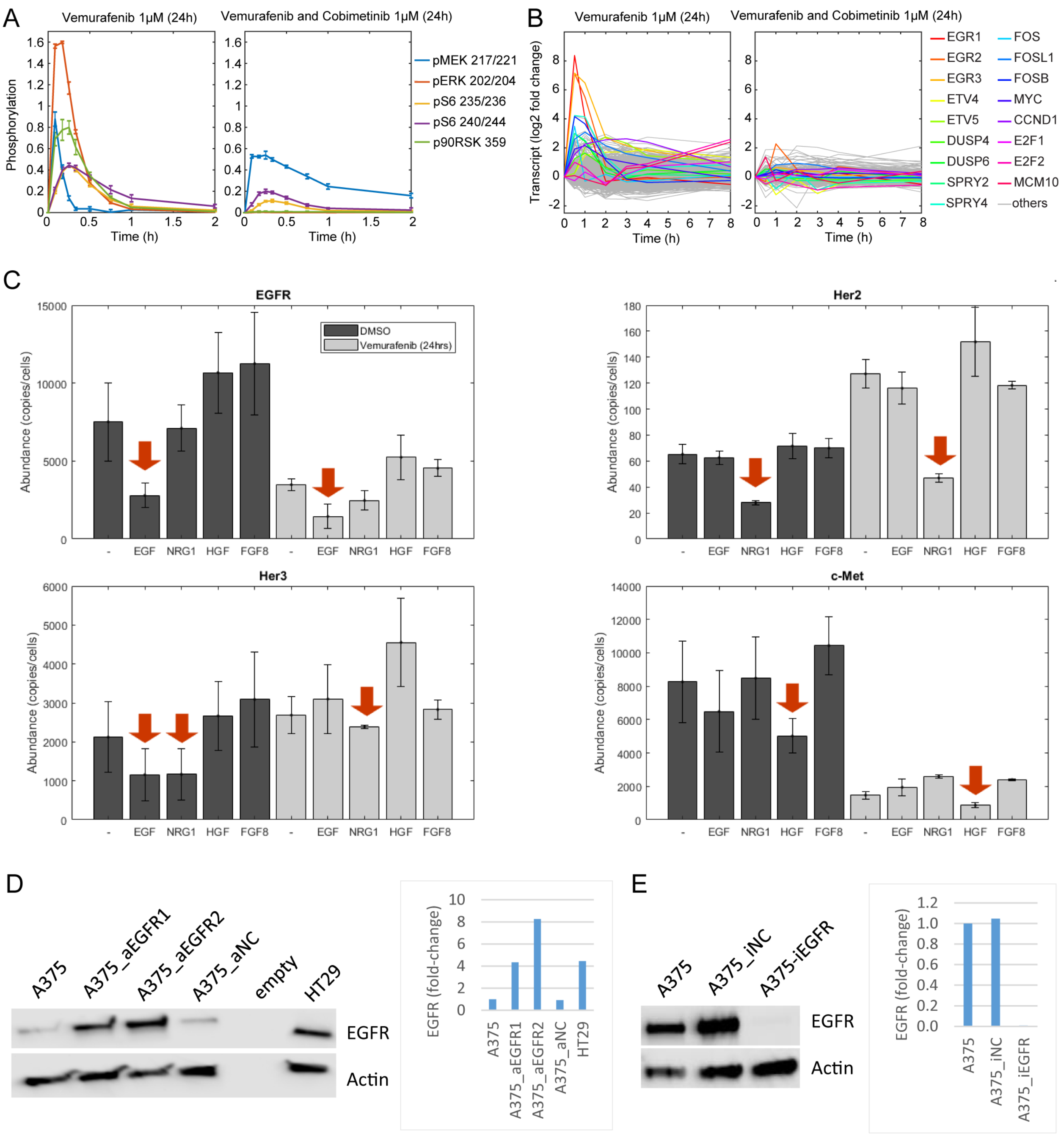


**Figure S2. Quantification of receptor abundance in A375, EGFR-upregulated A375, and EGFR-downregulated A375 cells and response to ligand stimulation in those cell lines. Related to Figure 2 and Figure 6. A)** Quantification of the phosphorylation levels of key MAPK and downstream signaling proteins on activating sites in A375 cells over the course of 2 h following exposure to vemurafenib (1 μM) or vemurafenib plus cobimetinib (1 μM each) for 24 hr and addition of 100 ng/mL EGF, as determined by immunofluorescence. The displayed are averages and standard deviations from three replicates. **B)** Transcript levels of key MAPK and downstream signaling proteins in A375 cells exposed to vemurafenib (1 μM) alone or with cobimetinib (1 μM) for 24 hr and then to 100 ng/mL EGF, as determined by transcriptomics. The displayed data is the average of 4 replicates. **C)** Absolute receptor abundances determined using a calibrated ELISA assay targeting EGFR, Her2, Her3 and c-Met in A375 cells treated for 24 hr with DMSO or 1 μM vemurafenib and then stimulated for 24 hr with the growth factors indicated on the x-axis. Arrows highlight receptors whose abundance is substantially decreased upon ligand addition; '–': no ligand addition). **D)** Quantification of EGFR protein levels by Western Blotting in A375, A375 CRISPRa cells expressing an sgRNA targeting EGFR (2 independent sgRNA: '_aEGFR1' and '_aEGFR2'), A375 CRISPRa cells expressing a non-targeting sgRNA guide ('_aNC'), and HT29 cells. **E)** Quantification of EGFR protein levels by Western Blotting in A375, A375 CRISPRi cells expressing an sgRNA targeting EGFR ('-iEGFR') and A375 cells carrying a CRISPRi non-targeting guide ('_iNC').

### Figure S3

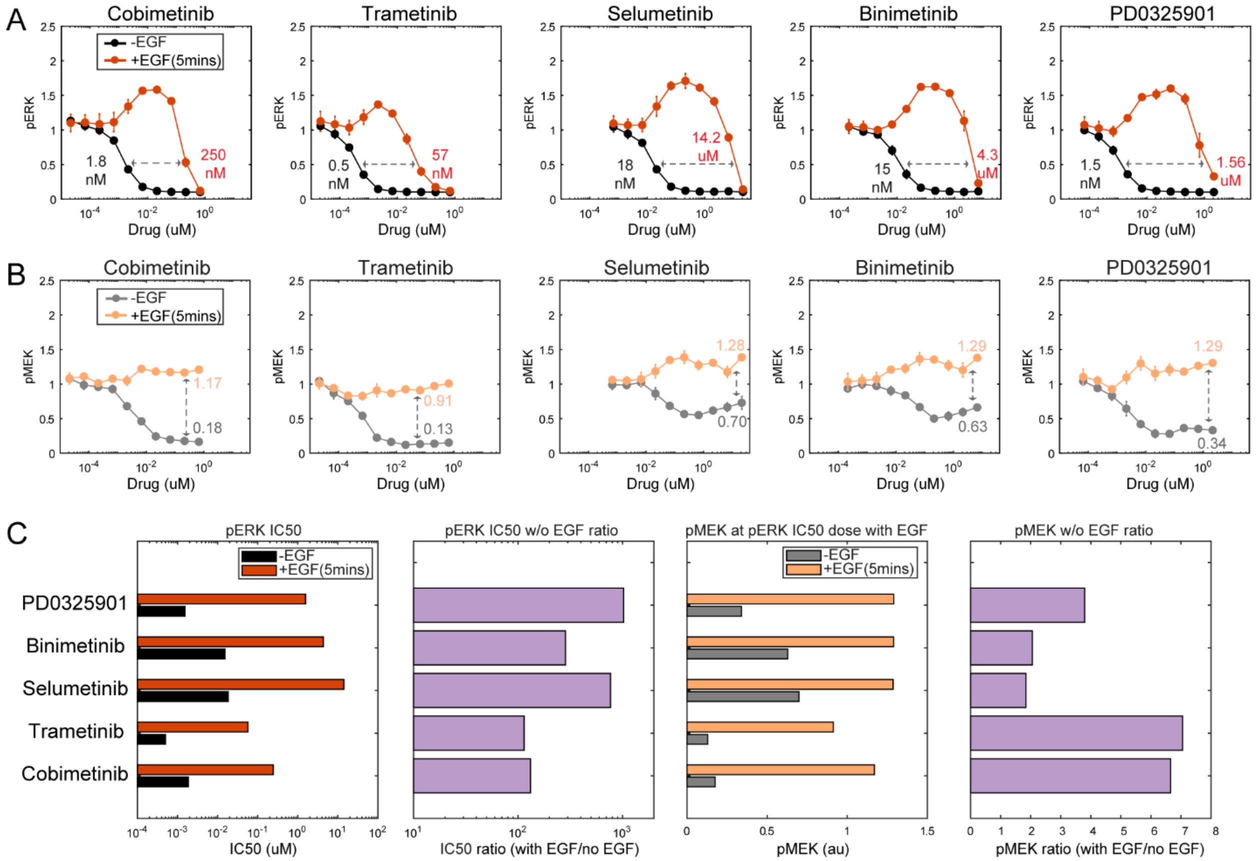


**Figure S3. ERK and MEK phosphorylation levels in A375 cells treated with various MEK inhibitors with or without EGF stimulation. Related to Figure 4. A-B)** ERK (A) and MEK (B) phosphorylation levels in A375 cells treated for 24 hr with vemurafenib, cobimetinib, selumetinib, binimetinib, or PD0325901 with or without stimulation for 5 min with 100 ng/mL EGF, as determined by immunofluorescence. $EC_{50}$ values for ERK phosphorylation with or without EGF stimulation for each inhibitor are shown in (A). Levels of MEK phosphorylation at the determined $EC_{50}$ with or without EGF stimulation are shown in (B). **C)** Bar graphs representing the quantification of data shown in (A) and (B).

### Figure S4

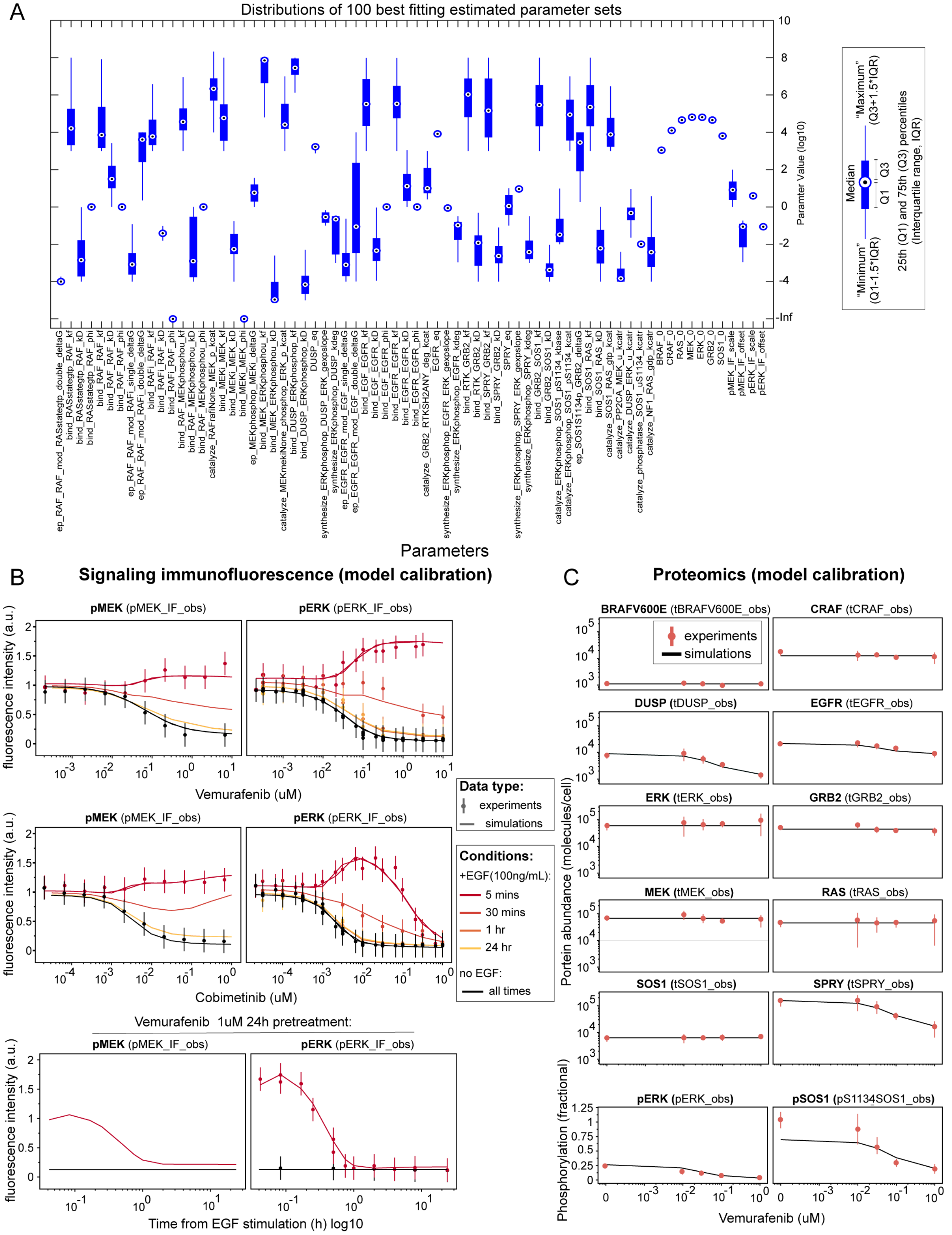

A
Distributions of 100 best fitting estimated parameter sets
Paramter Value (log10)
Parameters
Median
"Maximum" (Q3+1.5*IQR)
"Minimum" (Q1-1.5*IQR)
Q1
Q3
25th (Q1) and 75th (Q3) percentiles (Interquartile range, IQR)
B
Signaling immunofluorescence (model calibration)
pMEK (pMEK_IF_obs)
pERK (pERK_IF_obs)
fluorescence intensity (a.u.)
Vemurafenib (uM)
Cobimetinib (uM)
Vemurafenib 1uM 24h pretreatment:
Time from EGF stimulation (h) log10
Data type:
experiments
simulations
Conditions:
+EGF(100ng/mL):
5 mins
30 mins
1 hr
24 hr
no EGF:
all times
C
Proteomics (model calibration)
BRAFV600E (tBRAFV600E_obs)
CRAF (tCRAF_obs)
DUSP (tDUSP_obs)
EGFR (tEGFR_obs)
ERK (tERK_obs)
GRB2 (tGRB2_obs)
MEK (tMEK_obs)
RAS (tRAS_obs)
SOS1 (tSOS1_obs)
SPRY (tSPRY_obs)
pERK (pERK_obs)
pSOS1 (pS1134SOS1_obs)
experiments
simulations
Portein abundance (molecules/cell)
Phosphorylation (fractional)
Vemurafenib (uM)

**Figure S4. Data fitting and parameter estimation for the computational model MARM1. Related to Figure 5. A)** Box-plot showing the median and interquartile distributions of model parameters for the 100 best performing estimated parameter sets. **B-C)** Model fitting of training data for the best parameter set. Training data include signaling dynamics for pERK and pMEK at various doses of vemurafenib and cobimetinib without or with addition of 100 ng/mL EGF across multiple time-points (**B**) and absolute (phospho)-proteomics data for cells treated with four doses of vemurafenib for 24 hr, as shown in Figure 3 **(C)**.

### Figure S5

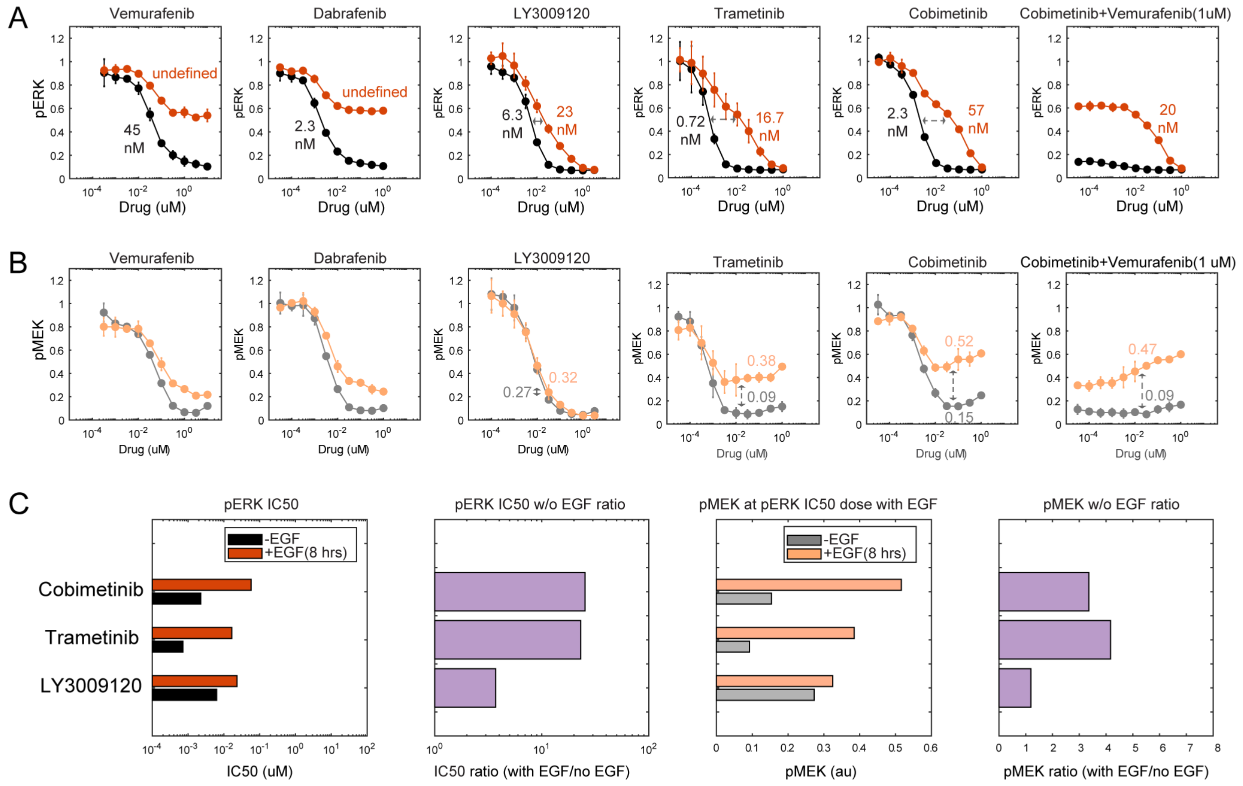


**Figure S5. ERK and MEK phosphorylation levels in EGFR-upregulated A375 cells treated with various RAF and MEK inhibitors with or without EGF stimulation. Related to Figure 6. A-B)** ERK (A) and MEK (B) phosphorylation levels in A375_aEGFR1 cells (EGFR-overexpressing) treated for 24 hr with vemurafenib, dabrafenib, LY3009120, trametinib, cobimetinib, or PD0325901 with or without stimulation for 8 hr with 100 ng/mL EGF. $EC_{50}$ values for ERK phosphorylation with or without EGF stimulation for each inhibitor are shown in (A). Levels of MEK phosphorylation at the determined $EC_{50}$ with or without EGF stimulation are shown in (B). **C)** Bar graphs representing the quantification of data shown in (A) and (B).

### Figure S6

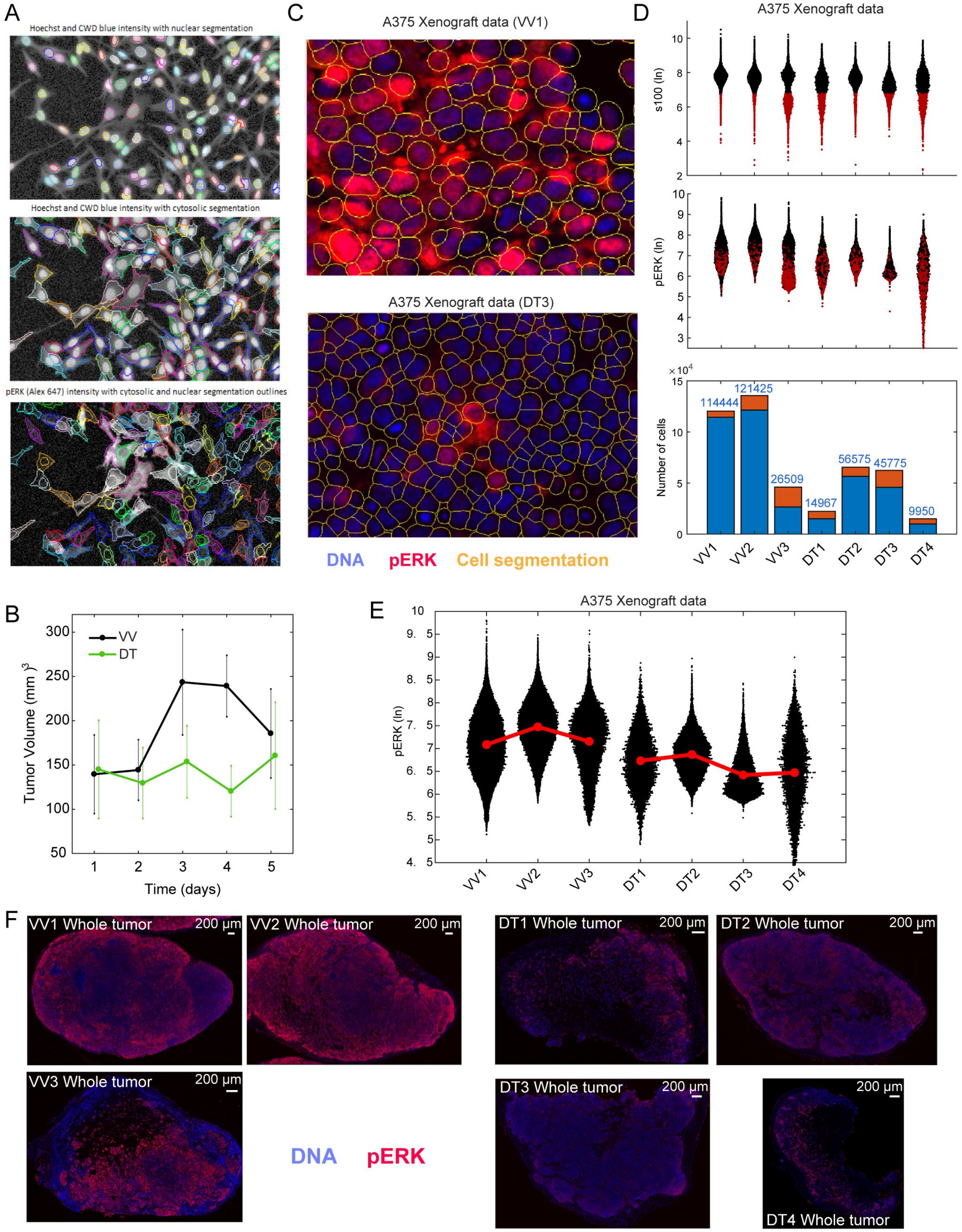
A
Hoechst and CWD blue intensity with nuclear segmentation
Hoechst and CWD blue intensity with cytosolic segmentation
pERK (Alex 647) intensity with cytosolic and nuclear segmentation outlines
C
A375 Xenograft data (VV1)
A375 Xenograft data (DT3)
DNA pERK Cell segmentation
D
A375 Xenograft data
s100 (ln)
pERK (ln)
Number of cells
114444
121425
26509
14967
56575
45775
9950
VV1 VV2 VV3 DT1 DT2 DT3 DT4
B
VV
DT
Tumor Volume (mm)^3
Time (days)
E
A375 Xenograft data
pERK (ln)
VV1 VV2 VV3 DT1 DT2 DT3 DT4
F
VV1 Whole tumor
VV2 Whole tumor
VV3 Whole tumor
DT1 Whole tumor
DT2 Whole tumor
DT3 Whole tumor
DT4 Whole tumor
200 µm
DNA pERK

**Figure S6. Quantification of single-cell ERK phosphorylation in A375 fixed cells and tumor xenografts. Related to Figure 7. A)** Example of segmentation used to quantify single-cell signaling in fixed cell lines. Data shown are segmentation of nuclei using Hoechst signal (upper panel), of cytosolic area using CWD staining (middle panel), and both segmentations overlaid with pERK staining in exemplary cells with pulsatile reactivation (bottom panel). **B)** Volume of A375 xenograft tumors treated with vehicle-vehicle (VV, N=3) or with dabrafenib plus trametinib (DT, N=4) over the course of 5 days of treatment. **C)** Example of segmentation used to quantify single-cell signaling in fixed tumors from xenografts. Data shown are immunofluorescence images of Hoechst and pERK staining of representative VV and DT samples overlaid with cytosolic segmentation. **D)** Quantification of pERK levels in single cells of A375 xenograft tumors. Segmented cells were gated to retain only S100-positive cells, a marker of melanoma, as identified visually based on the S100 distribution. S100-positive cells are displayed in black and S100-negative in red (first and second plots). The number of segmented cells that passed or not the S100 cutoff in each sample are shown in blue or orange, respectively (third plot). **E)** Single-cell distribution of ERK phosphorylation levels determined at 5 days post-injection in VV and DT treated A375 tumor xenografts. Red dots indicates average pERK levels and samples from the same groups are connected by a red line. **F)** Whole tumor images of all A375 xenografts used in this work.

**Movie S1. Related to Figure 7.** Movie of a representative single field of view (one out of 10 recorded) of A375 ERK:KTR fluorescent reporter cells treated with 1 µM vemurafenib and imaged over the course of 15 hr at 6 min resolution starting at 24 hr post-drug addition. *Upper left panel:* ERK:KTR-CFP raw intensities. *Upper right panel:* Nuclear masks of segmented cells color-coded accordingly to their quantified ERK activity (determined by the cytosolic CFP to nuclear CFP ratio). Cells for which the automatic tracking software was able to reconstruct a tracking sequence from start to end of the movie are labeled with their identifier number in black. One representative cell is highlighted in red in all panels (ID=8). *Lower left panel:* ERK activity of all cells continuously tracked for the entire duration of the movie. *Lower right panel:* Cumulative distribution of ERK activities for cells tracked for the entire duration of the movie.
