## Supplementary material for "Sporadic ERK pulses drive non-genetic resistance in drug-adapted BRAF^V600E^ melanoma cells": Table S1

**Table S1. Related to Figure 3. Abbreviations and HUGO names of proteins in the MAPK pathway.**

| Name Abbreviation | HUGO Name | Full name | HGNC ID |
|---|---|---|---|
| EGFR | EGFR | epidermal growth factor receptor | HGNC:3236 |
| MIG6 | ERRFI1 | ERBB receptor feedback inhibitor 1 | HGNC:18185 |
| SPRY4 | SPRY4 | sprouty RTK signaling antagonist 4 | HGNC:15533 |
| SPRY2 | SPRY2 | sprouty RTK signaling antagonist 2 | HGNC:11270 |
| GRB2 | GRB2 | growth factor receptor bound protein 2 | HGNC:4566 |
| CBL | CBL | Cbl proto-oncogene | HGNC:1541 |
| SHC1 | SHC1 | SHC adaptor protein 1 | HGNC:10840 |
| SHP2 | PTPN11 | protein tyrosine phosphatase non-receptor type 11 | HGNC:9644 |
| SOS1 | SOS1 | SOS Ras/Rac guanine nucleotide exchange factor 1 | HGNC:11187 |
| HRAS | HRAS | HRas proto-oncogene, GTPase | HGNC:5173 |
| KRAS | KRAS | KRAS proto-oncogene, GTPase | HGNC:6407 |
| NRAS | NRAS | NRAS proto-oncogene, GTPase | HGNC:7989 |
| ARAF | ARAF | A-Raf proto-oncogene, serine/threonine kinase | HGNC:646 |
| BRAF | BRAF | B-Raf proto-oncogene, serine/threonine kinase | HGNC:1097 |
| CRAF | RAF1 | Raf-1 proto-oncogene, serine/threonine kinase | HGNC:9829 |
| MEK1 | MAP2K1 | mitogen-activated protein kinase kinase 1 | HGNC:6840 |
| MEK2 | MAP2K2 | mitogen-activated protein kinase kinase 2 | HGNC:6842 |
| ERK1 | MAPK3 | mitogen-activated protein kinase 3 | HGNC:6877 |
| ERK2 | MAPK1 | mitogen-activated protein kinase 1 | HGNC:6871 |
| DUSP4 | DUSP4 | dual specificity phosphatase 4 | HGNC:3070 |
| DUSP6 | DUSP6 | dual specificity phosphatase 6 | HGNC:3072 |
