## Supplementary material for "Sporadic ERK pulses drive non-genetic resistance in drug-adapted BRAF^V600E^ melanoma cells": Table S2

**Table S2. Relate to Figure 2. List of oligonucleotides used for CRISPR cell lines.**

| Cell line name (manuscript) | Cell line name (internal & datasets) | sgRNA protospacer | Target | Oligonucleotides used for sgRNA construction | Our oligo name |
|---|---|---|---|---|---|
| A375_iEGFR | A375_Cri_1/9 | GCCGCGCGAGCTAGACGTCC | EGFR knockdown | TTGGCCGCGCGAGCTAGACGTCCGTTTAAGAGC | GS_LSP_001 |
| | | | | TTAGCTCTTAAACGGACGTCTAGCTCGCGCGGCCAACAAG | GS_LSP_009 |
| A375_aEGFR1 | A375_Cra_F7_5/13 | GAACCAGCAGCGGGGACCCA | EGFR overexpression | TTGGAACCAGCAGCGGGGACCCAGTTTAAGAGC | GS_LSP_005 |
| | | | | TTAGCTCTTAAACTGGGTCCCCGCTGCTGGTTCCAACAAG | GS_LSP_013 |
| A375_aEGFR2 | A375_Cra_F7_6/14 | GCCCCGCGGGACCTAGTCTC | EGFR overexpression | TTGGCCCCGCGGGACCTAGTCTCGTTTAAGAGC | GS_LSP_006 |
| | | | | TTAGCTCTTAAACGAGACTAGGTCCCGCGGGGCCAACAAG | GS_LSP_014 |
| A375_iNC | A375_Cri_NC_79/80 | GCTGCATGGGGCGCGAATCA | knockdown non targeting control | TTGGCTGCATGGGGCGCGAATCAGTTTAAGAGC | CC_LSP_079 |
| | | | | TTAGCTCTTAAACTGATTCGCGCCCCATGCAGCCAACAAG | CC_LSP_080 |
| A375_iST3GAL4 | | GGGCGCGGGTCCGGCCTGGGAG | ST3GAL4 knockdown | TTGGGGCGCGGGTCCGGCCTGGGAGGTTTAAGAGC | CC_LSP_001 |
| | | | | TTAGCTCTTAAACCTCCCAGGCCGGACCCGCGCCCCAACAAG | CC_LSP_002 |
| A375_iSEL1L | | GCAGGAAGAGCAGCGGCGAGG | SEL1L knockdown | TTGGCAGGAAGAGCAGCGGCGAGGGTTTAAGAGC | CC_LSP_003 |
| | | | | TTAGCTCTTAAACCCTCGCCGCTGCTCTTCCTGCCAACAAG | CC_LSP_004 |
| A375_iDPH1 | | GTATCCGGGGCAGCGGAGCA | DPH1 knockdown | TTGGTATCCGGGGCAGCGGAGCAGTTTAAGAGC | CC_LSP_005 |
| | | | | TTAGCTCTTAAACTGCTCCGCTGCCCCGGATACCAACAAG | CC_LSP_006 |
| A375_aNC | A375_Cra_NC_111/112 | GTGTCGTGATGCGTAGACGG | overexpression non targeting control | TTGGTGTCGTGATGCGTAGACGGGTTTAAGAGC | CC_LSP_111 |
| | | | | TTAGCTCTTAAACCCGTCTACGCATCACGACACCAACAAG | CC_LSP_112 |
| A375_aCDKN1C | | GCCGGGGCGCGCGGCTGATTGG | CDKN1C overexpression | TTGGCCGGGGCGCGCGGCTGATTGGGTTTAAGAGC | CC_LSP_007 |
| | | | | TTAGCTCTTAAACCCAATCAGCCGCGCGCCCCGGCCAACAAG | CC_LSP_008 |
| A375_aSLC4A1 | | GTCAGGAGAACCATGGGGACC | SLC4A1 overexpression | TTGGTCAGGAGAACCATGGGGACCGTTTAAGAGC | CC_LSP_009 |
| | | | | TTAGCTCTTAAACGGTCCCCATGGTTCTCCTGACCAACAAG | CC_LSP_010 |
| A375_aPOU5F1 | | GGATGTTTGCCTAATGGTGG | POU5F1 overexpression | TTGGGATGTTTGCCTAATGGTGGGTTTAAGAGC | CC_LSP_011 |
| | | | | TTAGCTCTTAAACCCACCATTAGGCAAACATCCCAACAAG | CC_LSP_012 |
