## Supplementary material for "Sporadic ERK pulses drive non-genetic resistance in drug-adapted BRAF^V600E^ melanoma cells": Table S3

**Table S3. Related to Figure 1, Figure 4, Figure 5 and Figure 6. Clinically accessible drug dose ranges for vemurafenib, dabrafenib, cobimetinib and trametinib.** Clinical accessible drug dose ranges were calculated using upper and lower bounds in plasma concentrations of human subjects treated with the dosage regimen as described in the table. Plasma concentrations in µg/mL were converted to µM using the drug's molecular weight (g/moL). In cases in which primary measurements were given as mean and standard deviations, upper and lower bounds were calculated as the mean minus or plus one standard deviation.

| Drug | Molecular Weight (g/moL) | Lower bound (µg/mL) | Upper bound (µg/mL) | Lower bound (µM) | Upper bound (µM) | N patients | Dosage regimen | Reference |
|---|---|---|---|---|---|---|---|---|
| Vemurafenib | 489.922 | 1.46 | 84.16 | 2.98 | 171.78 | 23 | 960 mg/day (four 240-mg tablets) orally taken every 12 hours. Measured at 1 day and 15 day after starting treatment. | PMID: 25899783 (Table 2) |
| Dabrafenib | 519.56 | 15.4 | 279.6 | 0.03 | 0.54 | 27 | 300 mg/day (two 75-mg tablets) orally take twice daily. Measured 15 days after starting treatment. | PMID: 28709799 (Table 1) |
| Cobimetinib | 531.3 | 24 | 344 | 0.05 | 0.65 | 19 | 60 mg/day (three 20-mg tablets) orally taken once daily. Measured 8 hours after starting treatment. | Clinical Pharmacology NDA Review (NDA 206192) Table 6 (Stage II) |
| Trametinib | 615.39 | 4.1 | 32.9 | 0.0067 | 0.053 | 27 | 2 mg/day (one 2 mg tablet) orally taken once daily. Measured 15 days after starting treatment. | PMID: 28709799 (Table 1) |
