## Supplementary material for "Sporadic ERK pulses drive non-genetic resistance in drug-adapted BRAF^V600E^ melanoma cells": Table S4

**Table S4. Related to Figure 1-7 and S1-S6. List of datasets used in this work and their appearance in main and supplementary figures.** List of datasets used in this work and their use in figures. IF: population-mean immunofluorescence microscopy, sc-IF: single-cell immunofluorescence microscopy, RNAseq: transcriptomics by RNAseq, Prot: absolute abundance and phospho- proteomics, LC: live-cell imaging, Sim: model simulations, Others: other data type as explained in the description. Replication (T) technical or (B) biological.

| Dataset file | Type | Description | N replicates | Used in.. |
|---|---|---|---|---|
| D1.csv | IF | Time-course pERK and pMEK in A375 cells pretreated for 24h with vemurafenib and EGF stimulated | **3 T** | **Fig. 2B** |
| D2.csv | IF | Time-course pERK in A375 cells pretreated for 24h with vemurafenib and NRG1, FGF8 and HGF stimulated | **1 T** | **Fig. 2B** |
| D3.csv | IF | Dose-response pERK in A375 cells pretreated for 24h with vemurafenib and/or cobimetinib and stimulated for 5 minutes with EGF, NRG1, FGF8 and HGF | **2 T** | **Fig. 2B, 2E, 5D** |
| D4.csv | IF | Dose-response pERK and pMEK in A375 cells pretreated for 24h with single and combined RAFi and MEKi and then stimulated or not for 5 min with EGF | **2 T** | **Fig. 4A, 4B, 5B, S3** |
| D5.csv | IF | Time-course pERK in A375 cells treated for a variable amount of time with vemurafenib and then stimulated at different times with EGF, NRG1, FGF8 and HGF | **1 T** | **Fig. 2A** |
| D6.csv | IF | Time-course pERK in A375 parental, iEGFR, aEGFR1 and aEGFR2 pretreated for 24 h with vemurafenib and then stimulated with EGF | **1 T** | **Fig. 2F** |
| D7.csv | IF | Dose-response pERK and pMEK in A375 cells aEGFR1 cells pretreated for 24h with single and combined RAFi and MEKi and then stimulated or not for 8h with EGF | **2T** | **Fig. 6E, 6F, S5** |
| D8.csv | IF | Dose- and time- course pERK in A375 cells pretreated with vemurafenib and or cobimetinib for 24h and then stimulated for different times with EGF | **2T** | **Fig. S4B** |
| D9.csv | IF | Dose-response pERK of A375 parental, aEGFR1, aEGFR2 and colorectal HT29 cells pretreated for 24h with vemurafenib and/or cobimetinib and then stimulated for 8 h with EGF | **2T** | **Fig. 6F** |

| | | | | |
|---|---|---|---|---|
| D10.csv | IF | Time-course pMEK, pERK, pS6, p90RKS in A375 cells pretreated for 24 h with vemurafenib alone or with cobimetinib and then stimulated with EGF | **3T** | **Fig. 2C, S2B** |
| ME.csv | IF | Dose-response matrixes of pERK for A375 and A375 aEGFR2 cells treated with vemurafenib and cobimetinib and then stimulated or not with EGF | **1T** | **Fig. 6A, 6B, 6C** |
| G1.csv | sc-IF | Time-course single-cell quantification of Edu, pERK and DNA content in A375 cells treated over 4 days with Vemurafenib alone or with different Cobimetinib concentrations | **2 T** | **Fig. 1A, 1C** |
| C1.csv | sc-IF | Time-course cell count and single-cell pERK for 8 melanoma cell lines treated over 4 days with Vemurafenib alone or with Cobimentinib | **2 T** | **Fig. 1D, S1C** |
| C2.csv | sc-IF | Time-course single-cell pERK for A375 cells treated over 4 days with different RAFi and/or MEKi and/or RTKi | **2 T** | **Fig. S1A** |
| C3.csv | sc-IF | Dose-response single-cell pERK for parental and clonal A375 cells treated with vemurafenib and trametinib | **2 T** | **Fig. 4C, 4F, S1B** |
| C4.csv | sc-IF | Time-course single-cell pERK in A37 cells treated over 10 h with vemurafenib | **2 T** | **Fig. 1B** |
| C5.csv | sc-IF | Single-cell EGFR abundance at the membrane for A375 parental and aEGFR2 cells treated for 24h with vemurafenib | **3 T** | **Fig. 6H** |
| C6.csv | sc-IF | Single-cell pERK in A375 A375 parental, aEGFR2, iEGFR and aCrisprControl cells tratead for 24h with vemurafenib and then stimulated for variable times and doses of EGF | **1 T** | **Fig. 6G** |
| DP1.csv | Prot | Absolute abundance of proteins in A375 cells treated for 24 h with different doses of vemurafenib | **4 T** | **Fig. 3B, 3D, S4C** |
| DP2.csv | Prot | Absolute phosphorylation of proteins in A375 cells treated for 24 h with different doses of vemurafenib | **4T** | **Fig. 3E, S4C** |
| DT1.csv | RNAseq | Transcript levels in A375 cells treated with different doses of Vemurafenib | **2T x 2B** | **Fig. 4C** |
| DT2.csv | RNAseq | Transcript levels time-course in A375 cells stimulate with EGF after 24 h pretreatment with Vemurafenib | **2T x 2B** | **Fig. 2D, S2B** |
| DT3.csv | RNAseq | Transcript levels time-course in A375 cells stimulate with EGF after 24 h pretreatment with Vemurafenib and Cobimetinib | **2T x 2B** | **Fig. S2B** |

| | | | | |
|---|---|---|---|---|
| LC1.csv | LC | Live-cell traces of A375 ERK reporter cells with successful automatic tracking over 16 hours of measurements in DMSO, Vemurafenib alone or with Cobimentinib. | **10 T** | **Fig. 1F, 1G, 1H, 1I, 7A** |
| LC2.csv | LC | Live-cell traces of A375 ERK reporter cells identified but with gaps in automatic tracking over 16 hours of measurements in DMSO, Vemurafenib alone or with Cobimentinib. | **10 T** | **Fig. 7A** |
| PS.csv | Sim | 100 best fit estimated parameter sets for the model | **-** | **Fig. S4A** |
| MS.csv | Sim | Simulations of multiples species for A375 parental and EGFR overexpressed cells treated with single-agent and combined vemurafenib and cobimetinib for 24 h and then stimulated or not with EGF | **-** | **Fig. 5A, 6A, 6B, 6C, 6D** |
| DX.csv | Others | Absolute abundance of receptors EGFR, Her2, Her3 and c-Met for A375 cells treated with Vemurafenib and stimulated with EGF, NRG1, FGF8, HGF | **3 T** | **Fig. S2C** |
| XSC.csv | Others | Single-cell quantification of pERK, S100 and Hoechest (DNA) in A375 mouse xenografts untreated or treated with dabrafenib and trametinib | **3-4 B** | **Fig. 7C, S6C-F** |
| XTV.csv | Others | Tumor volumes in A375 mouse xenografts untreated or treated with dabrafenib and trametinib | **3-4B** | **Fig. S6B** |
| DD.csv | Others | Plasma concentrations ranges for vemurafenib, cobimetinib, dabrafenib and trametinib as measured in patients from published works | **-** | **Fig. 1C, 4, 5B, 6A-F** |
| QP.csv | Others | QPCR validation of CRISPRa/I constructs using canonical control genes | **-** | **-** |
