## Supplementary material for "Sporadic ERK pulses drive non-genetic resistance in drug-adapted BRAF^V600E^ melanoma cells": Text S1

**Text S1. Related to Figure 5, Figure 6 and Figure S4.** Documentation for the model of MAPK signaling in $BRAF^{V600E}$ melanoma A375 cells (MARM1).

**Model code availability**

We provide MARM1 in SBML, BNG and PySB formats on Github (https://github.com/labsyspharm/marm1-supplement). In the same Github, we provide a Jupyter Notebook (*MARM1_construction.ipynb*) that reproduces the step-by-step construction of MARM1 in PySB, with a detailed description of the model assumptions and molecular mechanisms considered. We also provide a Jupyter Notebook for running model simulations (*MARM1_simulation.ipynb*). The Jupyter Notebooks are provided within a Docker container equipped with all the necessary software installations (e.g. energy-based BNG and PySB).

**MARM1: an overview**

The **Melanoma Adaptive Resistance Model 1.0 (MARM1)** was constructed using the rule-based modeling framework PySB (Lopez et al., 2013) extended to support energy-based BNG (Sekar et al., 2017). The support for energy BNG allows to write rule-based models with cooperativity that satisfy detailed balance by construction. In not energy-based (classic) rule-based modelling, detailed balance needs to be enforced manually when specifying rate laws, which can be cumbersome and error-prone for large models such as MARM1. Since many mechanisms of physiological MAPK cascade activation (e.g. RAF dimerization induced by Ras-GTP binding) and drug resistance (e.g. lower affinity of RAF inhibitors for RAF dimers) are based on molecular cooperativity (Kholodenko, 2015)(Rukhlenko et al., 2018), we use the support for energy-based BNG as a principled and convenient way to model cooperativity. A detailed description of how PySB was extended to support energy-based BNG and how energy-based rules are used is provided in following sections. Here, we describe the basic features of MARM1.

**MARM1 is a model of MAPK signaling in A375 melanoma cells**, which are homozygous for the $BRAF^{V600E}$ mutation and are a widely used cell line for the study of $BRAF^{V600E}$ melanoma. The model describes protein and drug species as intracellular concentrations in uM at a cell volume of 1pL = $10^{-12}$L. The unit of time is hours. The model includes 13 proteins, also termed 'basic species', which can form protein complexes with, or can be modified by, other species. The **13 basic species** contain in total **26 binding sites** that mediate their binding and **5 state variables** that represent modification states (either by phosphorylation or GDP/GTP loading). Binding and modification events generate **1007 complex species**, which is thus the number of Ordinary Differential Equations (ODEs) used in the model to simulate their changes over time. The model uses **55 rules** (**12 energy-based** and **43 non energy-based**, or classic, **rules**) to define the underlying biochemical network. The automatically generated biochemical network consists of **13164 reactions** that determine the rates of change of complex species over time. To model cooperativity, we used **25 energy patterns**, which assign energy values to patterns in species complexes, thus allowing the automatic calculation of the free energies of formation for each reactant species in each reaction. Free energies of formation of reactants are then used to calculate forward and backward reaction rates according to linear transition state theory (Sekar et al., 2017). The model contains **81 parameters**. **70 parameters were estimated by fitting the model to data** through optimization and we provide the **100 best-fitting parameter sets**. **7 parameters were manually set** since they represent modelling choices on the contribution of forward and backward rates for energy-based rate changes. There are **4 input parameters** used to initialize (i.e. set the pre-treatment condition) and perturb the model (i.e. set a change after pre-treatment): the concentration of the EGF ligand, the concentration of the RAF and MEK inhibitors, and the magnitude of EGFR overexpression or downregulation simulating CRISPRa or CRISPRi perturbations. We defined **10 observables** to extract the activity of key basic or complex species from simulations. These observables allow to sum up the contribution of multiple species to quantify overall activity of biological components (e.g. the sum of all active RAFs that are not bound by the RAF inhibitor as the total RAF catalytic activity).

##### Support for energy-based BNG in PySB

Energy-based BioNetGen (BNG) is a published extension of BNG (Sekar et al., 2017) to write models that 'satisfy detailed balance and cooperative interactions that are compactly specified as free energy contributions'. The PySB framework we used to build MARM1 defines models as a Python programs, which are in turn converted to another program's format to generate the underlying biochemical reaction network, in this case by using energy-based BNG. The resulting biochemical reaction network is imported in PySB for model analysis and simulations.

Energy-based BNG introduced two new syntactic elements to define the free energy of species: the *energy pattern* and the *energy rule*. In brief, 'an energy pattern is a pattern e*Pattern* that is assigned an energy value *EnergyP*' (Sekar et al., 2017), whit the following syntax:

> **begin energy patterns**
> *EnergyPatternLabel: ePattern EnergyP*
> **end energy patterns**

An energy rule is a BNG reaction rule is specified by a *rate distribution parameter* (ϕ) and an *energy of activation* (EA), instead of by reaction rates (i.e. forward and backward reaction rates). The keyword *Arrhenius* is used to associated the two parameters to an *energy rule label* and the reaction mechanism specified by its *substrates* (RP) and *products* (PP):

> **begin reaction rules**
> *EnergyRuleLabel: RP1 + .... +RPn <-> PP1+ ...+ PPm     Arrhenius(ϕ, EA)*
> **end reaction rules**

To support the generation of energy-BNG models from PySB, we introduced two corresponding syntactic elements, the *EnergyPattern* and the *EnergyRule* in PySB. In PySB an *EnergyPattern* has the following syntax:

*EnergyPattern ( EnergyPatternLabel, ePattern, EnergyP )*

And energy rules have the following syntax:

*Rule( EnergyRuleLabel , RP1 + .... +RPn | PP1+ ...+ PPm , ф , EA , energy=True)*

where the term *energy=True* distinguishes such rule from classic non-energy based rules. Because the two new syntaxes in both languages contains the same number and type of parameters, it is then straightforward to convert energy-based rules defined in PySB to the corresponding underlying energy-based BNG commands.

**Energy-based rules to model cooperativity**

We followed foundational work from *Kholodenko and colleagues* that demonstrated how to model activation mechanisms and drug resistance in the MAPK cascade by means of a thermodynamic derivation (Kholodenko, 2015)(Rukhlenko et al., 2018). We devised a formulation to impose the same type of cooperativity rates (Kholodenko, 2015) but expressed in terms of free energy using energy patterns and energy rules in energy-based BNG and PySB. Here, we describe how the formulation works using as an example cooperativity among RAFs and a RAF inhibitor. Following the original work (Kholodenko, 2015), we first describe the basal dimerization of RAFs and the binding of the RAF inhibitor to RAF. Note that in MARM1 we model both homo- and hetero-dimerization of BRAF and CRAF, but here for simplicity show a scenario with a single generic RAF specie. The two reactions are specified by the following PySB code:

```
# define monomers
Monomer('R', ['r', 'i'])
Monomer('I',['r'])
# define parameters
Parameter('kf_RR', 0.1)
Parameter('kb_RR', 1)
Parameter('RR_phi', 1)
Parameter('kf_RI', 0.1)
Parameter('kb_RI', 1)
Parameter('RI_phi', 0)
# define expressions
Expression('Gf_RR', log(kb_RR /kf_RR))
Expression('Gf_RI', log(kb_RI /kf_RI))
Expression('Ea0_RR', -(RR_phi* log(kb_RR/kf_RR) + log(kf_RR)))
Expression('Ea0_RI', -(RI_phi* log(kb_RI/kf_RI) + log(kf_RI)))
# define energy patterns
EnergyPattern('ep_RR', R(r=1) % R(r=1), Gf_RR)
EnergyPattern('ep_RI', R(i=1) % I(r=1), Gf_RI)
# define rules
```

```
Rule('RR_bind', R(r=None)+R(r=None)|R(r=1)%R(r=1),RR_phi, Ea0_RR, energy=True)
Rule('RI_bind', R(i=None)+I(r=None)|R(i=1)%I(r=1),RI_phi, Ea0_RI, energy=True)
```

This code defines the basic binding reactions between RAFs and between RAF and the RAF inhibitor, without any cooperativity. Key to the formulation is a parametrization in terms of forward and backward kinetic rates (*kf*, *kr*) that are converted to energy parameters by the use of expressions. For each energy reaction in MARM1, we convert its forward and backward rates (*kf* and *kb*) and distribution rate parameter *phi* into the free energy *Gf* and energy of activation *Ea0* according to the following formulas:

Gf = log(kb/kf)
Ea0 = -( phi * log(kb/kf) + log(kf))

The free energy *Gf* and energy of activation *Ea0* are then used as the parameters necessary to define the *EnergyPattern* and *Rule*, respectively. This formulation guarantees that biochemical network generated by energy-based BNG will be identical to the classical non-energy rule formulation that assumes rate independence (i.e. no cooperativity).

To impose cooperativity on the binding of RAFs depending on the RAF inhibitor binding, we define two additional *EnergyPattern* that modify the free energy of complex species containing patterns of single or double RAF inhibitor binding to RAF dimers:

```
# define parameters
Parameter('f', 0.01)
Parameter('g', 100)
#define expressions
Expression('Gf_IRR', log(f))
Expression('Gf_IRRI', log(f)+log(g))
# define expressions
EnergyPattern('ep_IRR', I(r=1) % R(r=2) % R(r=2, i=None), Gf_IRR)
EnergyPattern('ep_IRRI', I(r=1) % R(r=2) % R(r=2, i=3) % I(r=3), Gf_IRRI)
```

This code generates a biochemical network and rate laws that are identical to Kholodenko's original derivation in Figure 1A (Kholodenko, 2015). Thus, the parameters *f* and *g* represent the thermodynamic factors that modulate the propensity of RAF dimerization depending on the binding of one or two RAF inhibitors. This allows to model both paradoxical activation by RAF dimerization upon binding of the RAF inhibitors (f<1) and drug-resistance by lower affinity of one RAF protomer for the RAF inhibitor (g>1) as shown before (Kholodenko, 2015). In MARM1, we used the same formulation to impose cooperativity in many parts of the model, for example in the RAF dimerization-induced protection from RAF inhibitor binding, Ras-GTP induced RAF dimerization, EGF-induced EGFR dimerization and MEK inhibitor-induced reduction in affinity to phosphorylated MEK. Details are explained in the step-by-step construction code for MARM1 (https://github.com/labsyspharm/marm1-supplement, *MARM1_construction.ipynb*).

**Parameters estimated from data**

We list the 70 parameters that were estimated from data. We provide a description, their units, and the upper (UB) and lower (LB) bounds that constrained their parameter estimation. The estimated parameter sets are available in the supplementary dataset.

| | Parameter name | Description | Unit | UB (log10) | LB (log 10) |
|---|---|---|---|---|---|
| 1 | ep_RAF_RAF_mod _RASstategtp_double_deltaG | Thermodynamic factor that modifies the free energy of species containing a RAF dimer with both RAFs bound by Ras-GTP | no unit | 0 | -4 |
| 2 | bind_RASstategtp_RAF_kf | Forward rate of RasGTP binding to BRAF or CRAF | $\mu M^{-1} \cdot h^{-1}$ | 8 | 3 |
| 4 | bind_RASstategtp_RAF_kD | Dissociation constant for RasGTP and BRAF or CRAF | μM | 0 | -4 |
| 5 | bind_RAF_RAF_kf | Forward rate of BRAF and CRAF homo/hetero dimers | $\mu M^{-1} \cdot h^{-1}$ | 8 | 3 |
| 7 | bind_RAF_RAF_kD | Dissociation constant for BRAF and CRAF homo/hetero dimers | μM | 5 | 0 |
| 8 | ep_RAF_RAF_mod _RAFi_single_deltaG | Thermodynamic factor that modifies the free energy of species containing a RAF dimer with a single RAF bound by RAFi | no unit | 0 | -4 |
| 9 | ep_RAF_RAF_mod _RAFi_double_deltaG | Thermodynamic factor that modifies the free energy of species containing a RAF dimer with both RAFs bound by RAFi | no unit | 4 | 0 |
| 10 | bind_RAFi_RAF_kf | Forward rate of RAFi and BRAF or CRAF binding | $\mu M^{-1} \cdot h^{-1}$ | 8 | 3 |
| 12 | bind_RAFi_RAF_kD | Dissociation constant for RAFi and BRAF or CRAF binding | μM | 0 | -4 |
| 13 | bind_RAF_MEKphosphou_kf | Forward rate of BRAF or CRAF binding to unphosphorylated MEK | $\mu M^{-1} \cdot h^{-1}$ | 8 | 3 |
| 15 | bind_RAF_MEKphosphou_kD | Dissociation constant for BRAF or CRAF and unphosphorylated MEK binding | μM | 0 | -4 |

| | | | | | |
|---|---|---|---|---|---|
| 16 | catalyze_RAFrafiNone _MEK_p_kcat | Turnover number of MEK phosphorylation by an active RAF | $h^{-1}$ | 10 | 4 |
| 17 | bind_MEKi_MEK_kf | Forward rate of MEKi binding to unphosphorylated MEK | $\mu M^{-1} \cdot h^{-1}$ | 8 | 3 |
| 19 | bind_MEKi_MEK_kD | Dissociation constant for MEKi and unphosphorylated MEK binding | μM | 0 | -4 |
| 20 | ep_MEKphosphop_MEKi_deltaG | Thermodynamic factor that modifies the free energy of species containing a phosphorylated MEK protein bound by the MEK inhibitor | no unit | 4 | 0 |
| 21 | bind_MEK_ERKphosphou_kf | Forward rate of MEK binding to unphosphorylated ERK | $\mu M^{-1} \cdot h^{-1}$ | 8 | 3 |
| 22 | bind_MEK_ERKphosphou_kD | Dissociation constant for MEK and unphosphorylated ERK binding | μM | 0 | -5 |
| 23 | catalyze_MEKmekiNone _phosphop_ERK_p_kcat | Turnover number of ERK phosphorylation by an active MEK | $h^{-1}$ | 7 | 3 |
| 24 | bind_DUSP_ERKphosphop_kf | Forward rate of DUSP binding to phosphorylated ERK | $\mu M^{-1} \cdot h^{-1}$ | 8 | 3 |
| 25 | bind_DUSP_ERKphosphop_kD | Dissociation constant for DUSP and phosphorylated ERK binding | μM | 0 | -5 |
| 26 | DUSP_eq | Abundance of DUSP at equilibrium (production over degradation) | copies /cell | 3.5 | 1.5 |
| 27 | synthesize_ERKphosphop _DUSP_ERK_gexpslope | DUSP protein production rate catalyzed by active ERK | $h^{-1}$ | 2 | -1 |
| 28 | synthesize_ERKphosphop _DUSP_kdeg | DUSP degradation rate | $h^{-1}$ | -0.5 | -3 |
| 29 | ep_EGFR_EGFR_mod _EGF_single_deltaG | Thermodynamic factor that modifies the free energy of species containing a EGFR dimer with a single EGFR bound by EGF | no unit | 0 | -4 |
| 30 | ep_EGFR_EGFR_mod _EGF_double_deltaG | Thermodynamic factor that modifies the free energy of species containing a EGFR dimer with both EGFR bound by EGF | no unit | 4 | -4 |

| | | | | | |
|---|---|---|---|---|---|
| 31 | bind_EGF_EGFR_kf | Forward rate of EGF binding to EGFR | $\mu M^{-1} \cdot h^{-1}$ | 8 | 3 |
| 33 | bind_EGF_EGFR_kD | Dissociation constant for EGF and EGFR | μM | 0 | -4 |
| 34 | bind_EGFR_EGFR_kf | Forward rate of EGFR dimerization | $\mu M^{-1} \cdot h^{-1}$ | 8 | 3 |
| 36 | bind_EGFR_EGFR_kD | Dissociation constant for EGFR dimerization | μM | 4 | 0 |
| 37 | catalyze_GRB2 _RTKSH2ANY_deg_kcat | Degradation of EGFR dimers that are bound by GRB2 | $h^{-1}$ | 3 | 0 |
| 38 | EGFR_eq | Abundance of EGFR at equilibrium (production over degradation) | copies /cell | 5 | 3 |
| 39 | synthesize_ERKphosphop _EGFR_ERK_gexpslope | EGFR protein production rate catalyzed by active ERK | $h^{-1}$ | 2 | -1 |
| 40 | synthesize_ERKphosphop _EGFR_kdeg | EGFR degradation rate | $h^{-1}$ | -0.5 | -3 |
| 41 | bind_RTK_GRB2_kf | Forward rate of EGFR binding to GRB2 | $\mu M^{-1} \cdot h^{-1}$ | 8 | 3 |
| 42 | bind_RTK_GRB2_kD | Dissociation constant for EGFR binding to GRB2 | μM | 0 | -4 |
| 43 | bind_SPRY_GRB2_kf | Forward rate of SPRY binding to GRB2 | $\mu M^{-1} \cdot h^{-1}$ | 8 | 3 |
| 44 | bind_SPRY_GRB2_kD | Dissociation constant for SPRY binding to GRB2 | μM | 0 | -4 |
| 45 | SPRY_eq | Abundance of EGFR at equilibrium (production over degradation) | copies /cell | 1 | -1 |
| 46 | synthesize_ERKphosphop _SPRY_ERK_gexpslope | SPRY protein production rate catalyzed by active ERK | $h^{-1}$ | 2 | -1 |
| 47 | synthesize_ERKphosphop _SPRY_kdeg | SPRY degradation rate | $h^{-1}$ | -0.5 | -3 |
| 48 | bind_GRB2_SOS1_kf | Forward rate of GRB2 binding to SOS1 | $\mu M^{-1} \cdot h^{-1}$ | 8 | 3 |
| 49 | bind_GRB2_SOS1_kD | Dissociation constant for GRB2 binding to GRB2 | μM | 0 | -4 |
| 50 | catalyze_ERKphosphop _SOS1_pS1134_kbase | Turnover number of basal SOS1 phosphorylation | $\mu M \cdot h^{-1}$ | 1 | -2 |
| 51 | catalyze_ERKphosphop _SOS1_pS1134_kcat | Turnover number of SOS1 phosphorylation by an active ERK | $h^{-1}$ | 7 | 3 |
| 52 | ep_SOS1S1134p _GRB2_deltaG | Thermodynamic factor that modifies the binding affinity of SOS1 phosphorylated at S1134 to GRB2 | no unit | 4 | 0 |

| | | | | | |
|---|---|---|---|---|---|
| 53 | bind_SOS1_RAS_kf | Forward rate of SOS1 binding to RAS | $\mu M^{-1} \cdot h^{-1}$ | 8 | 3 |
| 54 | bind_SOS1_RAS_kD | Dissociation constant for SOS1 binding to RAS | μM | 0 | -4 |
| 55 | catalyze_SOS1_RAS_gtp_kcat | Turnover number of SOS1 loading GTP on Ras | $h^{-1}$ | 7 | 3 |
| 56 | catalyze_PP2A_MEK_u_kcatr | Ratio between the turnover rate of phosphorylation and dephosphorylation of MEK | no unit | 1 | -4 |
| 57 | catalyze_DUSP_ERK_u_kcatr | Ratio between the turnover rate of phosphorylation and dephosphorylation of ERK | no unit | 1 | -4 |
| 58 | catalyze_phosphatase _SOS1_uS1134_kcatr | Ratio between the turnover rate of phosphorylation and dephosphorylation of SOS1 at S11134 | no unit | 1 | -4 |
| 59 | catalyze_NF1_ RAS_gdp_kcatr | Ratio between the turnover rate of GTP loading and GTP hydrolysis of RAS | no unit | 1 | -4 |
| 60 | BRAF_0 | BRAF concentration | copies /cell | 3.5 | 2.5 |
| 61 | CRAF_0 | CRAF concentration | copies /cell | 4.5 | 3.5 |
| 62 | RAS_0 | RAS concentration | copies /cell | 5 | 4 |
| 63 | MEK_0 | MEK concentration | copies /cell | 5.5 | 4.5 |
| 64 | ERK_0 | ERK concentration | copies /cell | 5.5 | 4.5 |
| 65 | GRB2_0 | GRB2 concentration | copies /cell | 5 | 4 |
| 66 | SOS1_0 | SOS1 concentration | copies /cell | 4.5 | 3.5 |
| 67 | pMEK_IF_scale | Conversion factor from immunofluorescence (a.u.) to concentrations (μM) for pMEK | a.u./μM | 2 | 0 |
| 68 | pMEK_IF_offset | Offset from immunofluorescence (a.u.) to concentrations (μM) for pMEK | a.u./μM | -0.5 | -3 |
| 69 | pERK_IF_scale | Conversion factor from immunofluorescence (a.u.) to concentrations (μM) for pERK | a.u./μM | 2 | 0 |

| | | | | | |
|---|---|---|---|---|---|
| 70 | pERK_IF_offset | Offset from immunofluorescence (a.u.) to concentrations (μM) for pERK | a.u./μM | -0.5 | -3 |

**Parameters set manually**

We list the 7 parameters that control the contribution of forward and backward rates to overall rate changes coming from the energy of formations of reactants, also called rate distribution parameters ϕ (phi) (Sekar et al., 2017). A value of 1 and 0 for ϕ (phi) indicate that changes in energy of formation are reflected on the forward and backward rate, respectively. Choices between 0 and 1 that split the contribution between forward and backward rates are possible, but we decided to model binding reactions among proteins as having a ϕ=1 and binding reactions among a protein and a drug as having a ϕ=0. This reflects the idea that cooperativity in protein-protein interactions modifies their association rate, while cooperativity in protein-drug interactions modifies the dissociation rate. Each energy-based reaction has a rate distribution parameter ϕ.

| | Parameter name | Description | Unit | Value |
|---|---|---|---|---|
| 1 | bind_RASstategtp_RAF_phi | Rate distribution parameter for binding of Ras-GTP to RAF | no units | 1 |
| 2 | bind_RAF_RAF_phi | Rate distribution parameter for BRAF and CRAF homo- and hetero -dimerization | no unit | 1 |
| 3 | bind_RAFi_RAF_phi | Rate distribution parameter for binding of RAF inhibitor to BRAF and CRAF | no unit | 0 |
| 4 | bind_RAF_MEKphosphou_phi | Rate distribution parameter for binding of RAF to unphosphorylated MEK | no unit | 1 |
| 5 | bind_MEKi_MEK_phi | Rate distribution parameter for binding of MEK inhibitor to MEK | no unit | 0 |
| 6 | bind_EGF_EGFR_phi | Rate distribution parameter for binding of EGF to EGFR | no unit | 1 |
| 7 | bind_EGFR_EGFR_phi | Rate distribution parameter for EGFR dimerization | no unit | 1 |

**Input parameters**

We list the 4 input parameters that are used to set up initial conditions for pre-treatment and subsequent perturbations in the MAPK cascade.

| | Parameter name | Description | Unit |
|---|---|---|---|
| 1 | EGF_0 | Concentration of externally supplemented EGF | ng/mL |
| 2 | RAFi_0 | Concentration of the RAF inhibitor | μM |

| | | | |
|---|---|---|---|
| 3 | MEKi_0 | Concentration of the MEK inhibitor | μM |
| 4 | EGFR_crispr | Multiplier of basal EGFR expression. It can simulate basal (=1), CRISPRa overexpression (>1, typically =10) or CRISPRi downregulation (<1, typically =0.001) of EGFR. | no unit |

**Expressions**

There are 96 expressions defined in MARM1. A key role for expressions is to define energies in terms of kinetic rates, so that the model is parameterized on the kinetic rates but energy rules can be instantiated using energy parameters. Expressions are also used to define the backward rate of a reaction as the multiplication of its forward rate and its dissociation constant. Such parameterization facilitates the model fitting procedures because it resolves some scaling dependencies among parameters. We also use expression for converting initial conditions of species from copy number per cell to concentrations, and to defined constant terms such as the cell volume and Avogadro number.

**Species**

We list the 13 species that are modeled in MARM, specifying their type (Protein, Ligand or Drug), the biological entities they represent (e.g. ERK in the model stands for both ERK1 and ERK2 proteins), which parameter sets their initial abundance (unless the abundance is set within the model by feedback or other mechanisms; in that case abundance is set by reaction rates during simulation), the mechanisms that controls their abundance (constant, regulated or an external perturbation).

| | **Specie** | **Type** | **Represents** | **Initialization** | **Abundance is** |
|---|---|---|---|---|---|
| 1 | EGFR | Protein | EGFR | model | Regulated |
| 2 | GRB2 | Protein | GRB2 | GRB2_0 | Constant |
| 3 | SOS1 | Protein | SOS1 | SOS1_0 | Constant |
| 4 | SPRY2 | Protein | SPRY2+SPRY4 | model | Regulated |
| 5 | RAS | Protein | HRAS+NRAS+KRAS | RAS_0 | Constant |
| 6 | $BRAF^{V600E}$ | Protein | $BRAF^{V600E}$ | BRAF_0 | Constant |
| 7 | CRAF | Protein | CRAF | model | Constant |
| 8 | MEK | Protein | MEK1+MEK2 | MEK_0 | Constant |
| 9 | ERK | Protein | ERK1+ERK2 | ERK_0 | Constant |
| 10 | DUSP | Protein | DUSP4+DUSP6 | model | Regulated |
| 11 | EGF | Ligand | EGF | EGF_0 | Perturbation |
| 12 | RAFi | Drug | Vemurafenib | RAFi_0 | Perturbation |
| 13 | MEKi | Drug | Cobimetinib | MEKi_0 | Perturbation |

**Species modifications**

Some species are modified through catalysis and these modifications effect their activity. Here we list these modifications and how they affect the specie's activity.

| | **Specie's name** | **Modification** | **Description** |
|---|---|---|---|

| | | | |
|---|---|---|---|
| 1 | RAS | State:‘gdp’ or ‘gtp’ | Defines if Ras is GDP or GTP bound. Ras that is GTP bound binds more effectively to BRAF or CRAF, increasing their homo- and hetero-dimerization |
| 2 | MEK | Phospho: ‘u’ or ‘p’ | Defines if MEK is phosphorylated (‘p’) at activating sites S217/S221. Phosphorylated MEK is catalytic active and can phosphorylate ERK. |
| 3 | ERK | Phospho: ‘u’ or ‘p’ | Defines if ERK is phosphorylated (‘p’) at activating sites T202/Y204. Phosphorylated ERK is active and can phosphorylate and induce protein synthesis of various downstream and upstream effectors. |
| 4 | SOS1 | S1134: ‘u’ or ‘p’ | Defines if SOS1 is phosphorylated (‘p’) at the inhibitory site S1134. When SOS1 is phosphorylated it has a reduced affinity for GRB2. |

**Species binding sites**

We list the binding sites defined for each species. Binding sites are used to establish binding among species and create specie complexes.

| | Specie | | Binding site | Description |
|---|---|---|---|---|
| 1 | EGF | 1 | rtk | Binding site to EGFR |
| 2 | EGFR | 1 | rtk | Binding site to EGF |
| | | 2 | rtkf | Binding site to EGFR |
| | | 3 | SH2 | Binding site to GRB2 |
| 3 | GRB2 | 1 | SH2 | Binding site to EGFR |
| | | 2 | SH3 | Binding site to SOS1 and SPRY2 |
| 4 | SPRY | 1 | SH3 | Binding site to GRB2 |
| 5 | SOS1 | 1 | SH3 | Binding site to GRB2 |
| | | 2 | ras | Binding site to RAS |
| 6 | RAS | 1 | sos1 | Binding site to SOS1 |
| | | 2 | raf | Binding site to BRAF and CRAF |
| 7 | BRAF | 1 | RBD | Binding site to RAS |
| | | 2 | raf | Binding site to BRAF and CRAF |
| | | 3 | rafi | Binding site to RAF inhibitor |
| | | 4 | mek | Binding site to MEK |
| 8 | CRAF | 1 | RBD | Binding site to RAS |
| | | 2 | raf | Binding site to BRAF and CRAF |
| | | 3 | rafi | Binding site to RAF inhibitor |
| | | 4 | mek | Binding site to MEK |
| 9 | MEK | 1 | Dsite | Binding site to ERK |
| | | 2 | meki | Binding site to MEK inhibitor |
| | | 3 | Raf | Binding site to BRAF and CRAF |
| 10 | ERK | 1 | CD | Binding site to MEK and DUSP |
| 11 | DUSP | 1 | erk | Binding site to ERK |

| | | | | |
|---|---|---|---|---|
| 12 | RAFi | 1 | raf | Binding site to BRAF and CRAF |
| 13 | MEKi | 1 | mek | Binding site to MEK |

**Observables**

We list the 10 observables we used to extract biological relevant information from model simulations shown in main and supplementary text figures. Observables allow to quantify the activities of protein complexes controlled by multiple mechanisms. Observable typically sum up the contribution of individual protein complexes to estimate an overall biological activity. For example, the observable "drug free $RAF_2$" is the summation of all species containing a BRAF and CRAF homo- and hetero- dimers that are not bound by RAF inhibitors, thus quantifying the total catalytic activity of RAF dimers that are spread over many of the 1007 complex species. We provide the names by which observables are encoded in the model and the mnemonic names used in **Figure 5** and **Figure 6** of the main text.

| N | Observable's name in figures | Observable's name in model | Description |
|---|---|---|---|
| 1 | EGF-activated SOS1 | SOS1_active | SOS1 bound to a GRB2 that is bound to a activated EGFR |
| 2 | Ras-GTP | RAS_gtp | Ras in the GTP-bound state (state='gtp') |
| 3 | drug free $RAF_2$ | active_RAF_dimers | BRAF/CRAF in a homo- or- hetero dimer not bound by RAFi |
| 4 | drug free $BRAF^{V600E}$ | active_RAF_monomers | $BRAF^{V600E}$ in monomeric form not bound by RAFi |
| 5 | pMEK | pMEK | MEK in the phosphorylated state (phospho ='p') |
| 6 | drug free pMEK | active_MEK | MEK in the phosphorylated state (phospho='p') and not bound by MEKi |
| 7 | pERK | pERK | ERK in the phosphorylated state (phospho='p') |
| 8 | pSOS1 | pS1134SOS1 | SOS1 in the phosphorylated state at S1134 (S1134='p') |
| 9 | SPRY | tSPRY | total SPRY |
| 10 | DUSP | tDUSP | total DUSP |

**Energy patterns**

We list the 25 energy patterns used to defined the energy of formation of reactants.

| | Energy Pattern Name | Parameter | Description |
|---|---|---|---|
| 1 | ep_BRAF_BRAF_mod _RAS_double | log(ep_RAF_RAF_mod_ RASstategtp_double_deltaG) | Modifier of BRAF and CRAF homo/hetero dimerization by Ras-GTP binding |
| 2 | ep_BRAF_CRAF_mod _RAS_double | | |
| 3 | ep_CRAF_CRAF_mod _RAS_double | | |

| | | | |
|---|---|---|---|
| 4 | ep_bind_RASstategtp_BRAF | log(bind_RASstategtp_RAF_kD) | Basal binding of BRAF and CRAF to Ras-GTP |
| 5 | ep_bind_RASstategtp_CRAF | | |
| 6 | ep_bind_BRAF_BRAF | log(bind_RAF_RAF_kD) | Basal binding of BRAF and CRAF homo/hetero dimerization |
| 7 | ep_bind_BRAF_CRAF | | |
| 8 | ep_bind_CRAF_CRAF | | |
| 9 | ep_BRAF_BRAF_mod _RAFi_single | log(ep_RAF_RAF_mod _RAFi_double_deltaG) + log(ep_RAF_RAF_mod _RAFi_single_deltaG) | Modifier of BRAF and CRAF homo/hetero dimerization by RAFi binding to only one of the RAF protomers |
| 10 | ep_BRAF_CRAF_mod _RAFi_single | | |
| 11 | ep_CRAF_BRAF_mod _RAFi_single | | |
| 12 | ep_CRAF_CRAF_mod _RAFi_single | | |
| 13 | ep_BRAF_BRAF_mod _RAFi_double | log(ep_RAF_RAF_mod _RAFi_single_deltaG) | Modifier of BRAF and CRAF homo/hetero dimerization by RAFi binding to both RAF protomers |
| 14 | p_BRAF_CRAF_mod _RAFi_double | | |
| 15 | ep_CRAF_CRAF_mod _RAFi_double | | |
| 16 | ep_bind_BRAFi_RAF | log(bind_RAFi_RAF_kD) | Basal binding of BRAF and CRAF to the RAF inhibitor |
| 17 | ep_bind_CRAFi_RAF | | |
| 18 | ep_bind_BRAF_MEKphosphou | log(bind_RAF_ MEKphosphou_kD) | Basal binding of BRAF or CRAF to unphosphorylated MEK |
| 19 | ep_bind_CRAF_MEKphosphou | | |
| 20 | ep_bind_MEKi_MEK | log(bind_MEKi_MEK_kD) | Basal binding of unphosphorylated MEK to the MEK inhibitor |
| 21 | ep_MEKphosphop_MEKi_single | log(ep_MEKphosphop _MEKi_deltaG) | Modifier of phosphorylated MEK binding to the MEK inhibitor |
| 22 | ep_EGFR_EGFR_mod_EGF_single | log(ep_EGFR_EGFR_mod _EGF_single_deltaG) | Modifier of EGFR homo- |

| | | | |
|---|---|---|---|
| | | | dimerization by EGF binding to only one protomer |
| 23 | ep_EGFR_EGFR_mod_EGF_double | log(ep_EGFR_EGFR_mod_EGF_double_deltaG) + log(ep_EGFR_EGFR_mod_EGF_single_deltaG) | Modifier of EGFR homo-dimerization by EGF binding to both protomers |
| 24 | ep_bind_EGF_EGFR | log(bind_EGF_EGFR_kD) | Basal binding of EGF and EGFR |
| 25 | ep_bind_EGFR_EGFR | log(bind_EGFR_EGFR_kD) | Basal dimerization of EGFR |

**Energy-based reaction rules**

We list the 12 energy-based rules used to model cooperativity and define reaction networks that satisfy detailed balanced by construction. For each energy-based rule, the parameters listed are the rate distribution parameter ϕ (phi), and the forward rate and dissociation constant that are used to defined the energy of activation Ea_0. Thus, the model is parameterized on kinetic rates by the internal representation is based on energy.

| N | Reaction Name | Parameters | Description |
|---|---|---|---|
| 1 | RASgtp_and_BRAF_bind_and_dissociate | bind_RASstategtp_RAF_phi ; log(bind_RASstategtp_RAF_kD) ; log(bind_RASstategtp_RAF_kf) | Ras-GTP binding to BRAF or CRAF |
| 2 | RASgtp_and_CRAF_bind_and_dissociate | | |
| 3 | BRAF_and_BRAF_bind_and_dissociate | bind_RAF_RAF_phi ; log(bind_RAF_RAF_kD) ; log(bind_RAF_RAF_kf) | BRAF and CRAF homo/hetero dimerization |
| 4 | BRAF_and_CRAF_bind_and_dissociate | | |
| 5 | CRAF_and_CRAF_bind_and_dissociate | | |
| 6 | RAFi_and_BRAF_bind_and_dissociate | bind_RAFi_RAF_phi ; log(bind_RAFi_RAF_kD) ; log(bind_RAFi_RAF_kf) | RAFi binding to BRAF or CRAF |
| 7 | RAFi_and_CRAF_bind_and_dissociate | | |
| 8 | BRAF_and_uMEK_bind_and_dissociate | bind_RAF_MEKphosphou_phi ; log(bind_RAF_MEKphosphou_kD) ; log(bind_RAF_MEKphosphou_kf) | BRAF or CRAF binding to unphosphorylated MEK |
| 9 | CRAF_and_uMEK_bind_and_dissociate | | |
| 10 | MEKi_and_MEK_bind_and_dissociate | bind_RAF_MEKphosphou_phi ; log(bind_RAF_MEKphosphou_kD) ; log(bind_RAF_MEKphosphou_kf) | MEKi binding to unphosphorylated MEK |

| | | | |
|---|---|---|---|
| 11 | EGF_and_EGFR _bind_and_dissociate | bind_EGF_EGFR_phi ; log(bind_EGF_EGFR_kD) ; log(bind_EGF_EGFR_kf) | EGF binding to EGFR |
| 12 | EGFR_and_EGFR _bind_and_dissociate | bind_EGFR_EGFR_phi ; log(bind_EGFR_EGFR_kD) ; log(bind_EGFR_EGFR_kf) | EGFR dimerization |

**Reaction rules**

We list the 40 non energy-based 'classic' rules used to define reaction networks. Note that backward reaction rates are defined as the product between the forward rate and the dissociation constant, which are the parameters inferred from data.

| N | Reaction Name | Parameter | Description |
|---|---|---|---|
| 1 | GRB2_degrades_activeRTK_dimers | catalyze_GRB2 _RTKSH2ANY_deg_kcat | Degradation of active EGFR dimer bound by GRB2 |
| 2 | GRB2_degrades_doubleactiveRTK_dimers | catalyze_GRB2 _RTKSH2ANY_deg_kcat | Degradation of both active EGFR dimers bound by GRB2 |
| 3 | GRB2_degrades_activeRTK_monomers | catalyze_GRB2 _RTKSH2ANY_deg_kcat | Degradation of active EGFR monomer bound by GRB2 |
| 4 | BRAF_BRAF_phosphorylates_MEK | catalyze_RAFrafiNone _MEK_p_kcat | Phosphorylation of MEK by either BRAF and CRAF homo- and hetero-dimer or by monomeric $BRAF^{V600E}$ |
| 5 | BRAF_CRAF_phosphorylates_MEK | | |
| 6 | CRAF_BRAF_phosphorylates_MEK | | |
| 7 | CRAF_CRAF_phosphorylates_MEK | | |
| 8 | BRAFV600E_phosphorylates_MEK_1 | | |
| 9 | BRAFV600E_phosphorylates_MEK_2 | | |
| 10 | BRAFV600E_phosphorylates_MEK_3 | | |
| 11 | BRAFV600E_phosphorylates_MEK_4 | | |
| 12 | MEK_is_dephosphorylated | catalyze_PP2A _MEK_u_kcat | MEK is dephosphorylated |
| 13 | MEK_binds_uERK | bind_MEK _ERKphosphou_kf | MEK binds unphosphorylated ERK |
| 14 | MEK_dissociates_from_ERK | bind_MEK_ERKphosphou_kf * bind_MEK_ERKphosphou_kD | MEK dissociates unphosphorylated ERK |
| 15 | pMEK_phosphorylates_ERK | catalyze_MEKmekiNone _phosphop_ERK_p_kcat | MEK phosphorylates |

| | | | |
|---|---|---|---|
| | | | and dissociates from ERK |
| 16 | DUSP_binds_pERK | bind_DUSP_ERKphosphop_kf | DUSP binding to phosphorylated ERK |
| 17 | DUSP_dissociates_from_ERK | bind_DUSP_ERKphosphop_kf* bind_DUSP_ERKphosphop_kD | DUSP unbinding from phosphorylated ERK |
| 18 | DUSP_dephosphorylates_ERK | catalyze_DUSP_ERK_u_kcat | DUSP dephosphorylates and dissociates from ERK |
| 19 | basal_synthesis_DUSP | synthesize_ERKphosphop _DUSP_ksyn | DUSP basal protein synthesis |
| 20 | basal_degradation_DUSP | synthesize_ERKphosphop _DUSP_kdeg | DUSP protein degradation |
| 21 | ERK_synthesizes_DUSP | synthesize_ERKphosphop _DUSP_kmodslope | DUSP protein synthesis positively regulated by activated ERK |
| 22 | basal_synthesis_EGFR | synthesize_ERKphosphop _EGFR_ksyn | EGFR basal protein synthesis |
| 23 | basal_degradation_EGFR | synthesize_ERKphosphop _EGFR_kdeg | EGFR protein degradation |
| 24 | ERK_synthesizes_EGFR | synthesize_ERKphosphop _EGFR_kmodslope | EGFR protein synthesis positively regulated by activated ERK |
| 25 | transactivated_RTK _dimers_bind_GRB2 | bind_RTK_GRB2_kf | GRB2 binds to a EGFR protomer in a dimer that is transactivated by EGF binding to the other protomer |
| 26 | GRB2_dissociates _from_RTK | bind_RTK_GRB2_kf * bind_RTK_GRB2_kD | GRB2 unbinding from EGFR |
| 27 | SPRY_binds_GRB2 | bind_SPRY_GRB2_kf | SPRY2 binding to GRB2 |
| 28 | SPRY_dissociates_from_GRB2 | bind_SPRY_GRB2_kf * bind_SPRY_GRB2_kD | SPRY2 unbinding from GRB2 |
| 29 | basal_synthesis_SPRY | synthesize_ERKphosphop _SPRY_ksyn | SPRY basal protein synthesis |

| | | | |
|---|---|---|---|
| 30 | basal_degradation_SPRY | synthesize_ERKphosphop _SPRY_kdeg | SPRY protein degradation |
| 31 | ERK_synthesizes_SPRY | synthesize_ERKphosphop _SPRY_kmodslope | SPRY protein synthesis positively regulated by activated ERK |
| 32 | SOS1_is_dephosphorylated | catalyze_phosphatase _SOS1_uS1134_kcat | SOS1 basal dephosphorylation |
| 33 | GRB2_binds_SOS1 | bind_GRB2_SOS1_kf | GRB2 binding to SOS1 |
| 34 | GRB2_dissociates_from_SOS1 | bind_GRB2_SOS1_kf * bind_GRB2_SOS1_kD | GRB2 unbinding from SOS1 |
| 35 | SOS1_is_phosphorylated | catalyze_ERKphosphop _SOS1_pS1134_kbase | SOS1 baseline phosphorylation |
| 36 | pERK_phosphorylates_SOS1 | catalyze_ERKphosphop _SOS1_pS1134_kcat | SOS1 phosphorylation by activated ERK |
| 37 | pSOS1_has_modulated _affinity_to_GRB2 | bind_GRB2_SOS1_kD * bind_GRB2_SOS1_kf * ep_SOS1S1134p_GRB2_deltaG | SOS1 phosphorylated at S1134 reduces its affinity to GRB2 |
| 38 | RTK_and_GRB2_bound _SOS1_binds_RASgdp | bind_SOS1_RAS_kf | GDP-bound Ras binds a GRB2 that is bound to an EGFR dimer |
| 39 | SOS1_dissociates_from_RAS | bind_SOS1_RAS_kf * bind_SOS1_RAS_kD | SOS1 unbinding from Ras in GDP- or GTP- bound form |
| 40 | SOS1_catalyzes_RAS _guanosine_exchange | catalyze_SOS1_ RAS_gtp_kcat | SOS1 catalyzes the GTP-loading on Ras (this interaction does not require binding of SOS1 to Ras) |
| 41 | RAS_hydrolisis_GTP | catalyze_NF1 _RAS_gdp_kcat | GTP-bound RAS is hydrolyzed to GDP- bound RAS |
| 42 | GTP_hydrolysis _dissociates_BRAF_from_RAS | catalyze_NF1 _RAS_gdp_kcat | Hydrolysis of GTP-bound RAS while bound to BRAF or CRAF also disassemble |
| 43 | GTP_hydrolysis _dissociates_CRAF_from_RAS | | |

| | | | |
|---|---|---|---|
| | | | it from the RAF molecule2 |

**Detailed description of mechanisms and assumptions in MARM1**

Here below, we describe the molecular mechanisms encoded in MARM1. This is the text that can be found in the Jupyter Notebook *MARM1_construction.ipynb* provided at Github (https://github.com/labsyspharm/marm1-supplement).The notebook allows to follow the description of the implemented molecular mechanisms together with a step-by-step construction of the MARM1 PySB model.

In this notebook we will provide a step-by-step construction of the MARM model. The model is constructed using the rule-based modeling tool PySB . To start, we import all required pysb classes and instantiate the model:

```
from pysb import Model, Monomer, Parameter, Expression,  Rule, Observable, In
itial, Annotation, EnergyPattern, ANY
from pysb.bng import generate_equations
from pysb.export import export
from sympy import exp, log
```

```
Model();
```

In the following, we split the model into individual submodules of strongly interacting components, which we will describe individually. The model describes protein and drug species as intracellular concentrations in uM at a volume of `1pL = 1e-12L`. For ligands and drugs we assume that the extracellular compartment is much bigger than the intracellular compartement and that equilibration is fast such that concentrations can be assumed to be constanct over time (implemented by specifying `fixed=True` in the initialization). For EGF, the parameter that specifies the abundance, `EGF_0` is assumed to have units [ng/ml] and is accordingly transformed into [uM] using the appropriate molecular weight `m_Da_EGF`. For the drugs Vemurafenib (`RAFi`) and Cobimetinib (`MEKi`), the respective parameters, `RAFi_0` and `MEKi_0`, are assumed to be in uM. The unit of time is hours.

```
Expression('N_Avogadro', 6.02214085700000e+23)
Expression('volume', 1.00000000000000e-12)
Expression('m_Da_EGF', 6200.00000000000)

Monomer('RAFi', ['raf'])
Annotation(RAFi, 'http://identifiers.org/chebi/63637', 'is')
Parameter('RAFi_0', 0.0)
Expression('initRAFi', RAFi_0)
Initial(RAFi(raf=None), initRAFi, fixed=True)

Monomer('MEKi', ['mek'])
```

```
Annotation(MEKi, 'http://identifiers.org/chebi/90851', 'is')
Parameter('MEKi_0', 0.0)
Expression('initMEKi', MEKi_0)
Initial(MEKi(mek=None), initMEKi, fixed=True)

Monomer('EGF', ['rtk'])
Annotation(EGF, 'http://identifiers.org/uniprot/P01133', 'is')
Parameter('EGF_0', 0.0)
Expression('initEGF', 6.02214085854916e+23*EGF_0*(m_Da_EGF*N_Avogadro)**(-1))
Initial(EGF(rtk=None), initEGF, fixed=True);
```

### *EGFR Signaling*

The EGFR submodule describes the interaction between Egfr, Egf, Grb2, Sos1 and Ras (which describes the action K-Ras, N-Ras and H-Ras). Here, we instantiate the respective monomers and provide uniprot IDs as monomer annotations.

```
Monomer('EGFR', ['SH2', 'rtk', 'rtkf'])
Annotation(EGFR, 'http://identifiers.org/uniprot/P00533', 'is')

Monomer('GRB2', ['SH2', 'SH3'])
Annotation(GRB2, 'http://identifiers.org/uniprot/P62993', 'is')

Monomer('SOS1', ['S1134', 'SH3', 'ras'], {'S1134': ['p', 'u']})
Annotation(SOS1, 'http://identifiers.org/uniprot/Q07889', 'is')

Monomer('RAS', ['raf', 'sos1', 'state'], {'state': ['gdp', 'gtp']})
Annotation(RAS, 'http://identifiers.org/uniprot/P01111', 'is')
Annotation(RAS, 'http://identifiers.org/uniprot/P01116', 'is')
Annotation(RAS, 'http://identifiers.org/uniprot/P01112', 'is');
```

#### EGF activation

To describe activation of Egfr through Egf, we implement base binding rates between Egf and Egfr. We implement these rules as energy rules, which allow us to later use the energy-based formalism in BioNetGen to describe allosteric interactions between the two molecules. Energy rules are parameterised using three distinct parameters: the `kD` parameter which describes the affinity between the two molecules, the `kf` parameter which describes the timescale of the interaction and the `phi` parameter which takes values between 0 and 1 and balance whether allosteric interactions modulate the affinity by changing the association rate (`kf`, only association if `phi=0`) or the dissociation rate (`kr = kf*kD`, only dissociation if `phi=1`).

We implement the ligand-receptor binding using the `rtk` binding site on the ligand and the `rtkf` binding site on the receptor. We implement the receptor-receptor binding using the `rtk` binding site on the receptor. This formalism simplifies future extensions of the model to additional ligands or receptors by ensuring that each receptor can only bind one ligand and one receptor.

```
# binding Egf-Egfr
Parameter('bind_EGF_EGFR_kf', 10.0)
Parameter('bind_EGF_EGFR_kD', 0.01)
Parameter('bind_EGF_EGFR_phi', 1.0)
Expression('bind_EGF_EGFR_kr', bind_EGF_EGFR_kf*bind_EGF_EGFR_kD)
Expression('Gf_bind_EGF_EGFR', log(bind_EGF_EGFR_kD))
Expression('Ea0_bind_EGF_EGFR', -(bind_EGF_EGFR_phi*log(bind_EGF_EGFR_kD) + l
og(bind_EGF_EGFR_kf)))
Rule('EGF_and_EGFR_bind_and_dissociate',
     EGF(rtk=None) + EGFR(rtkf=None) | EGF(rtk=1) % EGFR(rtkf=1),
     bind_EGF_EGFR_phi, Ea0_bind_EGF_EGFR, energy=True)
EnergyPattern('ep_bind_EGF_EGFR', EGF(rtk=1) % EGFR(rtkf=1), Gf_bind_EGF_EGFR
)

# binding Egfr-Egfr
Parameter('bind_EGFR_EGFR_kf', 10.0)
Parameter('bind_EGFR_EGFR_kD', 100.0)
Parameter('bind_EGFR_EGFR_phi', 1.0)
Expression('bind_EGFR_EGFR_kr', bind_EGFR_EGFR_kf*bind_EGFR_EGFR_kD)
Expression('Gf_bind_EGFR_EGFR', log(bind_EGFR_EGFR_kD))
Expression('Ea0_bind_EGFR_EGFR', -(bind_EGFR_EGFR_phi*log(bind_EGFR_EGFR_kD)
+ log(bind_EGFR_EGFR_kf)))
Rule('EGFR_and_EGFR_bind_and_dissociate',
     EGFR(rtk=None) + EGFR(rtk=None) | EGFR(rtk=1) % EGFR(rtk=1),
     bind_EGFR_EGFR_phi, Ea0_bind_EGFR_EGFR, energy=True)
EnergyPattern('ep_bind_EGFR_EGFR', EGFR(rtk=1) % EGFR(rtk=1), Gf_bind_EGFR_EG
FR);
```

To implement allosteric interaction between Egf ligands and Egfr , we add two energy patterns that allow modulation of the affinities of Egfr-Egf and Egfr-Egfr interactions through binding of ligands.

```
# Egf mediated affinity modulation
Parameter('ep_EGFR_EGFR_mod_EGF_single_deltaG', 0.001)
Parameter('ep_EGFR_EGFR_mod_EGF_double_deltaG', 0.001)
Expression('ep_EGFR_EGFR_mod_EGF_single_Gf', log(ep_EGFR_EGFR_mod_EGF_single_
deltaG))
Expression('ep_EGFR_EGFR_mod_EGF_double_Gf',
           log(ep_EGFR_EGFR_mod_EGF_double_deltaG) + log(ep_EGFR_EGFR_mod_EGF
_single_deltaG))
EnergyPattern('ep_EGFR_EGFR_mod_EGF_single',
              EGFR(rtk=1, rtkf=None) % EGFR(rtk=1, rtkf=2) % EGF(rtk=2),
              ep_EGFR_EGFR_mod_EGF_single_Gf)
EnergyPattern('ep_EGFR_EGFR_mod_EGF_double',
```

```
              EGF(rtk=2) % EGFR(rtk=1, rtkf=2) % EGFR(rtk=1, rtkf=3) % EGF(rt
k=3),
              ep_EGFR_EGFR_mod_EGF_double_Gf);

Annotation(EGF_and_EGFR_bind_and_dissociate, 'http://identifiers.org/pubmed/1
6946702', 'isDescribedBy')
Annotation(EGFR_and_EGFR_bind_and_dissociate, 'http://identifiers.org/pubmed/
16946702', 'isDescribedBy');
```

#### GRB2 recruitement

Recruitement of Grb2 to receptor requires transactivation of multiple Egfr phosphorylation sites that are recognized by the Grb2 Src2 homology 2 (SH2) domain . To simplify these interactions we require Egfr to be bound to an activating ligand-Egfr complex to recruit Grb2 to the plasma membrane. To simplify notation the binding site on the Egfr monomer is also called `SH2`, although Egfr does not harbor any SH2 domain.

The dissociation is implemented as unconditional rule to avoid the creation of irreversible states (e.g, dissociation of receptor dimers could protect from dissociation of Grb2). Accordingly, this rule is not implemented as energy rule, as energy rules always require the same conditions for association and dissociation rules. Nevertheless, we parameterize the rule using `kf` and `kD` parameters, although, no `phi` parameter is required here as the rule kinetics are never modulated through energy patterns.

```
# EGFR-Grb2 binding
Parameter('bind_RTK_GRB2_kf', 10.0)
Parameter('bind_RTK_GRB2_kD', 0.01)
Expression('bind_RTK_GRB2_kr', bind_RTK_GRB2_kf*bind_RTK_GRB2_kD)
Rule('transactivated_RTK_dimers_bind_GRB2',
     EGFR(rtk=1, rtkf=ANY) % EGFR(SH2=None, rtk=1) + GRB2(SH2=None)
     >>
     EGFR(rtk=1, rtkf=ANY) % EGFR(SH2=2, rtk=1) % GRB2(SH2=2),
     bind_RTK_GRB2_kf)
Rule('GRB2_dissociates_from_RTK',
     EGFR(SH2=1) % GRB2(SH2=1) >> EGFR(SH2=None) + GRB2(SH2=None),
     bind_RTK_GRB2_kr)

Annotation(transactivated_RTK_dimers_bind_GRB2, 'http://identifiers.org/pubme
d/16777603', 'isDescribedBy');
```

The recruitement of Grb2 to Egfr is not oly important for signal transduction, but also regulates the balance between slower, clathrin-independent and faster, clathrin-dependent (Grb2 dependent) receptor endocytosis .

We describe the baseline Egfr expression (determined by clathrin-independent endocytosis) using basal Egfr expression and synthesis rules. Similar to binding rules, synthesis rules are parameterized equilibrium abundance parameter `[]_eq`, which regulates steady-state behaviour, and degradation rate `[]_kdeg`, which regulates the timescale of the rule. The equilibrium abundance can be modulated by the `[]_crispr` factor by up or down-regulating the synthesis rate `[]_ksyn`. To simplify specification of estimation boundaries for the `[...]_eq` parameter, we assume that the parameter is specified in [molecules/cell] and accordingly rescale the respective synthesis rate to [uM/h].

We implement clathrin-dependent endocytosis as three seperate rules for Egfr dimers with one Grb2 bound, with two Grb2 bound and Grb2 bound Egfr monomers.
The `delete_molecules=True` parameter encodes that only Egfr molecules are degradede and all other molecules are remain as possibly fragmented comlexes. The formulation as three seperate rules ensures that Egfr endocytosis rate constant with respect to Grb2 stochiometry in the complex and that dissociation of Egfr homodimers does not protect from endocytosis.

```
# basal synthesis + degradation
Parameter('EGFR_eq', 10000.0)
Parameter('synthesize_ERKphosphop_EGFR_kdeg', 10.0)
Parameter('EGFR_crispr', 1.0)
Expression('synthesize_ERKphosphop_EGFR_ksyn',
           1000000.0*EGFR_eq*(N_Avogadro*volume)**(-1)*synthesize_ERKphosphop
_EGFR_kdeg*EGFR_crispr)
Rule('basal_synthesis_EGFR',
     None >> EGFR(SH2=None, rtk=None, rtkf=None),
     synthesize_ERKphosphop_EGFR_ksyn)
Rule('basal_degradation_EGFR',
     EGFR() >> None,
     synthesize_ERKphosphop_EGFR_kdeg,
     delete_molecules=True)

# Grb2 mediated degradation
Parameter('catalyze_GRB2_RTKSH2ANY_deg_kcat', 10.0)
Rule('GRB2_degrades_activeRTK_dimers',
     EGFR(SH2=ANY, rtk=1) % EGFR(SH2=None, rtk=1) >> None,
     catalyze_GRB2_RTKSH2ANY_deg_kcat,
     delete_molecules=True)
Rule('GRB2_degrades_doubleactiveRTK_dimers',
     EGFR(SH2=ANY, rtk=1) % EGFR(SH2=ANY, rtk=1) >> None,
     catalyze_GRB2_RTKSH2ANY_deg_kcat,
     delete_molecules=True)
Rule('GRB2_degrades_activeRTK_monomers',
     EGFR(SH2=ANY, rtk=None) >> None,
     catalyze_GRB2_RTKSH2ANY_deg_kcat,
     delete_molecules=True);
```

##### SOS1 activation

Grb2 recruitement by Egfr also induces association of Sos1 to the plasma-membrane, where it can activate Ras by inducing nucleotide exchange . In the model we implement that Grb2 and Sos1 preform complexes in the cytoplasm and that Sos1 is activated through association to the plasma-membrane, which is mediated recruitement of Grb2 by Egfr. Given that the training data does not include any data or perturbations at the Ras level, we implement a simple catalysis rule for activation of Ras, where SOS1 only binds GDP bound Ras and catalysis is implemented without homonucleotide exchange, allosteric feedback mechanisms or interactions with cdc25 . Similarly, we implement inactivation of Ras through GTPase activation proteins, such as NF1, as a simple linear reaction with constant rate.

As previous binding reactions, the rules are paramterized using `kf` and `kD` values. As for the Grb2-Egfr interaction, the binding domains on Sos1 and Grb2 are implemented as `SH3` sites on both molecules, simplifying the much more intricate binding dynamics . The translocation of Grb2 and Sos1 to the plasma membrane is not explicitely implemented in the model but implicitely encoded by requiring Grb2 and Egfr association for binding to Ras. To reduce correlations between parameter estimates, we implement the inactivation rate `[...]_gdp_kcat` of Ras as product of the activation rate `[...]_gtp_kcat` and a scaling factor `[...]_gdp_kcatr`. The requirement of `raf=None` in the inactivation rule will be explained in the Section on Raf activation.

```
# binding Grb2-Sos1
Parameter('bind_GRB2_SOS1_kf', 10.0)
Parameter('bind_GRB2_SOS1_kD', 0.01)
Expression('bind_GRB2_SOS1_kr', bind_GRB2_SOS1_kf*bind_GRB2_SOS1_kD)
Rule('GRB2_binds_SOS1',
     GRB2(SH3=None) + SOS1(SH3=None) >> GRB2(SH3=1) % SOS1(SH3=1),
     bind_GRB2_SOS1_kf)
Rule('GRB2_dissociates_from_SOS1',
     GRB2(SH3=1) % SOS1(SH3=1) >> GRB2(SH3=None) + SOS1(SH3=None),
     bind_GRB2_SOS1_kr)

# binding Sos1-Ras
Parameter('bind_SOS1_RAS_kf', 10.0)
Parameter('bind_SOS1_RAS_kD', 0.01)
Expression('bind_SOS1_RAS_kr', bind_SOS1_RAS_kf*bind_SOS1_RAS_kD)
Rule('RTK_and_GRB2_bound_SOS1_binds_RASgdp',
     GRB2(SH2=ANY, SH3=1) % SOS1(SH3=1, ras=None) + RAS(sos1=None, state='gdp
')
     >>
     GRB2(SH2=ANY, SH3=1) % SOS1(SH3=1, ras=2) % RAS(sos1=2, state='gdp'),
     bind_SOS1_RAS_kf)
Rule('SOS1_dissociates_from_RAS',
     SOS1(ras=1) % RAS(sos1=1) >> SOS1(ras=None) + RAS(sos1=None),
     bind_SOS1_RAS_kr)

# activation + inactivation Ras
Parameter('catalyze_SOS1_RAS_gtp_kcat', 0.01)
Parameter('catalyze_NF1_RAS_gdp_kcatr', 100.0)
Expression('catalyze_NF1_RAS_gdp_kcat', catalyze_SOS1_RAS_gtp_kcat*catalyze_N
F1_RAS_gdp_kcatr)
```

```
Rule('SOS1_catalyzes_RAS_guanosine_exchange',
     SOS1(ras=1) % RAS(sos1=1, state='gdp') >> SOS1(ras=None) + RAS(sos1=None
, state='gtp'),
     catalyze_SOS1_RAS_gtp_kcat)
Rule('RAS_hydrolisis_GTP',
     RAS(raf=None, state='gtp') >> RAS(raf=None, state='gdp'),
     catalyze_NF1_RAS_gdp_kcat);
```

### *ERK signaling*

The Erk submodule describes the interaction between the two Rafs c-Raf and B-Raf, Mek (which describes Mek1 and Mek2) and Erk (which describes Erk1 and Erk2). Again, we instantiate the respective monomers and provide uniprot IDs as monomer annotations.

```
Monomer('BRAF', ['AA600', 'RBD', 'mek', 'raf', 'rafi'], {'AA600': ['E']})
Annotation(BRAF, 'http://identifiers.org/uniprot/P15056', 'is')

Monomer('CRAF', ['RBD', 'mek', 'raf', 'rafi'])
Annotation(CRAF, 'http://identifiers.org/uniprot/P04049', 'is')

Monomer('MEK', ['Dsite', 'meki', 'phospho', 'raf'], {'phospho': ['p', 'u']})
Annotation(MEK, 'http://identifiers.org/uniprot/Q02750', 'is')
Annotation(MEK, 'http://identifiers.org/uniprot/P36507', 'is')

Monomer('ERK', ['CD', 'phospho'], {'phospho': ['p', 'u']})
Annotation(ERK, 'http://identifiers.org/uniprot/P27361', 'is')
Annotation(ERK, 'http://identifiers.org/uniprot/P28482', 'is');
```

#### Raf activation

GTP-bound Ras activates Raf by recruiting it to the membrane where it is phosphorylated and dimerizes. As understanding of the exact mechanisms of this activation process are still incomplete, we implemented a simplistic model of this process that captures essential features of the interaction. We implement a base dimerization rate between Raf monomers and only allow binding of Raf monomers to GTP-bound Ras. To implement the Ras-mediated dimerization of Raf, we implement an energy pattern for Ras-Raf tetramers modulates both binding affinities.

We implement both binding rules as energy rules, where we assume that affinities for B-Raf and C-Raf are the same for all interactions. As energy rules require symmetric forward and backward reactions, inactivation of Ras would protect Raf from unbinding from Ras. Accordingly, we add two additional inactivation reactions that dissociate Raf from Ras and required that no Raf was bound in the baseline inactivation.

```
# Raf-Raf binding
Parameter('bind_RAF_RAF_kf', 10.0)
Parameter('bind_RAF_RAF_kD', 0.01)
Parameter('bind_RAF_RAF_phi', 1.0)
```

```
Expression('bind_BRAF_BRAF_kr', bind_RAF_RAF_kf*bind_RAF_RAF_kD)
Expression('Gf_bind_BRAF_BRAF', log(bind_RAF_RAF_kD))
Expression('Ea0_bind_BRAF_BRAF', -(bind_RAF_RAF_phi*log(bind_RAF_RAF_kD) + lo
g(bind_RAF_RAF_kf)))
Expression('bind_BRAF_CRAF_kr', bind_RAF_RAF_kf*bind_RAF_RAF_kD)
Expression('Gf_bind_BRAF_CRAF', log(bind_RAF_RAF_kD))
Expression('Ea0_bind_BRAF_CRAF', -(bind_RAF_RAF_phi*log(bind_RAF_RAF_kD) + lo
g(bind_RAF_RAF_kf)))
Expression('bind_CRAF_CRAF_kr', bind_RAF_RAF_kf*bind_RAF_RAF_kD)
Expression('Gf_bind_CRAF_CRAF', log(bind_RAF_RAF_kD))
Expression('Ea0_bind_CRAF_CRAF', -(bind_RAF_RAF_phi*log(bind_RAF_RAF_kD) + lo
g(bind_RAF_RAF_kf)))
Rule('BRAF_and_BRAF_bind_and_dissociate',
     BRAF(raf=None) + BRAF(raf=None) | BRAF(raf=1) % BRAF(raf=1),
     bind_RAF_RAF_phi,
     Ea0_bind_BRAF_BRAF,
     energy=True)
Rule('BRAF_and_CRAF_bind_and_dissociate',
     BRAF(raf=None) + CRAF(raf=None) | BRAF(raf=1) % CRAF(raf=1),
     bind_RAF_RAF_phi,
     Ea0_bind_BRAF_CRAF,
     energy=True)
Rule('CRAF_and_CRAF_bind_and_dissociate',
     CRAF(raf=None) + CRAF(raf=None) | CRAF(raf=1) % CRAF(raf=1),
     bind_RAF_RAF_phi,
     Ea0_bind_CRAF_CRAF,
     energy=True)
EnergyPattern('ep_bind_BRAF_BRAF', BRAF(raf=1) % BRAF(raf=1), Gf_bind_BRAF_BR
AF)
EnergyPattern('ep_bind_BRAF_CRAF', BRAF(raf=1) % CRAF(raf=1), Gf_bind_BRAF_CR
AF)
EnergyPattern('ep_bind_CRAF_CRAF', CRAF(raf=1) % CRAF(raf=1), Gf_bind_CRAF_CR
AF)

# Ras_gtp-RAF binding
Parameter('bind_RASstategtp_RAF_kf', 10.0)
Parameter('bind_RASstategtp_RAF_kD', 0.01)
Parameter('bind_RASstategtp_RAF_phi', 1.0)
Expression('bind_RASstategtp_BRAF_kr', bind_RASstategtp_RAF_kf*bind_RASstateg
tp_RAF_kD)
Expression('Gf_bind_RASstategtp_BRAF', log(bind_RASstategtp_RAF_kD))
Expression('Ea0_bind_RASstategtp_BRAF',
           -(bind_RASstategtp_RAF_phi*log(bind_RASstategtp_RAF_kD) + log(bind
_RASstategtp_RAF_kf)))
```

```
Expression('bind_RASstategtp_CRAF_kr', bind_RASstategtp_RAF_kf*bind_RASstateg
tp_RAF_kD)
Expression('Gf_bind_RASstategtp_CRAF', log(bind_RASstategtp_RAF_kD))
Expression('Ea0_bind_RASstategtp_CRAF',
           -(bind_RASstategtp_RAF_phi*log(bind_RASstategtp_RAF_kD) + log(bind
_RASstategtp_RAF_kf)))
Rule('RASgtp_and_BRAF_bind_and_dissociate',
     RAS(raf=None, state='gtp') + BRAF(RBD=None) | RAS(raf=1, state='gtp') %
BRAF(RBD=1),
     bind_RASstategtp_RAF_phi,
     Ea0_bind_RASstategtp_BRAF,
     energy=True)
Rule('RASgtp_and_CRAF_bind_and_dissociate',
     RAS(raf=None, state='gtp') + CRAF(RBD=None) | RAS(raf=1, state='gtp') %
CRAF(RBD=1),
     bind_RASstategtp_RAF_phi,
     Ea0_bind_RASstategtp_CRAF,
     energy=True)
EnergyPattern('ep_bind_RASstategtp_BRAF', RAS(raf=1, state='gtp') % BRAF(RBD=
1), Gf_bind_RASstategtp_BRAF)
EnergyPattern('ep_bind_RASstategtp_CRAF', RAS(raf=1, state='gtp') % CRAF(RBD=
1), Gf_bind_RASstategtp_CRAF)

# Ras-Raf inactivation
Rule('GTP_hydrolysis_dissociates_BRAF_from_RAS',
     RAS(raf=1, state='gtp') % BRAF(RBD=1) >> RAS(raf=None, state='gdp') + BR
AF(RBD=None),
     catalyze_NF1_RAS_gdp_kcat)
Rule('GTP_hydrolysis_dissociates_CRAF_from_RAS',
     RAS(raf=1, state='gtp') % CRAF(RBD=1) >> RAS(raf=None, state='gdp') + CR
AF(RBD=None),
     catalyze_NF1_RAS_gdp_kcat)

# Ras induces Raf dimerization
Parameter('ep_RAF_RAF_mod_RASstategtp_double_deltaG', 1000.0)
Expression('ep_RAF_RAF_mod_RASstategtp_double_Gf', log(ep_RAF_RAF_mod_RASstat
egtp_double_deltaG))
EnergyPattern('ep_BRAF_BRAF_mod_RAS_double',
              RAS(raf=2, state='gtp') % BRAF(RBD=2, raf=1) % BRAF(RBD=3, raf=
1) % RAS(raf=3, state='gtp'),
              ep_RAF_RAF_mod_RASstategtp_double_Gf)
EnergyPattern('ep_BRAF_CRAF_mod_RAS_double',
              RAS(raf=2, state='gtp') % BRAF(RBD=2, raf=1) % CRAF(RBD=3, raf=
1) % RAS(raf=3, state='gtp'),
```

```
              ep_RAF_RAF_mod_RASstategtp_double_Gf)
EnergyPattern('ep_CRAF_CRAF_mod_RAS_double',
              RAS(raf=2, state='gtp') % CRAF(RBD=2, raf=1) % CRAF(RBD=3, raf=
1) % RAS(raf=3, state='gtp'),
              ep_RAF_RAF_mod_RASstategtp_double_Gf);
```

#### Raf inhibition

Although Vemurafenib is an ATP-competitive inihibitor, it is known to exert allosteric effects. Here, we adapt a previously published model for these allosteric interactions between C-Raf, B-Raf and Vemurafenib . The model implements binding between Vemurafenib (`RAFi`) and C-Raf and B-Raf (`RAF`) as energy rule as well as two sets of energy patterns that modulate `RAF-RAF` and `RAF-RAFi` affinities in single and double drug bound RAF-RAF homo and heterodimers.

In this implementation, C-Raf and B-Raf both correspond to kinase monomers `R` of the Kholodenko Core Model (KCM). `K1` of the KCM is equivalent to the parameter `bind_RAF_RAF_kD`, which was introduced in the previous Section. `K2` in the KCM is equivalent to `bind_RAFi_RAF_kD`, `f` to `ep_RAF_RAF_mod_RAFi_single_deltaG` and `g` to `ep_RAF_RAF_mod_RAFi_double_deltaG`.

```
# Binding RAFi-Raf
Parameter('bind_RAFi_RAF_kf', 10.0)
Parameter('bind_RAFi_RAF_kD', 0.01)
Parameter('bind_RAFi_RAF_phi', 1.0)
Expression('bind_BRAFi_RAF_kr', bind_RAFi_RAF_kf*bind_RAFi_RAF_kD)
Expression('Gf_bind_BRAFi_RAF', log(bind_RAFi_RAF_kD))
Expression('Ea0_bind_BRAFi_RAF', -(bind_RAFi_RAF_phi*log(bind_RAFi_RAF_kD) +
log(bind_RAFi_RAF_kf)))
Expression('bind_CRAFi_RAF_kr', bind_RAFi_RAF_kf*bind_RAFi_RAF_kD)
Expression('Gf_bind_CRAFi_RAF', log(bind_RAFi_RAF_kD))
Expression('Ea0_bind_CRAFi_RAF', -(bind_RAFi_RAF_phi*log(bind_RAFi_RAF_kD) +
log(bind_RAFi_RAF_kf)))
Rule('RAFi_and_BRAF_bind_and_dissociate',
     RAFi(raf=None) + BRAF(rafi=None) | RAFi(raf=1) % BRAF(rafi=1),
     bind_RAFi_RAF_phi,
     Ea0_bind_BRAFi_RAF,
     energy=True)
Rule('RAFi_and_CRAF_bind_and_dissociate',
     RAFi(raf=None) + CRAF(rafi=None) | RAFi(raf=1) % CRAF(rafi=1),
     bind_RAFi_RAF_phi,
     Ea0_bind_CRAFi_RAF,
     energy=True)
EnergyPattern('ep_bind_BRAFi_RAF', RAFi(raf=1) % BRAF(rafi=1), Gf_bind_BRAFi_
RAF)
```

```
EnergyPattern('ep_bind_CRAFi_RAF', RAFi(raf=1) % CRAF(rafi=1), Gf_bind_CRAFi_
RAF)

# RAFi mediated affinity modulation
Parameter('ep_RAF_RAF_mod_RAFi_single_deltaG', 0.001)
Parameter('ep_RAF_RAF_mod_RAFi_double_deltaG', 1000.0)
Expression('ep_RAF_RAF_mod_RAFi_single_Gf', log(ep_RAF_RAF_mod_RAFi_single_de
ltaG))
Expression('ep_RAF_RAF_mod_RAFi_double_Gf',
           log(ep_RAF_RAF_mod_RAFi_double_deltaG) + log(ep_RAF_RAF_mod_RAFi_s
ingle_deltaG))

## Single Vemurafenib bound Raf dimers
EnergyPattern('ep_BRAF_BRAF_mod_RAFi_single',
              BRAF(raf=1, rafi=None) % BRAF(raf=1, rafi=2) % RAFi(raf=2),
              ep_RAF_RAF_mod_RAFi_single_Gf)
EnergyPattern('ep_BRAF_CRAF_mod_RAFi_single',
              BRAF(raf=1, rafi=None) % CRAF(raf=1, rafi=2) % RAFi(raf=2),
              ep_RAF_RAF_mod_RAFi_single_Gf)
EnergyPattern('ep_CRAF_BRAF_mod_RAFi_single',
              CRAF(raf=1, rafi=None) % BRAF(raf=1, rafi=2) % RAFi(raf=2),
              ep_RAF_RAF_mod_RAFi_single_Gf)
EnergyPattern('ep_CRAF_CRAF_mod_RAFi_single',
              CRAF(raf=1, rafi=None) % CRAF(raf=1, rafi=2) % RAFi(raf=2),
              ep_RAF_RAF_mod_RAFi_single_Gf)

## Double Vemurafenib bound Raf dimers
EnergyPattern('ep_BRAF_BRAF_mod_RAFi_double',
              RAFi(raf=2) % BRAF(raf=1, rafi=2) % BRAF(raf=1, rafi=3) % RAFi(
raf=3),
              ep_RAF_RAF_mod_RAFi_double_Gf)
EnergyPattern('ep_BRAF_CRAF_mod_RAFi_double',
              RAFi(raf=2) % BRAF(raf=1, rafi=2) % CRAF(raf=1, rafi=3) % RAFi(
raf=3),
              ep_RAF_RAF_mod_RAFi_double_Gf)
EnergyPattern('ep_CRAF_CRAF_mod_RAFi_double',
              RAFi(raf=2) % CRAF(raf=1, rafi=2) % CRAF(raf=1, rafi=3) % RAFi(
raf=3),
              ep_RAF_RAF_mod_RAFi_double_Gf);
```

#### Mek activation

Raf activation of Mek is mediated through phosphorylation at to S218 and S222 . As neither mass-spectrometry nor immunofluorescence measurements resolved individual phosphorylations on these sites, we decided to combine both into a single `phospho` site in the model. We implement the respective Mek phosphorylation as two step catalytic process including binding and phosphorylation. In agreement with previous studies, we assume that binding of Raf to Mek does not require prior activation of Raf, but is specific for unphosphorylated Mek .

```
# Raf-Mek binding
Parameter('bind_RAF_MEKphosphou_kf', 10.0)
Parameter('bind_RAF_MEKphosphou_kD', 0.01)
Parameter('bind_RAF_MEKphosphou_phi', 1.0)
Expression('bind_BRAF_MEKphosphou_kr', bind_RAF_MEKphosphou_kf*bind_RAF_MEKph
osphou_kD)
Expression('Gf_bind_BRAF_MEKphosphou', log(bind_RAF_MEKphosphou_kD))
Expression('Ea0_bind_BRAF_MEKphosphou',
           -(bind_RAF_MEKphosphou_phi*log(bind_RAF_MEKphosphou_kD) + log(bind
_RAF_MEKphosphou_kf)))
Expression('bind_CRAF_MEKphosphou_kr', bind_RAF_MEKphosphou_kf*bind_RAF_MEKph
osphou_kD)
Expression('Gf_bind_CRAF_MEKphosphou', log(bind_RAF_MEKphosphou_kD))
Expression('Ea0_bind_CRAF_MEKphosphou',
           -(bind_RAF_MEKphosphou_phi*log(bind_RAF_MEKphosphou_kD) + log(bind
_RAF_MEKphosphou_kf)))
Rule('BRAF_and_uMEK_bind_and_dissociate',
     BRAF(mek=None) + MEK(phospho='u', raf=None) | BRAF(mek=1) % MEK(phospho=
'u', raf=1),
     bind_RAF_MEKphosphou_phi,
     Ea0_bind_BRAF_MEKphosphou,
     energy=True)
Rule('CRAF_and_uMEK_bind_and_dissociate',
     CRAF(mek=None) + MEK(phospho='u', raf=None) | CRAF(mek=1) % MEK(phospho=
'u', raf=1),
     bind_RAF_MEKphosphou_phi,
     Ea0_bind_CRAF_MEKphosphou,
     energy=True)
EnergyPattern('ep_bind_BRAF_MEKphosphou', BRAF(mek=1) % MEK(phospho='u', raf=
1), Gf_bind_BRAF_MEKphosphou)
EnergyPattern('ep_bind_CRAF_MEKphosphou', CRAF(mek=1) % MEK(phospho='u', raf=
1), Gf_bind_CRAF_MEKphosphou);
```

For the activation step, we implement a physiological activation through Raf-Ras tetramers, which is implemented in four rules that account for all possible Raf homo- and heterodimer configurations, as well as oncogenic activation through B-Raf V600E mutation, which is implemented in four rules that account for B-Raf V600E molecules complexes that do not signal as dimers. In both cases, we require that the activating Raf molecule is not bound by an inhibitor and assume that kinetic rates are the same in both cases.

Experimental data presented in this study indicated phospho-Mek accumulation for physiological signaling but not for oncogenic signaling, which is consistent with previous studies . we required Mek not to be bound by an inhibitor. The exact molecular mechanisms for this difference between dimer mediated and V600E mediated are unknown, yet multiple studies point towards KSR, a scaffolding protein which was not included in the model, as a possible mediator of this difference .

```
# phosphorylation
Parameter('catalyze_RAFrafiNone_MEK_p_kcat', 10.0)
## Raf-dimer mediated
Rule('BRAF_BRAF_phosphorylates_MEK',
     MEK(phospho='u', raf=1) % BRAF(RBD=ANY, mek=1, raf=2, rafi=None) % BRAF(
RBD=ANY, raf=2)
     >>
     MEK(phospho='p', raf=None) + BRAF(RBD=ANY, mek=None, raf=2, rafi=None) %
BRAF(RBD=ANY, raf=2),
     catalyze_RAFrafiNone_MEK_p_kcat)
Rule('BRAF_CRAF_phosphorylates_MEK',
     MEK(phospho='u', raf=1) % BRAF(RBD=ANY, mek=1, raf=2, rafi=None) % CRAF(
RBD=ANY, raf=2)
     >>
     MEK(phospho='p', raf=None) + BRAF(RBD=ANY, mek=None, raf=2, rafi=None) %
CRAF(RBD=ANY, raf=2),
     catalyze_RAFrafiNone_MEK_p_kcat)
Rule('CRAF_BRAF_phosphorylates_MEK',
     MEK(phospho='u', raf=1) % CRAF(RBD=ANY, mek=1, raf=2, rafi=None) % BRAF(
RBD=ANY, raf=2)
     >>
     MEK(phospho='p', raf=None) + CRAF(RBD=ANY, mek=None, raf=2, rafi=None) %
BRAF(RBD=ANY, raf=2),
     catalyze_RAFrafiNone_MEK_p_kcat)
Rule('CRAF_CRAF_phosphorylates_MEK',
     MEK(phospho='u', raf=1) % CRAF(RBD=ANY, mek=1, raf=2, rafi=None) % CRAF(
RBD=ANY, raf=2)
     >>
     MEK(phospho='p', raf=None) + CRAF(RBD=ANY, mek=None, raf=2, rafi=None) %
CRAF(RBD=ANY, raf=2),
     catalyze_RAFrafiNone_MEK_p_kcat)
## B-Raf V600E mediated
Rule('BRAFV600E_phosphorylates_MEK_1',
     MEK(meki=None, phospho='u', raf=1) % BRAF(AA600='E', mek=1, raf=None, ra
fi=None)
     >>
     MEK(meki=None, phospho='p', raf=None) + BRAF(AA600='E', mek=None, raf=No
ne, rafi=None),
     catalyze_RAFrafiNone_MEK_p_kcat)
Rule('BRAFV600E_phosphorylates_MEK_2',
```

```
    MEK(meki=None, phospho='u', raf=1) % BRAF(AA600='E', RBD=None, mek=1, ra
f=ANY, rafi=None)
    >>
    MEK(meki=None, phospho='p', raf=None) + BRAF(AA600='E', RBD=None, mek=No
ne, raf=ANY, rafi=None),
    catalyze_RAFrafiNone_MEK_p_kcat)
Rule('BRAFV600E_phosphorylates_MEK_3',
    MEK(meki=None, phospho='u', raf=1) % BRAF(AA600='E', RBD=ANY, mek=1, raf
=2, rafi=None) % BRAF(RBD=None, raf=2)
    >>
    MEK(meki=None, phospho='p', raf=None) + BRAF(AA600='E', RBD=ANY, mek=Non
e, raf=2, rafi=None) % BRAF(RBD=None, raf=2),
    catalyze_RAFrafiNone_MEK_p_kcat)
Rule('BRAFV600E_phosphorylates_MEK_4',
    MEK(meki=None, phospho='u', raf=1) % BRAF(AA600='E', RBD=ANY, mek=1, raf
=2, rafi=None) % CRAF(RBD=None, raf=2)
    >>
    MEK(meki=None, phospho='p', raf=None) + BRAF(AA600='E', RBD=ANY, mek=Non
e, raf=2, rafi=None) % CRAF(RBD=None, raf=2),
    catalyze_RAFrafiNone_MEK_p_kcat);
```

PP2A and PPP1 have been suggested as putative phosphatases of Mek1, but exact regulatory mechanisms remain poorly understood. Accordingly we implement MEK dephosphorylation as conversion rule with constant rate.

```
# dephosphorylation
Parameter('catalyze_PP2A_MEK_u_kcatr', 1.0)
Expression('catalyze_PP2A_MEK_u_kcat', catalyze_RAFrafiNone_MEK_p_kcat*cataly
ze_PP2A_MEK_u_kcatr)
Rule('MEK_is_dephosphorylated', MEK(phospho='p') >> MEK(phospho='u'), catalyz
e_PP2A_MEK_u_kcat);
```

#### Mek inhibition

Cobimetinib is an allosteric (type III) inhibitor that potently and selectively inhibits Erk activation when binding Mek. Recent studies reported a distinct potency of allosteric Mek inhibitor for Ras-GTP versus B-Raf V600E driven Mek signaling. One component of this difference in potency is the accumulation of phosphorylated Mek in Ras-GTP driven signaling, but not in B-Raf V600E driven signaling that was already described in the previous Section. A second component is the differential affinity of allosteric Mek inihibitors towards phosphorylated MEK . Accordingly, we implement binding as energy rule and add an energy pattern that allows Mek phosphorylation to modulation the respective affinity.

```
# MEKi binding
Parameter('bind_MEKi_MEK_kf', 10.0)
Parameter('bind_MEKi_MEK_kD', 0.01)
Parameter('bind_MEKi_MEK_phi', 1.0)
```

```
Expression('bind_MEKi_MEK_kr', bind_MEKi_MEK_kf*bind_MEKi_MEK_kD)
Expression('Gf_bind_MEKi_MEK', log(bind_MEKi_MEK_kD))
Expression('Ea0_bind_MEKi_MEK', -(bind_MEKi_MEK_phi*log(bind_MEKi_MEK_kD) + l
og(bind_MEKi_MEK_kf)))
Rule('MEKi_and_MEK_bind_and_dissociate',
     MEKi(mek=None) + MEK(meki=None) | MEKi(mek=1) % MEK(meki=1),
     bind_MEKi_MEK_phi,
     Ea0_bind_MEKi_MEK,
     energy=True)
EnergyPattern('ep_bind_MEKi_MEK', MEKi(mek=1) % MEK(meki=1), Gf_bind_MEKi_MEK
);

# Modulated pMEK affinity
Parameter('ep_MEKphosphop_MEKi_deltaG', 100.0)
Expression('ep_MEKphosphop_MEKi_Gf', log(ep_MEKphosphop_MEKi_deltaG))
EnergyPattern('ep_MEKphosphop_MEKi_single', MEK(meki=1, phospho='p') % MEKi(m
ek=1), ep_MEKphosphop_MEKi_Gf);
```

#### Erk activation

Mek phosphorylates Erk1 at T202 and T185 and Erk2 at Y204 and Y187. As for Mek phosphorylation sites, neither mass-spectrometry nor immunofluorescence measurements resolved individual phosphorylations, both sites were combined into a single `phospho` site in the model.

As for the Mek phosphorylation we implement Erk phosphorylation as two step catalysis reaction. The binding between Mek and Erk is mediated through interaction of the MAPK-Docking site on the N-Terminus of MEK (`Dsite` site in the model) with the common docking domain on Erk (`CD` site in the model) . In combination with Mek's nuclear export sequence, this interaction allows Mek to act as a cytosolic anchor for inactive Erk , which suggests that this interaction is not dependent of Mek phosphorylation. After Mek activation, Erk releases from Mek and shuttles in and out of the nucleus, which suggests that affinity between Mek and Erk depends on Erk phosphorylation . In the model, we implement the dependence on Erk phosphorylation as binary interaction, where Mek only binds unphosphorylated Erk. Nuclear shuttling of Mek and Erk was not implemented in the model.

```
# Mek-Erk binding
Parameter('bind_MEK_ERKphosphou_kf', 10.0)
Parameter('bind_MEK_ERKphosphou_kD', 0.01)
Expression('bind_MEK_ERKphosphou_kr', bind_MEK_ERKphosphou_kf*bind_MEK_ERKpho
sphou_kD)
Rule('MEK_binds_uERK',
     MEK(Dsite=None) + ERK(CD=None, phospho='u') >> MEK(Dsite=1) % ERK(CD=1,
phospho='u'),
     bind_MEK_ERKphosphou_kf)
Rule('MEK_dissociates_from_ERK',
     MEK(Dsite=1) % ERK(CD=1) >> MEK(Dsite=None) + ERK(CD=None),
     bind_MEK_ERKphosphou_kr)
```

```
# Erk phosphorylation
Parameter('catalyze_MEKmekiNone_phosphop_ERK_p_kcat', 10.0)
Rule('pMEK_phosphorylates_ERK',
     ERK(CD=1, phospho='u') % MEK(Dsite=1, meki=None, phospho='p')
     >>
     ERK(CD=None, phospho='p') + MEK(Dsite=None, meki=None, phospho='p'),
     catalyze_MEKmekiNone_phosphop_ERK_p_kcat);
```

### Feedback mechanisms

Activation of Erk through phosphorylation induces a variety of different negative feedback mechanisms that inhibit signal transduction in Egfr and Erk pathways . In this module we specifically implement four of these mechanisms that, specifically the increase in Spry, Egfr and Dusp expression levels as well as phosphorylation of Sos1. For this purpose we introduce two new monomers, namely Dusp (which describes Dusp4 and Dusp6) and Spry (which describes Spry2 and Spry4).

Some of the negative feedback mechanisms that were described in previous studies are omitted in this model, including Erk negative feedback phosphorylations on Mek, and Raf. As the timescale of phosphorylation reactions is in the order of seconds, a high inhibitory potency of these feedback mechanisms should give rise to phospho-Erk pulses on a timescale of seconds. Yet the experimentally observed timescale of phospho-Erk pulses was in the order of minutes, which suggests that it is unlikely that phosphorylations feedbacks are the primary determinant of the pulse shape. This hypothesis is backed by previous studies on phospho-turnover in Egfr mediated signaling . Accordingly we only expect a subtle effect of the negative feedback mechanisms, which would be difficult to be resolve in the model, given that no experimental data on respective phosphorylation sites or Ras-GTP levels were available.

```
Monomer('DUSP', ['erk'])
Annotation(DUSP, 'http://identifiers.org/uniprot/Q13115', 'is')
Annotation(DUSP, 'http://identifiers.org/uniprot/Q16828', 'is')

Monomer('SPRY', ['SH3'])
Annotation(SPRY, 'http://identifiers.org/uniprot/O43597', 'is')
Annotation(SPRY, 'http://identifiers.org/uniprot/Q9C004', 'is');
```

#### Spry expression

The proteomic data collected in this study confirmed previous findings that phosphorylation of Erk enhances expression of Spry. Increase in Spry expression inhibits Egfr signaling and is mediated by regulation of downstream transcription factors . Specifically, Spry competes with Sos1 for the binding of Grb2 . Accordingly higher levels of Spry can antagonize Sos1 signaling by sequestering Grb2 in Spry:Grb2 complexes.

Here, we implement basal synthesis and degradation reactions for Spry, which are parameterised with `[...]_eq` and `[...]_kdeg` parameters as done for Egfr expression in the Section on Grb2 activation. On top of this basal expression, we implement an additional expression rule for which the expression rate linearly depends on the abundance of phospho-Erk. To avoid correlations of parameter estimates, we normalize this expression rate with the basal Spry degradation rate. The binding between Spry and Grb2 is parametrized using `[...]_kD` and `[...]_kf` parameters as previously described. Competitive binding between Spry and Sos1 for Grb2 is implemented by specifying the same binding site on Grb2 (`SH3`) for the binding of both reactions.

```
# basal expression and degradation
Parameter('SPRY_eq', 10000.0)
Parameter('synthesize_ERKphosphop_SPRY_kdeg', 10.0)
Expression('synthesize_ERKphosphop_SPRY_ksyn',
           1000000.0*SPRY_eq*(N_Avogadro*volume)**(-1)*synthesize_ERKphosphop
_SPRY_kdeg)
Rule('basal_synthesis_SPRY',
     None >> SPRY(SH3=None),
     synthesize_ERKphosphop_SPRY_ksyn)
Rule('basal_degradation_SPRY',
     SPRY() >> None,
     synthesize_ERKphosphop_SPRY_kdeg, delete_molecules=True)

# Erk mediated expression
Parameter('synthesize_ERKphosphop_SPRY_ERK_gexpslope', 1000.0)
Expression('synthesize_ERKphosphop_SPRY_kmodslope',
           synthesize_ERKphosphop_SPRY_ERK_gexpslope*synthesize_ERKphosphop_S
PRY_kdeg)
Rule('ERK_synthesizes_SPRY',
     None + ERK(phospho='p') >> SPRY(SH3=None) + ERK(phospho='p'),
     synthesize_ERKphosphop_SPRY_kmodslope)

# Spry-Grb2 binding
Parameter('bind_SPRY_GRB2_kf', 10.0)
Parameter('bind_SPRY_GRB2_kD', 0.01)
Expression('bind_SPRY_GRB2_kr', bind_SPRY_GRB2_kf*bind_SPRY_GRB2_kD)
Rule('SPRY_binds_GRB2',
     SPRY(SH3=None) + GRB2(SH3=None) >> SPRY(SH3=1) % GRB2(SH3=1),
     bind_SPRY_GRB2_kf)
Rule('SPRY_dissociates_from_GRB2',
     SPRY(SH3=1) % GRB2(SH3=1) >> SPRY(SH3=None) + GRB2(SH3=None),
     bind_SPRY_GRB2_kr);
```

#### Sos1 phosphorylation

Phospho-proteomic data collected in this study confirmed previous findings that phosphorylation of Erk enhances phosphorylation of Sos1 at S1134. Phosphorylation of Sos1 inhibits Egfr signaling and is mediated through upregulation of Rsk expression and phosphorylation . Specifically it creates a docking site for 14-3-3 on Sos1 and thereby sequestering it from Grb2 .

We do not include Rsk or 14-3-3 in the model as absolute abundances were measured for neither of proteins. Instead we decided to implement basal phosphorylation and dephosphorylation rules as well as an phospho-Erk dependent phosphorylation rule for which the phosphorylation linearly depends on Erk phosphorylation. The baseline rules were implemented using constant rate `[...]_kbase` and `[...]_kcat` respectively. To avoid parameter correlations, the phospho-Erk dependent phosphorylation rate was normalized by the baseline phosphorylation rate.

```
# base phosphorylation rate
Parameter('catalyze_ERKphosphop_SOS1_pS1134_kbase', 0.0)
Parameter('catalyze_ERKphosphop_SOS1_pS1134_kcat', 10.0)
Parameter('catalyze_phosphatase_SOS1_uS1134_kcatr', 1.0)
Expression('catalyze_phosphatase_SOS1_uS1134_kcat',
           catalyze_ERKphosphop_SOS1_pS1134_kcat*catalyze_phosphatase_SOS1_uS
1134_kcatr)
Rule('SOS1_is_phosphorylated',
     SOS1(S1134='u') >> SOS1(S1134='p'),
     catalyze_ERKphosphop_SOS1_pS1134_kbase)
Rule('SOS1_is_dephosphorylated',
     SOS1(S1134='p') >> SOS1(S1134='u'),
     catalyze_phosphatase_SOS1_uS1134_kcat)
Rule('pERK_phosphorylates_SOS1',
     ERK(phospho='p') + SOS1(S1134='u') >> ERK(phospho='p') + SOS1(S1134='p')
,
     catalyze_ERKphosphop_SOS1_pS1134_kcat);
```

As the binding between as Grb2 and Sos1 was not implemented as energy rule, we cannot use an energy pattern to implement the modulated affinity of Grb2 to Sos1. Instead, we implement an additional unbinding reaction for Sos1 that is phosphorylation dependent and mimics the effect of an energy pattern.

```
Parameter('ep_SOS1S1134p_GRB2_deltaG', 2.0)
Expression('__reverse_bind_GRB2_SOS1_local1', bind_GRB2_SOS1_kD*bind_GRB2_SOS
1_kf*ep_SOS1S1134p_GRB2_deltaG)
Rule('pSOS1_has_modulated_affinity_to_GRB2',
     SOS1(S1134='p', SH3=1) % GRB2(SH3=1) >> SOS1(S1134='p', SH3=None) + GRB2
(SH3=None),
     __reverse_bind_GRB2_SOS1_local1);
```

#### Egfr expression

Proteomic data collected in this study confirmed previous findings that phosphorylation of Erk increases Egfr expression. Increased Egfr expression promotes Egfr signaling and is mediated through regulation of downstream transcription factors .

As baseline Egfr expression and degradation rules are already implemented in the Section on GRB2 activation, we here simply add an expression rule for which the rate linearly depends on phospho-

Erk. As for Spry expression, we normalized the phospho-Erk dependent expression rate by the baseline degradation rate.

```
Parameter('synthesize_ERKphosphop_EGFR_ERK_gexpslope', 1000.0)
Expression('synthesize_ERKphosphop_EGFR_kmodslope',
           synthesize_ERKphosphop_EGFR_ERK_gexpslope*synthesize_ERKphosphop_E
GFR_kdeg)
Rule('ERK_synthesizes_EGFR',
     None + ERK(phospho='p') >> EGFR(SH2=None, rtk=None, rtkf=None) + ERK(pho
spho='p'),
     synthesize_ERKphosphop_EGFR_kmodslope);
```

#### Dusp expression

The proteomic data collected in this study confirmed previous findings that phosphorylation of Erk enhances expression of Dusp. Increase in Dusp expression inhibits Erk signaling and is mediated by regulation of downstream transcription factors . Specifically, Dusp are phosphatases that dephosphorylated Erk on the same residues that are phosphorylated by Mek. In the proteomic data we individually measured Dusp4 and Dusp6 abundances. Dusp4 has primarily nuclear localisation , while Dusp6 has primarily cytosolic localisation . As we did not implement shuttling of Erk between the nucleus and cytosol, we lumped Dusp4 and Dusp6 species.

We implement the dephosphorylation as two-step catalytic process. As both Dusps interact with Erk via the common docking domain that also mediates interaction between Mek to Erk , the binding was implemented using the same site on the Erk (`CD`) monomer. This could lead to competitive binding between Mek and Erk, yet as Dusp, in contrast to Mek, seem to preferentially bind the phosphorylated form of Erk , we constrained binding of Dusp to Erk on Erk phosphorylation, which makes the two binding reactions independent.

```
# expression
Parameter('DUSP_eq', 10000.0)
Parameter('synthesize_ERKphosphop_DUSP_ERK_gexpslope', 1000.0)
Parameter('synthesize_ERKphosphop_DUSP_kdeg', 10.0)
Expression('synthesize_ERKphosphop_DUSP_ksyn',
           1000000.0*DUSP_eq*(N_Avogadro*volume)**(-1)*synthesize_ERKphosphop
_DUSP_kdeg)
Expression('synthesize_ERKphosphop_DUSP_kmodslope',
           synthesize_ERKphosphop_DUSP_ERK_gexpslope*synthesize_ERKphosphop_D
USP_kdeg)
Rule('basal_synthesis_DUSP',
     None >> DUSP(erk=None),
     synthesize_ERKphosphop_DUSP_ksyn)
Rule('basal_degradation_DUSP',
     DUSP() >> None,
     synthesize_ERKphosphop_DUSP_kdeg,
```

```
     delete_molecules=True)
Rule('ERK_synthesizes_DUSP',
     None + ERK(phospho='p') >> DUSP(erk=None) + ERK(phospho='p'),
     synthesize_ERKphosphop_DUSP_kmodslope)

# Dusp-Erk binding
Parameter('bind_DUSP_ERKphosphop_kf', 10.0)
Parameter('bind_DUSP_ERKphosphop_kD', 0.01)
Expression('bind_DUSP_ERKphosphop_kr', bind_DUSP_ERKphosphop_kf*bind_DUSP_ERK
phosphop_kD)
Rule('DUSP_binds_pERK',
     DUSP(erk=None) + ERK(CD=None, phospho='p') >> DUSP(erk=1) % ERK(CD=1, ph
ospho='p'),
     bind_DUSP_ERKphosphop_kf)
Rule('DUSP_dissociates_from_ERK',
     DUSP(erk=1) % ERK(CD=1) >> DUSP(erk=None) + ERK(CD=None),
     bind_DUSP_ERKphosphop_kr)

# Erk dephosphorylation
Parameter('catalyze_DUSP_ERK_u_kcatr', 1.0)
Expression('catalyze_DUSP_ERK_u_kcat', catalyze_MEKmekiNone_phosphop_ERK_p_kc
at*catalyze_DUSP_ERK_u_kcatr)
Rule('DUSP_dephosphorylates_ERK',
     ERK(CD=1, phospho='p') % DUSP(erk=1) >> ERK(CD=None, phospho='u') + DUSP
(erk=None),
     catalyze_DUSP_ERK_u_kcat);
```

#### *Initialization*

For all monomers for which we did not implement expression and basal degradation rules we implemented initial abundances that were assumed to be constant in all experiments. To simplify specification of estimation boundaries for these parameters, these parameters were assumed to be specified in [molecules/cell] and then accordingly transformed to yield model initializations in [uM].

```
Parameter('BRAF_0', 0.0)
Parameter('CRAF_0', 0.0)
Parameter('RAS_0', 0.0)
Parameter('MEK_0', 0.0)
Parameter('ERK_0', 0.0)
Parameter('GRB2_0', 0.0)
Parameter('SOS1_0', 0.0)

Expression('initBRAF', 1000000.0*BRAF_0*(N_Avogadro*volume)**(-1))
Expression('initCRAF', 1000000.0*CRAF_0*(N_Avogadro*volume)**(-1))
Expression('initRAS', 1000000.0*RAS_0*(N_Avogadro*volume)**(-1))
Expression('initMEK', 1000000.0*MEK_0*(N_Avogadro*volume)**(-1))
```

```
Expression('initERK', 1000000.0*ERK_0*(N_Avogadro*volume)**(-1))
Expression('initGRB2', 1000000.0*GRB2_0*(N_Avogadro*volume)**(-1))
Expression('initSOS1', 1000000.0*SOS1_0*(N_Avogadro*volume)**(-1))

Initial(BRAF(AA600='E', RBD=None, mek=None, raf=None, rafi=None), initBRAF)
Initial(CRAF(RBD=None, mek=None, raf=None, rafi=None), initCRAF)
Initial(RAS(raf=None, sos1=None, state='gdp'), initRAS)
Initial(MEK(Dsite=None, meki=None, phospho='u', raf=None), initMEK)
Initial(ERK(CD=None, phospho='u'), initERK)
Initial(GRB2(SH2=None, SH3=None), initGRB2)
Initial(SOS1(S1134='u', SH3=None, ras=None), initSOS1);
```

#### *Link to experimental data*

To calibrate the model to experimental data we create a series of expression that account for normalizations in data generation. For absolute proteomic measurements we create expressions `t[...]_obs` that convert the unit of model simulations from [uM] to [molecules/cell]. For phospho-proteomic measurements we create expressions `p[...]_obs` that are equal to phospho-abundances normalized by total abundances. For immuno-fluorescence measurements we created expressions `p[...]_IF_obs` that were first normalized by total abundances and then scaled by a mulitplicative factor `[...]_IF_scale` and an additive factor `[...]_IF_offset`. For species that describe multiple protein isoforms, absolute proteomic data was averaged over both isoforms.

```
# proteomic measurements
Observable('tBRAF', BRAF())
Observable('tBRAF600E', BRAF(AA600='E'))
Observable('tCRAF', CRAF())
Observable('tRAS', RAS())
Observable('tRAFi', RAFi())
Observable('tMEK', MEK())
Observable('tMEKi', MEKi())
Observable('tERK', ERK())
Observable('tDUSP', DUSP())
Observable('tEGFR', EGFR())
Observable('tEGF', EGF())
Observable('tGRB2', GRB2())
Observable('tSPRY', SPRY())
Observable('tSOS1', SOS1())
Expression('tBRAF_obs', 1.0e-6*tBRAF*N_Avogadro*volume)
Expression('tCRAF_obs', 1.0e-6*tCRAF*N_Avogadro*volume)
Expression('tRAS_obs', 1.0e-6*tRAS*N_Avogadro*volume)
Expression('tMEK_obs', 1.0e-6*tMEK*N_Avogadro*volume)
Expression('tDUSP_obs', 1.0e-6*tDUSP*N_Avogadro*volume)
```

```
Expression('tEGFR_obs', 1.0e-6*tEGFR*N_Avogadro*volume)
Expression('tEGF_obs', 1.0e-6*tEGF*N_Avogadro*volume)
Expression('tGRB2_obs', 1.0e-6*tGRB2*N_Avogadro*volume)
Expression('tSPRY_obs', 1.0e-6*tSPRY*N_Avogadro*volume)
Expression('tSOS1_obs', 1.0e-6*tSOS1*N_Avogadro*volume)
Expression('tERK_obs', 1.0e-6*tERK*N_Avogadro*volume)

# phospho-proteomic measurements
Observable('pERK', ERK(phospho='p'))
Observable('pS1134SOS1', SOS1(S1134='p'))
Expression('pERK_obs', pERK*tERK**(-1))
Expression('pS1134SOS1_obs', pS1134SOS1*tSOS1**(-1))

# immunofluorescence measurements
Parameter('pMEK_IF_scale', 1.0)
Parameter('pMEK_IF_offset', 0.1)
Parameter('pERK_IF_scale', 1.0)
Parameter('pERK_IF_offset', 0.1)
Observable('pMEK', MEK(phospho='p'))
Expression('pMEK_obs', pMEK*tMEK**(-1))
Expression('pERK_IF_obs', pERK_IF_offset + pERK_IF_scale*pERK_obs)
Expression('pMEK_IF_obs', pMEK_IF_offset + pMEK_IF_scale*pMEK_obs);
```

#### *Observables*

We add observables that are used to extract the biological activity of proteins of interest:

```
#active SOS1
Observable('SOS1_active', GRB2(SH2=ANY, SH3=1) % SOS1(SH3=1))
#unphosphorylated SOS1
Observable('uS1134SOS1', SOS1(S1134='u'))
#Ras-GDP and -GTP
Observable('RAS_gtp', RAS(state='gtp'))
Observable('RAS_gdp', RAS(state='gdp'))
#active RAF dimers
Observable('active_RAF_dimers', BRAF(RBD=ANY, raf=1, rafi=None) % BRAF(RBD=AN
Y, raf=1) + BRAF(RBD=ANY, raf=1, rafi=None) % CRAF(RBD=ANY, raf=1) + CRAF(RBD
=ANY, raf=1, rafi=None) % BRAF(RBD=ANY, raf=1) + CRAF(RBD=ANY, raf=1, rafi=No
ne) % CRAF(RBD=ANY, raf=1)),
#active RAF monomers
Observable('active_RAF_monomers', BRAF(AA600='E', raf=None, rafi=None)),
#active and unphosphorylated MEK
Observable('active_MEK', MEK(meki=None, phospho='p'));
```

```
Observable('uMEK', MEK(phospho='u'))
#unophosphorylated ERK
Observable('uERK', ERK(phospho='u'));
```
