## Supplementary material for "Sporadic ERK pulses drive non-genetic resistance in drug-adapted BRAF^V600E^ melanoma cells": Key Resource Table

**KEY RESOURCES TABLE**

| REAGENT or RESOURCE | SOURCE | IDENTIFIER |
|---|---|---|
| Antibodies | | |
| Phospho-p44/42 MAPK (Erk1/2) (Thr202/Tyr204) antibody | Cell Signaling Technology | Cat# 4370, RRID:AB_2315112 |
| Phospho-MEK1/2 (Ser217/221) antibody | Cell Signaling Technology | Cat# 9121, RRID:AB_331648 |
| Phospho-p90RSK (Thr359) (D1E9) antibody | Cell Signaling Technology | Cat# 8753, RRID:AB_2783561) |
| Phospho-S6 Ribosomal Protein (Ser240/244) (D68F8) XP Rabbit mAb (Alexa Fluor 488 Conjugate) antibody | Cell Signaling Technology | Cat# 5018, RRID:AB_10695861 |
| Phospho-S6 Ribosomal Protein (Ser235/236) (D57.2.2E) XP Rabbit mAb (Alexa Fluor 555 Conjugate) antibody | Cell Signaling Technology | Cat# 3985, RRID:AB_10693792 |
| EGFR Monoclonal Antibody (199.12) | Thermo Fisher Scientific | Cat# MA5-13319, RRID:AB_10985841 |
| Alexa Fluor 647, Donkey anti-Rabbit IgG (H+L) Secondary Antibody | Thermo Fisher Scientific | Cat# A-31573 |
| Alexa Fluor 488, Donkey anti-Mouse IgG (H+L) Secondary Antibody | Thermo Fisher Scientific | Cat# A21202 |
| EGF Receptor (D38B1) XP® Rabbit mAb | Cell Signaling Technology | Cat# 4267 |
| β-Actin Antibody (C4) | Santa Cruz Biotechnology | Cat# sc-47778 |
| Anti-rabbit IgG, HRP-linked Antibody | Cell Signaling Technology | Cat# 7074 |
| Anti-mouse IgG, HRP-linked Antibody | Cell Signaling Technology | Cat# 7076 |
| Biological Samples | | |

| | | |
|---|---|---|
| A375 Xenograft Formalin-Fixed Paraffin-Embedded (FFPE) tissue slides | Fallahi-Sichani et al., 2017 | N\A |
| Chemicals, Peptides, and Recombinant Proteins | | |
| Vemurafenib, RAF inhibitor | MedChem Express | Cat# HY-12057 |
| Dabrafenib, RAF inhibitor | MedChem Express | Cat# HY-14660 |
| PLX8394, RAF inhibitor | MedChem Express | Cat# HY-18972 |
| LY3009120, RAF inhibitor | MedChem Express | Cat# HY-12558 |
| AZ628, RAF inhibitor | MedChem Express | Cat# HY-11004 |
| Cobimetinib, MEK inhibitor | MedChem Express | Cat# HY-13064 |
| Trametinib, MEK inhibitor | MedChem Express | Cat# HY-10999 |
| Selumetinib, MEK inhibitor | MedChem Express | Cat# HY-50706 |
| Binimetinib, MEK inhibitor | MedChem Express | Cat# HY-15202 |
| PD0325901, MEK inhibitor | MedChem Express | Cat# HY-10254 |
| Lapatinib, ERBB inhibitor | MedChem Express | Cat# HY-50898 |
| Erlotinib, ERBB inhibitor | MedChem Express | Cat# HY-50896 |
| SHP099, SHP2 inhibitor | MedChem Express | Cat# HY-100388 |
| Recombinant Human EGF, growth factor | Peprotech | Cat# 100-15 |
| Recombinant Human Heregulinβ-1, growth factor | Peprotech | Cat# 100-03 |
| Recombinant Human FGF-8a, growth factor | Peprotech | Cat# 100-25A |
| Recombinant Human HGF, growth factor | Peprotech | Cat#100-39 |
| EdU (5-ethynyl-2'-deoxyuridine) | Lumiprobe, Hunt Valley, MD | Cat# 10540 |
| Critical Commercial Assays | | |
| Deposited Data | | |

| | | |
|---|---|---|
| Gene expression data (RNA seq) of A375 melanoma cells were deposited in the GEO (Gene Expression Omnibus) database: https://www.ncbi.nlm.nih.gov/geo/ | GEO (Gene Expression Omnibus) | Accession number: GSE127988<br><br>Secure token for anonymous review: wvclkgmqddwpbqz |
| Protein abundance and phosphorylation data (SRM proteomics) were deposited in the Panorama public database at https://panoramaweb.org/labkey/ | Panorama (Repository Software for Targeted Mass Spectrometry Assays from Skyline) | https://panoramaweb.org/fWeE7i.url<br><br>Anonymous reviewer login info:<br><br><br><br>Password: %h4JDQCY |
| Processed datasets used to generate main and supplementary figures in the manuscript, provided as .csv files | Synapse database | https://www.synapse.org/#!Synapse:syn20551877/files/<br><br>Synapse ID: syn20551877<br><br>DOI:10.7303/syn20551877.<br><br>The repository is public and can be download without registration. |
| Experimental Models: Cell Lines | | |
| Human: A-375 (A375), Melanoma Cell Line | MGH Cancer Center, primary source ATCC | ATCC Cat# CRL-1619, RRID:CVCL_0132 |
| Human: HT-29 (HT29), Colorectal Cell Line | Merrimack Pharmaceuticals | NCI-DTP Cat# HT-29, RRID:CVCL_0320 |

| | | |
|---|---|---|
| Human:C32, Melanoma Cell Line | MGH Cancer Center, primary source ATCC | ATCC Cat# CRL-1585, RRID:CVCL_1097 |
| Human: K2, Melanoma Cell Line | MGH Cancer Center, primary source ATCC | RRID:CVCL_AT85 |
| Human: MMAc-SF (MMACSF), Melanoma Cell Line | MGH Cancer Center, primary source RIKEN BioResource Center | RCB Cat# RCB1200, RRID:CVCL_1420 |
| Human: MZ-MEL-7 (MZ7MEL), Melanoma Cell Line | MGH Cancer Center, primary source Johannes Gutenberg University Mainz | RRID:CVCL_1436 |
| Human: RVH-421 (RVH421), Melanoma Cell Line | MGH Cancer Center, primary source ATCC | RRID:CVCL_1672 |
| Human: SKMEL28, Melanoma Cell Line | MGH Cancer Center, primary source ATCC | CLS Cat# 300337/p495_SK-MEL-28, RRID:CVCL_0526 |
| Human: WM115, Melanoma Cell Line | MGH Cancer Center, primary source ATCC | ATCC Cat# CRL-1675, RRID:CVCL_0040 |
| Human: HEK293T, Cell Line | ATCC | ATCC Cat# CRL-3216, RRID:CVCL_0063 |
| A375_aEGFR1 (A375 with constitutive EGFR overexpression by CRISPRa; sgRNA1) | This work | N/A |
| A375_aEGFR2 (A375 with constitutive EGFR overexpression by CRISPRa; sgRNA2) | This work | N/A |
| A375_iEGFR (A375 with constitutive EGFR knockdown by CRISPRi; sgRNA1) | This work | N/A |

| | | |
|---|---|---|
| A375 stably expressing ERK-KTR:CFP reporter | This work | N/A |
| Oligonucleotides | | |
| See Table S2 for oligonucleotide sequences | This work | N/A |
| Recombinant DNA | | |
| psPAX2 | Addgene | Cat# 12260 |
| pCMV-VSV-G | Addgene | Cat# 8454 |
| pMH0001, expresses dCas9-BFP-KRAB, for CRISPRi | Addgene | Cat# 85969 |
| pHRdSV40-dCas9-10xGCN4_v4-P2A-BFP, expresses dCas9 tagged with 10 copies of the GCN4 peptide v4 and BFP, for CRISPR a | Addgene | Cat# 60903 |
| pHRdSV40-scFv-GCN4-sfGFP-VP64-GB1-NLS, expresses an antibody that binds to the GCN4 peptide from the SunTag system, and is fused to VP64, for CRISPRa | Addgene | Cat# 60904 |
| pU6-sgRNA EF1Alpha-puro-T2A-BFP | Addgene | Cat# 60955 |
| 4_pPB_ERKKTRmTq2_H2BVenus_mCherryGemin, fluorescent reporter for H2B(Venus), ERK:TTR (mTurquoise), Geminin (mCherry) | Fallahi-Sichani et al., 2017 | N\A |
| pCMV_hyPBase | Fallahi-Sichani et al., 2017 | N\A |
| Software and Algorithms | | |
| PySB, open-source programming framework for Systems Biology modelling in Python | Lopez et al., 2013 | http://pysb.org/ |
| eBNG, energy-based modeling in BioNetGen | Sekar et al., 2017 | https://github.com/RuleWorld/bionetgen |
| Amici (Advanced Multilanguage Interface to CVODES and IDAS) framework used for parameter estimation | Fröhlich et al., 2017 | https://github.com/ICB-DCM/AMICI |
| Pypesto, a widely applicable and highly customizable toolbox for parameter estimation. | Stapor et al., 2018 | http://snakemake.googlecode.com |

| | | |
|---|---|---|
| Snakemake, workflow management system is a tool to create reproducible and scalable data analyses. | Koster and Rahmann, 2012 | https://snakemake.readthedocs.io/en/stable/ |
| Columbus, data storage and analysis system for high throughput microscopy software | PerkinElmer | http://www.perkinelmer.com/product/image-data-storage-and-analysis-system-columbus |
| Software for automatic segmentation, quantification and tracking fluorescent reporter cells live-cell imaging | Cappell SD et al., 2016 | https://github.com/scappell/Cell_tracking |
| Skyline software for analysis of proteomics samples | MacLean et al., 2010 | https://skyline.ms/project/home/software/Skyline/begin.view |
| Ashlar for stitching and registration of cyclic microscopy images of large tissue sections | Muhlich et al, in preparation | https://github.com/labsyspharm/ashlar |
| Ilastik1.3.2, is a simple, user-friendly tool for interactive image classification, segmentation and analysis. | (Sommer et al., 2011) | https://www.ilastik.org/ |
| CellProfiler3.1.8. is a free, open-source software to quantitatively measure phenotypes of images automatically. | (Lamprecht et al., 2007) | https://cellprofiler.org/ |
| histoCAT, Histology Topography Cytometry Analysis Toolbox | (Schapiro et al., 2017) | https://github.com/BodenmillerGroup/histoCAT |
| Melanoma Adaptive Resistance Model (MARM1). Provided in SBML, BNG and PySB formats. Jupyter Notebooks are provided in a Docker container for the step-by-step construction and simulation of MARM1. | This work | https://github.com/labsyspharm/marm1-supplement |
| Other | | |
| SuperSignal™ West Dura Extended Duration Substrate | Thermo Fisher Scientific | Cat# 34076 |
| HCS CellMask™ Blue Stain | Thermo Fisher Scientific | Cat# 34076 |

| | | |
|---|---|---|
| Hoechst 33342 | Invitrogen | Cat# H3570 |
